# Gaussian selection on a polygenic trait from the gene’s-eye view

**DOI:** 10.64898/2026.02.23.706325

**Authors:** Philibert Courau, Amaury Lambert, Emmanuel Schertzer

## Abstract

We study a polygenic trait under Gaussian selection at statistical equilibrium, where allelic effect, mutation rate and mutational bias are heterogeneous across loci. The model assumes a large number *L* of biallelic sites subject to reversible mutations, each allele described by its frequency in the population. Using a diffusion approximation, a mean-field approximation and neglecting linkage disequilibrium, we predict consistent phenomena across several regimes of selection: (1) a small deviation − Δ* of the trait mean from its optimal value appears and persists due to genetic mutations not aligned with selection; (2) while this deviation is often undetectable at the trait level, it leaves a substantial signature at the locus level by favoring alleles reducing it, resulting in genic selection with mean coefficient *s*\* proportional to – Δ* acting pervasively; (3) with stronger selection on the trait, (3a) the value of Δ* is decreased but the intensity of genic selection is increased in inverse proportion, resulting in an essentially constant, non negligible value of *s*\*. We show how the stationary distribution of allelic frequencies can be obtained from Δ*. The latter can then be characterized semi-explicitly as the solution to a fixed-point equation. Finally, we find tractable expressions for several macroscopic observables of interest (genetic variance, description of the fluctuations of the trait mean as an Ornstein-Uhlenbeck process). The orders of magnitude of the macroscopic observables can be derived on a wide region of the parameter space. The model shows good fit and can straightforwardly be extended to accommodate pleiotropy, dominance, and some forms of epistasis. We also discuss the different types of breakdown which may occur (Bulmer effect, Hill-Robertson effect, breakdown of the Ornstein-Uhlenbeck approximation for the dynamics of the trait mean, depletion of genetic variability due to low mutation rates).

---

The polygenic limit can be defined as the limit obtained from an explicit individual-based model of a biological population when the number of loci *L* influencing a polygenic trait is large, and the number of organisms *N* within the population is large. There are several important tools that can be used to describe such a population. When the trait is sufficiently additive, the infinitesimal model [26, 4] is typically used to describe inheritance of the trait over a small number of generations. Under the infinitesimal model, the offspring of two parents with respective trait values *z*^*A*^ and *z*^*B*^ has trait value normally distributed around (*z*^*A*^ + *z*^*B*^)*/*2, with variance determined only by the inbreeding coefficient between the two parents and a population-wide parameter called the segregation variance *V*_*S*_. This model can be used to describe the inheritance of any additive trait as long as the number of segregating loci determining this trait is large (and no locus has a disproportionate effect on the trait), even in a relatively small population.

Gaussian selection is typically considered a natural assumption to describe the evolution of morphological traits in a stationary population [73]. From the infinitesimal model, assuming the population size is large and selection is not too strong and Gaussian, one naturally concludes that the trait distribution within the population is Gaussian (the mathematical literature on this subject is growing, e.g [61, 15]). Furthermore, Lande stated in [45] that the population trait mean under Gaussian selection would fluctuate as an Ornstein-Uhlenbeck process with variance determined by the strength of selection.

This trait-based approach, when used to describe a population at stationarity, requires an effective way to predict the value of the segregation variance *V*_*S*_ from microscopic parameters such as mutational parameters and selection parameters. Since the genes determine the trait and only the genes are transmitted, the genetic composition of the population evolves autonomously, without the need to specify any auxiliary variable, and from it the trait distribution should be deduced. Various approaches have been developed (see for instance [89], Chapter 28). Of these approaches, the most remarkable one, which models the joint effect of selection, mutation and genetic drift, takes the gene’s-eye view (in the sense discussed in [1]). This approach explicitly models the dynamics at one or several given loci, typically using diffusion approximations, and uses a mean-field approximation to deduce the genetic variance within the population at stationarity. This approach was notably applied to describe a stationary population under Gaussian selection in [67, 12, 18, 86, 75]. In a spatial setting, it was used in [96, 79] for directional selection and in [64] for Gaussian selection.

Most discussions of this subject assume no trait mutational bias. This means they assume that mutations have zero mean on average. This is hard to justify from a theoretical standpoint, as this would imply that mutation favors the same trait values as selection, which would be a strange coincidence, in particular if this were satisfied across distinct environments (as noted for instance in [18, 19, 90, 68, 51]). In fact, empirical evidence for trait mutational bias is abundant [30, 53, 74, 33, 94, 93, 82] (see Discussion 3.4).

The first theoretical work to consider stabilizing selection with biased mutations is Kimura’s [41] article, but his mathematical analysis of bias was inaccurate, and was only corrected in [90]. Both articles adopt the “continuum-of-alleles” perspective in which a population at equilibrium has infinitely many segregating alleles at every given locus (which is now deemed an unrealistic description of natural populations [73]), and neglect genetic drift. A biallelic model for Gaussian selection with mutational bias was notably discussed by Brian Charlesworth in [18, 19], in a simplified setting in which all loci have the same effect on the trait and the same mutational parameters (mutational bias is also included in [86], but no explicit method is given to compute the stationary distribution at a given locus). Here, we extend this model to a setting in which each biallelic locus has its own mutation rates (with a corresponding mutational bias) and genetic effect. The difficulty of this approach is that genes are coupled in a nontrivial manner by recombination and selection. Now, if we assume that genes are in perfect Hardy-Weinberg linkage equilibrium (HWLE, see SI 6.2 for a mathematical discussion about the domain of validity of this assumption), then selection remains as the only coupling force–the selective advantage of an allele at a given locus depends on the entire trait distribution, which itself depends only on allele frequencies at all other loci.

We show using various mathematical tools (diffusion approximation, mean field theory, fixed-point equations, slow-fast approximations) that this dependence vanishes with the number of loci. This lets us characterize properties of the trait distribution that emerge directly from the sole description of genes, such as the genetic variance, the distance to the selection optimum, and the magnitude of the fluctuations of the population mean trait value at statistical equilibrium–though our model can in principle be extended to describe the dynamics out-of-equilibrium. The distribution of allele frequencies can itself be obtained from a single observable Δ* which solves a simple fixed-point equation. Our derivation is self-contained in the appendix.

Our computations distinguish three regimes which we call weak, moderate and strong selection, based on the value of the **selection-drift ratio [51]** defined as a product of the effective population size, the strength of Gaussian selection, the squared average effect size of a locus and the number of loci (see (4)). Moderate selection describes a wide array of selection strengths for which we expect the same underlying genetic architecture in the limit (as will be seen in Fig. 3), strong selection corresponds to the transition between moderate selection and pervasive loss of allelic diversity due to selection, and weak selection is the transition between the genetic architecture under moderate selection and that in the absence of selection.

## 1 Description of the population model

### 1.1 The individual-based model

We consider a large, panmictic population composed of *N* diploid organisms, displaying an additive quantitative trait *z* controlled by *L* biallelic loci. We assume that *N* ≫ 1 and *L* ≫ 1.

A genotype *g* = (*g*_*l*_)_*l*∈[*L*]_ determines the value *Z*(*g*) of the trait as

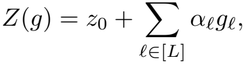

where *g*_*l*_ ∈ {0, 1, 2} is the gene content and *α*_*l*_ ≥ 0 is the effect of the trait-increasing allele at locus _*l*_. We define the **mean allelic effect** 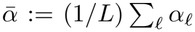. The model can accomodate extensions to polyploidy, pleiotropy, dominance or some forms of epistasis (see SI 10). We assume that *z*_0_ = 0 without loss of generality.

We define the probability of mutation per generation from the trait-decreasing allele to the trait-increasing allele at locus _*l*_, and back, as 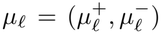. We assume that the (*α*_*l*_, *µ*_*l*_)_*l*∈[*L*]_ are exchangeable, such that (*α*_*l*_, *µ*_*l*_) has a prescribed distribution which is not heavy-tailed, so that in particular there are no large-effect mutations.

The logfitness of an organism with trait value *z* is given by

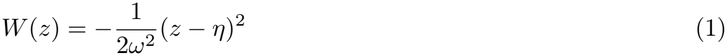

with *ω*^*−*2^ the **selection strength** and *η* the **selection optimum**.

Reproduction occurs via Wright-Fisher sampling: at each generation, organisms independently choose two parents with probability proportional to their fitness and the two parental genomes recombine. Importantly, model details regarding recombination do not matter, but we assume that recombination is strong enough to allow us to neglect linkage disequilibrium (this assumption is discussed in SI 6.2). Our simulations assume all sites are found on a single chromosome with evenly-spaced loci, in which a single crossover occurs per genome per generation.

### 1.2 Three trait values

As was discussed for instance in [51], a model studying the dynamics of a polygenic trait under Gaussian selection from the gene’s-eye view distinguishes three important theoretical trait values. The first is the **selection optimum** *η*. The second is the **mutational equilibrium** *z*_*M*_ defined as the mean trait value that the population would reach at equilibrium if selection were completely absent

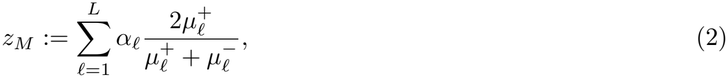

because 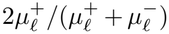 is the mean value of *g*_*l*_ at mutational equilibrium. This value has no *a priori* reason to coincide with *η*, and we call *z*_*M*_ ^−^ *η* the **trait mutational bias**. The third is the **heterozygous trait value** *z*_*H*_ , which is the trait value of an organism heterozygous at every locus

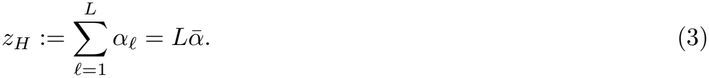

We may assume without loss of generality that the trait is measured in units such that *z*_*H*_ = 1 so that in particular 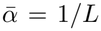. Other key ecological observables are represented in Fig. 1 and are explicitly defined in Section 2.

**Figure 1:**
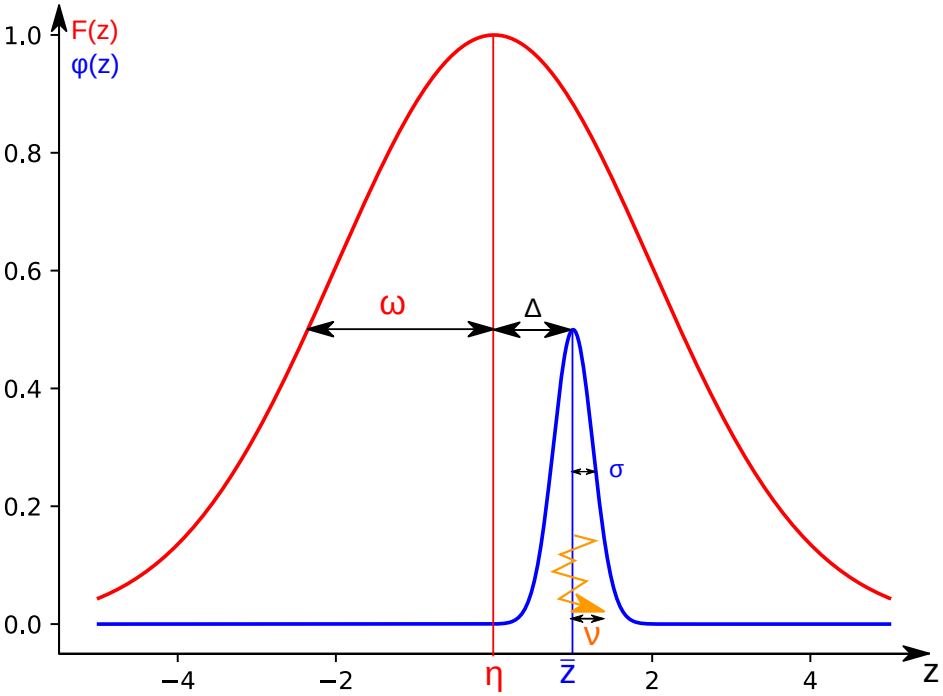
Phenotypic observables. The wide curve is the Gaussian fitness function *F* (*z*) = *e*^*W* (*z*)^, which has typical width *ω* and is centered on *η*. The narrow curve is the distribution of the trait in the population, which has mean 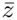 and variance *σ*^2^. The deviation of 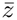 from the optimum is 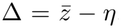. The broken line represents the fluctuations of 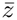 through time, which have magnitude *v*.

### 1.3 Assumptions

The details of our explicit quantitative assumptions are given in SI 2. Some assumptions are made for mathematical convenience, others are necessary to reach the polygenic limit, defined as the regime in which allelic frequencies obey the polygenic equation in the form of a special kind of Wright-Fisher diffusion (see below eq (12)). Here, we informally explain the underlying philosophy.

The mathematically convenient assumptions are as follows. Define 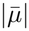 the mean mutation probability

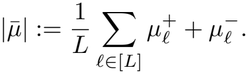

We assume that (A1) all loci have mutation probabilities of order 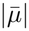 and allelic effect *α*_*l*_ 1*/L*, so that no one locus disproportionately contributes to the trait or the genetic variance. We assume that (A2) mutations are weaker than genetic drift 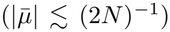 and that (A3, A6) the mutational bias is not too large or too small. We assume that (A4) the selection optimum *η* is accessible, meaning many different genotypes can realize the selection optimum (in particular, the population close to the optimum will not be depleted in genetic variability). In practice, this means *η* belongs to the interval (0, 2), and is not too close to the boundaries 0 and 2. We assume that (A5) selection is strong enough to have a detectable footprint on the genes, but not so strong as to completely deplete genetic variability in the trait (that is, *L* ≲ 2*Nω*^*−*2^ ≲ *L*^2^*N*).

We argue in SI 6 that the equations we obtain further require a sufficiently large population size and a sufficient mutational input to provide an accurate description of the system.

### 1.4 Diffusion approximation

We let 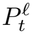 denote the frequency of the trait-increasing allele at locus _*l*_, where time is now measured in units of 2*N* generations; that is, 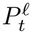 represents the frequency at generation ⌊2*Nt*⌋. We define

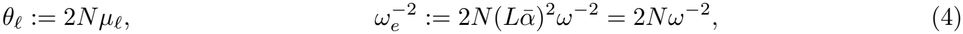

where in the last equality we used *ā* = 1*/L*. We call *θ*_*l*_ the **mutation rate**, and 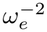 the **selection-drift ratio**. From now on, the model is parameterized by (*N, L, ω*_*e*_, *η*, (*θ*_*l*_)_*l*∈[*L*]_, (*α*_*l*_)_*l*∈[*L*]_).

As explained in Supplementary Information (SI 3), neglecting linkage disequilibrium allows approximating allelic frequency dynamics of the individual loci by Wright-Fisher diffusions which are only coupled by selection

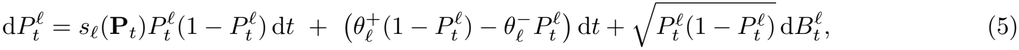

where 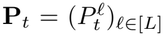 and (*B*_*l*_)_*l*_ are independent Brownian motions. Classically, the selection coefficient *s*_*l*_ is the regression coefficient of the fitness *W* on genotype *g*_*l*_

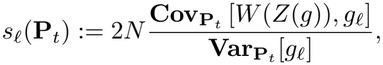

where for **p** = (*p*_*l*_)_*l*∈[*L*]_ ∈ [0, 1]^*L*^, **Cov**_**p**_ and **Var**_**p**_ are the covariance and variance of a vector of independent random variables *g* = (*g*_*l*_)_*l*∈[*L*]_ such that *g*_*l*_ is *Binomial*(2, *p*_*l*_) (Hardy-Weinberg Linkage Equilibrium). We can show that *s*_*l*_ (see SI 3.2) satisfies

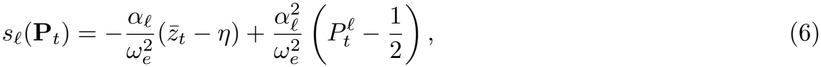

where 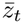 is the mean trait value at time *t*

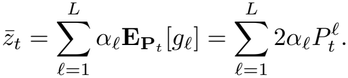

Here, **E**_**p**_ is the expectation associated to the random variable *g* just defined with law determined by **p**.

As a result, allelic frequencies 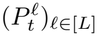 are now coupled only by their weighted mean 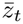. Defining the deviation to the optimum

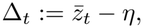

we can rewrite the selection coefficient as

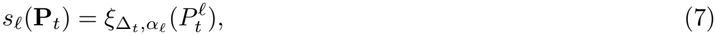

where for *δ* ∈ ℝ, *a* ∈ ℝ_+_ we set

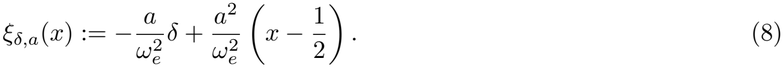

This selection coefficient was computed in earlier works, dating back to Wright in [91]. The first term is a directional selection term which pulls the mean trait value towards the optimum, selecting trait-increasing alleles when Δ_*t*_ *<* 0 (the trait is below the optimum) and trait-decreasing alleles when Δ_*t*_ *>* 0 (the trait is above the optimum). The second term is known as Robertson’s underdominant effect [67], though it dates back to Wright’s earlier article [91]. In [32] (their equation (7)), the selection coefficient is found to be (in our notation)

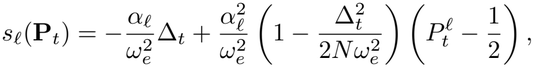

and in particular Robertson’s underdominant term does not have the same form as in our work. This is due to the fact that they consider an out-of-equilibrium regime in which Δ_*t*_ can be very large. Since we are focused on equilibrium, we have (see Table 1) 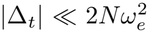 , so their correction term is negligible. When considering fluctuating selection (under which the optimum *η* is replaced by a stochastic process), as in [9], this correction may become significant, but this is beyond our article.

**Table 1:**
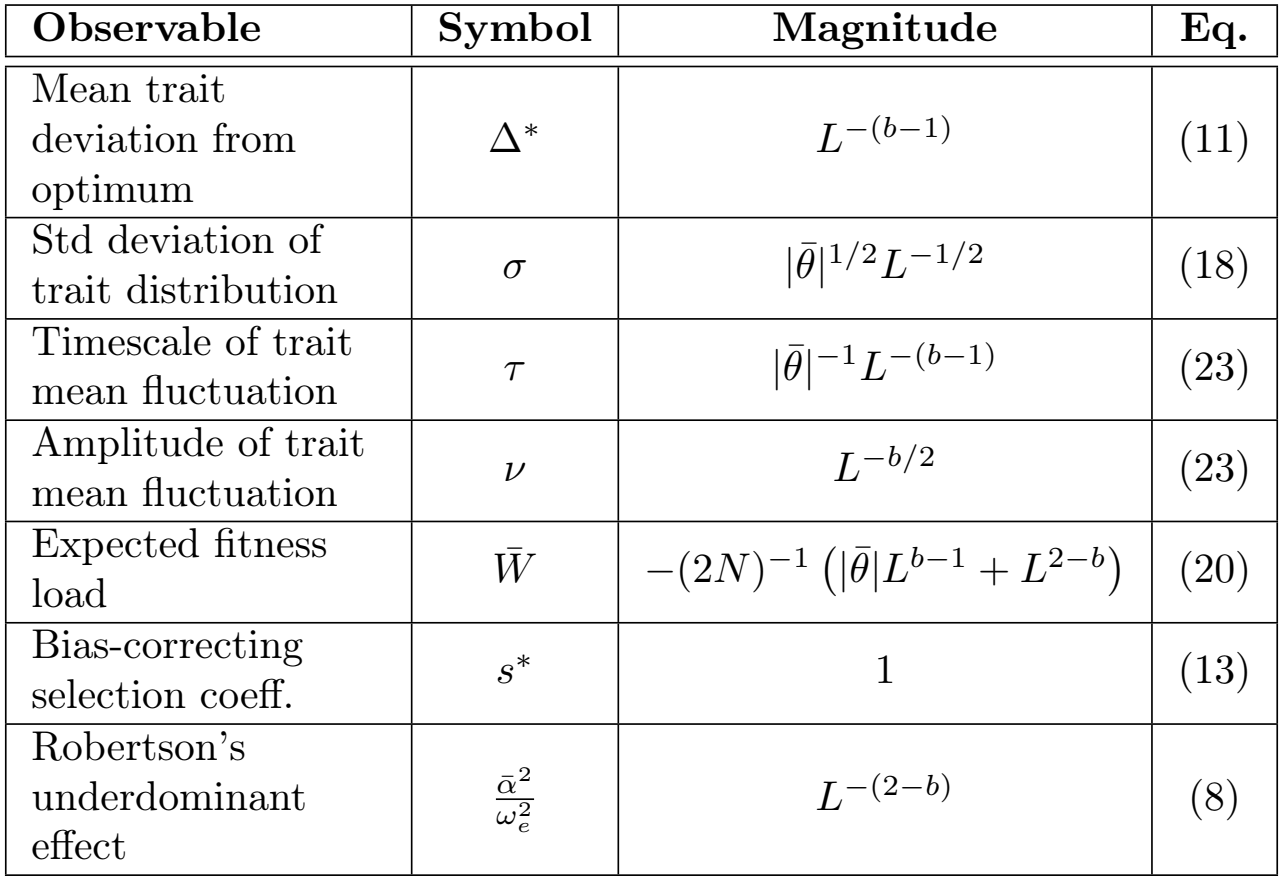
Order of magnitude of macroscopic observables as a function of the number *L* of loci and *b* ∈ [1, 2], assuming 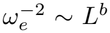 , interpolating between the weak regime (*b* = 1), the moderate regime (*b* ∈ (1, 2)) and the strong regime (*b* = 2). This table is derived in SI 5.2.

| Observable | Symbol | Magnitude | Eq. |
|---|---|---|---|
| Mean trait deviation from optimum | Δ<sup>*</sup> | L<sup>−(b−1)</sup> | (11) |
| Std deviation of trait distribution | σ | \|θ̄\|<sup>1/2</sup>L<sup>−1/2</sup> | (18) |
| Timescale of trait mean fluctuation | τ | \|θ̄\|<sup>−1</sup>L<sup>−(b−1)</sup> | (23) |
| Amplitude of trait mean fluctuation | ν | L<sup>−b/2</sup> | (23) |
| Expected fitness load | W̄ | −(2N)<sup>−1</sup> (\|θ̄\|L<sup>b−1</sup> + L<sup>2−b</sup>) | (20) |
| Bias-correcting selection coeff. | s<sup>*</sup> | 1 | (13) |
| Robertson's underdominant effect | $\frac{\bar{\alpha}^2}{\omega_e^2}$ | L<sup>−(2−b)</sup> | (8) |

The equation (5) for 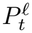 can be further simplified using a mean-field approximation.

## 2 Macroscopic observables from gene’s-eye view

### 2.1 Mean field approximation on the locus dynamics and bias-correcting selection coefficient

*Mean-field theory* is a powerful approach for analyzing complex systems composed of a large number *L* of interacting components. When the constituents interact through a weighted mean of their individual characteristics, → ∞ as *L* this weighted mean behaves deterministically as by the law of large numbers, and the constituents effectively behave independently [16].

In the case where the allelic effects and mutation rates (*α*_*l*_, *θ*_*l*_) ≡ (*α, θ*) are equal across loci, by replacing the mean with its expectation, we get

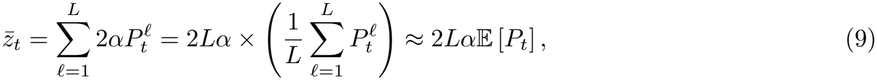

where *P*_*t*_ has the law of 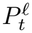 (which here does not depend on _*l*_). When the (*α*_*l*_, *θ*_*l*_) are now distributed, we replace the *weighted* mean with an ‘augmented’ expectation which includes averaging also over the distribution of (*α*_*l*_, *θ*_*l*_)

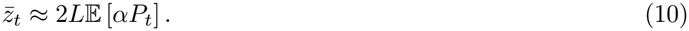

In other words, we define the random vector 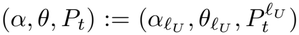 where _*lU*_ is uniformly distributed on [*L*], as the typical locus. The expectation E can now be taken at a specific locus (for instance 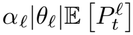), or at the typical locus (for instance E [*α θ P*_*t*_]).

We can now define Δ* as the **mean deviation at statistical equilibrium**

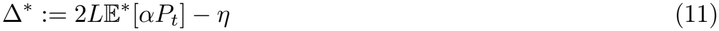

where E* is the expectation at equilibrium, which is also the time average

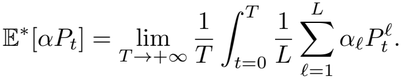

Using a mean-field approximation, we may replace Δ_*t*_ with Δ* in the selection coefficient in (7). The allelic frequency 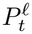 at each locus _*l*_ then independently obeys the following stochastic differential equation which is autonomous in the sense that each term only depends on *P*_*l*_ itself and Δ*

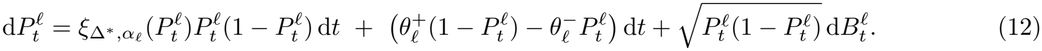

We call (12) the polygenic equation, and the regime in which it is accurate the polygenic limit. A formal justification for this mean-field approximation is spelled out in Section 2.4, where we also describe the dynamics of the trait.

Let us briefly discuss the expression of 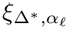. In the absence of selection, mutations naturally drive the trait to the mutational equilibrium *z*_*M*_ defined in (2). In view of (8), this occurs when 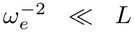. Then Δ* *≈ z*_*M*_ *η*, and selection is negligible at the genetic and trait levels (see SI 5.4). As soon as 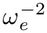 is of order *L* or higher, what shapes the value of Δ* is the balance between mutation (pushing 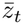 to *z*_*M*_) and selection (pushing 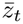 to *η*).

We may now introduce the following important quantity

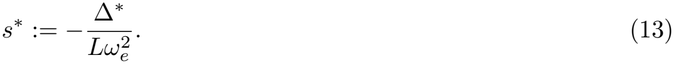

In particular, the directional selection term of the selection coefficient 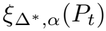 is *Lαs*\*, with expectation *s*\*. We may interpret *s*\* as a *bias-correcting selection coefficient*. It reflects the selective advantage of trait-increasing alleles when Δ* *<* 0 (the trait is on average below the optimum) and their disadvantage when Δ* *>* 0 (the trait is on average above the optimum). Furthermore, the magnitude of *s*\* is informative: *s*\* is the strength of directional selection that should be imposed on the system to bring the mean trait value from its mutational equilibrium *z*_*M*_ to 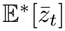 , which is itself a compromise between mutation and selection. In particular, *s*\* depends on the strength of selection 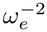 , both through the denominator in (13) and the observable Δ*. Through Δ*, *s*\* also depends on the mutational bias *z*_*M*_ ^−^ *η*.

### 2.2 Bridging microscopic and macroscopic scales to solve for the deviation to the optimum

The consistency between microscopic (locus-level) and macroscopic (trait-level) dynamics is central to the meanfield approach and allows for an explicit determination of Δ* as the solution to a fixed-point problem.

In (12), take (Δ*, *α*_*l*_, *θ*_*l*_) = (*δ, α, θ*). The stationary density of this diffusion [40] is then given for *p* ∈ (0, 1) by

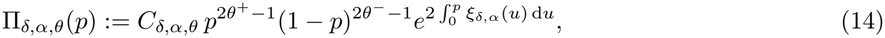

where *C*_*δ,α,θ*_ is a normalizing constant. Using the mean-field approximation to average trait contributions across loci, from (11) we deduce that Δ* satisfies the following fixed-point equation:

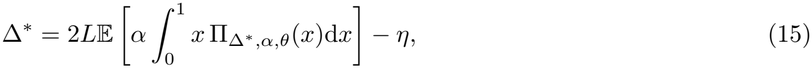

where the expectation is now only taken over the distribution of (*α*_*l*_, *θ*_*l*_). The latter relation embodies a *self-consistency condition* that allows tying the dynamics of allelic frequencies with macroscopic observables. We deduce from this fixed-point equation in SI 5.2 that (except when 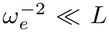 , where Δ* ≈ *z*_*M*_ ^−^ *η*) the mean trait deviation always scales as

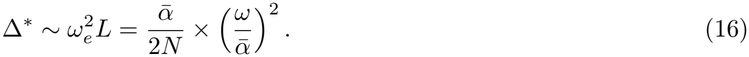

This equation can be interpreted as follows. We can see 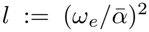 as a number of loci such that Δ* corresponds to a change of frequency of order 1 over *l* loci. It is as if selection tolerated *l* maladapted loci pushing the population towards the mutational equilibrium. In particular, stronger selection implies smaller *l*, and therefore less tolerance to maladaptation.

Similarly, we deduce in SI 5.2 that the trait variance *σ*^2^ can be expressed as a result of the independent, genetic additive contributions

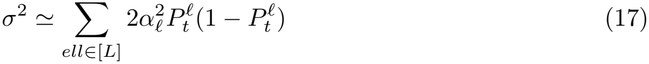

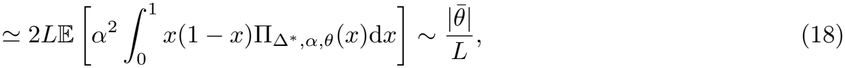

where 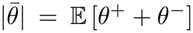 is the mean mutation rate. We plot in Fig. 2 the result of simulations using the individual-based model against our theoretical predictions for different selection strengths. As expected, stronger selection leads to a smaller distance to the optimum Δ*. We also see that when selection is sufficiently strong, it depletes the genetic variance *σ*^2^. More precisely, we see that when selection is weak, our theoretical predictions for Δ_*t*_ and *σ*_*t*_ match simulation results even when *L* and *N* are as low as *L* = 100 and *N* = 50. On the other hand, when selection is strong, we see a mismatch in the genetic variance *σ*^2^ due to the build-up of negative Linkage Disequilibrium (LD): this is the Bulmer effect [13]. The mismatch is decreased when the population size is increased (*N* = 500). Furthermore, the decrease in *σ*^2^ due to the Bulmer effect decreases the effectiveness of the bias-correcting coefficient of selection, increasing the equilibrium distance to the optimum Δ* with respect to expectations. Following the Quasi-Linkage Equilibrium approach [57, 56], we argue in SI 6.2 that the Bulmer effect (in a setting with uniform crossover) can be neglected if 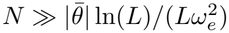. In the worst-case scenario of strong selection 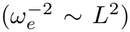 , this requires 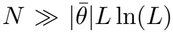). If we ignore the ln(*L*), this means the Bulmer effect can be neglected as long as the population size *N* is much greater than the number of segregating loci. We develop a first-order correction for the Bulmer effect in SI 10. The method is based on the Quasi-Linkage Equilibrium approach [57, 56], and yields a good approximation (see Fig. S8).

**Figure 2:**
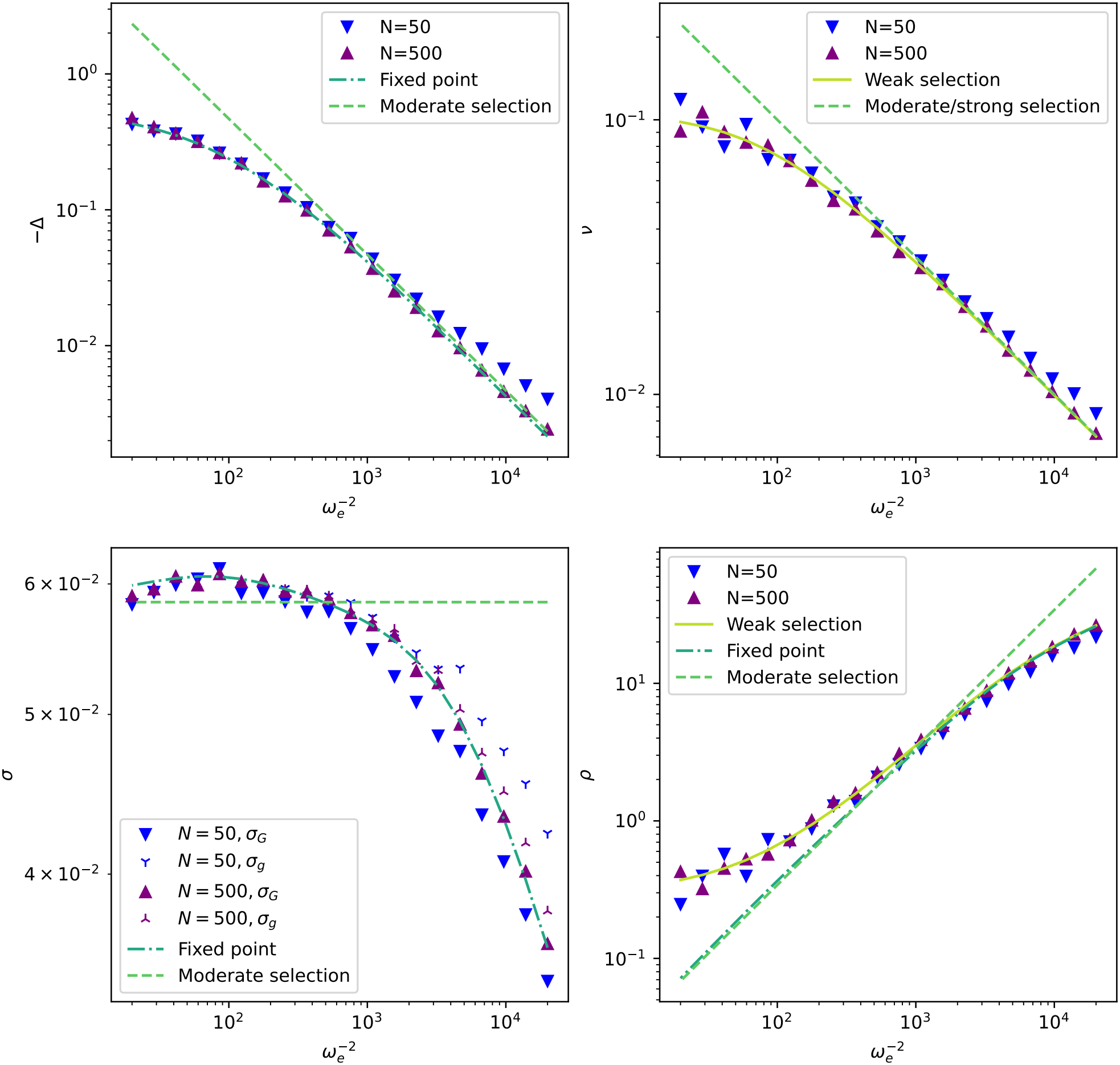
Comparison of theoretical predictions with numerical simulations of populations with size *N* (*N* = 50 or *N* = 500) for the macroscopic observables of Fig. 1 for varying selection regimes, from weak selection (left) to strong selection (right). The simulations were carried out with *L* = 100, the same mutation probabilities for all loci *θ* = (0.1, 0.2), and allelic effects (*α*_*l*_)_*l*∈[*L*]_ exponentially distributed with parameter *L* (in particular *ā* = 1*/L*), so that *z*_*M*_ = 2*/*3. The selection optimum is *η* = 1.2. The same (*α*_*l*_)_*l*∈[*L*]_ were used in all simulations, and the predictions were made conditional on the (*α*_*l*_)_*l*∈[*L*]_. The simulations were run for *T* = 1, 000*N* generations, and each observable was measured as an average over the last 500*N* generations. The fixed-point predictions are derived from (15) in SI 5.1. The moderate selection predictions are derived from the fixed-point equation under the assumption 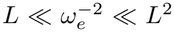 (see SI 5.2). For the magnitude of the fluctuations *v* and the autocorrelation parameter *ρ*, the predictions for weak selection use corrections derived in SI 6.5. For the genetic variance, the simulation results distinguish between the genetic variance in the trait within the population 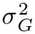 (filled triangles), and the genic variance 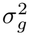 (three-pointed stars), which is the variance in the trait if the population was in linkage equilibrium (neglecting correlations between pairs of loci). It is computed using (17).

**Figure 3:**
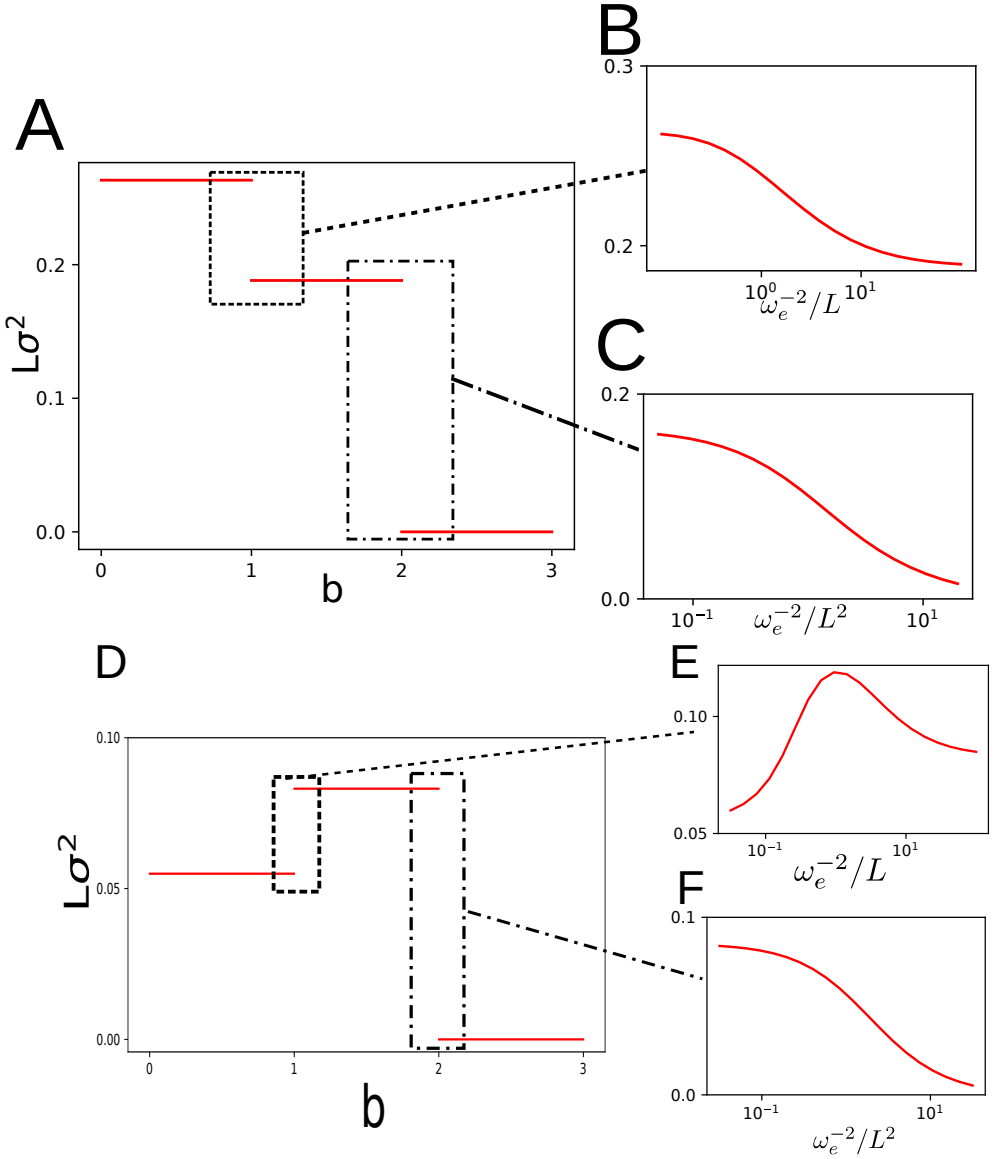
We take the theoretical limit *L, N* → +∞ , and plot the predicted rescaled genetic variance *Lσ*^2^ at equilibrium as a function of the selection power *b* (see (19)). In **A**,**B**,**C** there is no mutational bias (*θ*_*l*_ = (0.1, 0.1)). **D**,**E**,**F** are equivalent to **A**,**B**,**C**, but with strong mutational bias (*θ*_*l*_ = (0.01, 0.1)). **A** We expect the rescaled genetic variance *Lσ*^2^ in the limit to converge to a step function of the parameter *b*. When *b <* 1 the genetic variance is equivalent to that of a neutral model (no selection), when *b >* 2 we expect the genetic variance to be completely depleted, and when *b* ∈ (1, 2), *Lσ*^2^ converge to a specific value. At the discontinuity *b* = 1, many different values of *Lσ*^2^ are possible depending on how 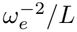 scales with *L*, which corresponds to what we call the regime of weak selection. We zoomed into this window in **B.**Similarly, at the discontinuity *b* = 2 many different values of *Lσ*^2^ are possible depending on how 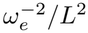 scales with *L*, which corresponds to the regime of strong selection, and is zoomed into in **C**. When *b* ∈ (1, 2), we are in the regime of moderate selection. Note the contrast with Fig. 2 (in which *L, N* have finite values): in the latter, there is no clear window in which the moderate selection regime is accurate, whereas in **A** it occupies the whole interval *b* ∈ (1, 2). In all figures, we took *η* = 1.5 and let (*α*_*l*_)_*l*∈[*L*]_ be exponentially distributed with parameter *L* (in particular 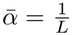). Details on the numerical approximations used to solve the fixed-point equation (15) are available is SI 12

### 2.3 Selection regimes

#### Persistence of the bias-correcting coefficient *s*\* across scales and the moderate selection regime

The mean selection coefficient of trait-increasing alleles *s*\* (defined in (13)) is proportional to the product of the trait deviation Δ* from the optimum and of the strength of selection 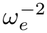. At the trait level, we see from (16) that the strength of selection has a direct effect on Δ* which keeps the product 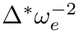 of order *L*. As a result, *s*\* remains of order 1 as long as 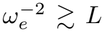 (see also SI 5.4). In other words, the mutational bias always has a substantial effect at the locus level, of the same order as genetic drift. When 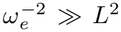 , Robertson’s underdominant effect becomes leading 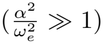 , resulting in a depletion of genetic variance with a high concentration of allelic frequencies at 0 and 1. In this “ultra-strong” selection regime, the outcome of competition between two alleles at a given locus depends on which is more frequent, rather than on the sign and magnitude of their allelic effects.

In the case when 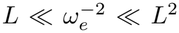 , something remarkable occurs: *s*\* becomes independent of *ω*_*e*_ (SI 5.3). This is because, on one hand, the law of a locus only depends on *s*\*, since Robertson’s underdominant term (second term in (8)) is negligible 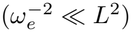. On the other hand, since Δ* is of order 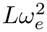 , we may neglect Δ* in the left-hand side of the fixed-point equation (15), which then reads

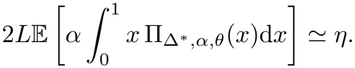

This can then be seen as an equation on *s*\*, independent of *ω*_*e*_. This motivates us to label the regime *L* 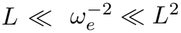 as a **moderate selection regime**. We are then led to label the regime 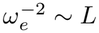 as **weak selection**, and the regime 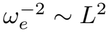 as **strong selection**.

In the moderate selection regime, the same genetic architecture, and accordingly similar macroscopic observables, can arise for very different values of 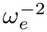. We illustrate this in Fig. 3, in which the theoretical prediction for the rescaled genetic variance *Lσ*^2^ is plotted in the limit *L* → +∞ , as a function of the **selection power** defined with 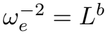, equivalent to

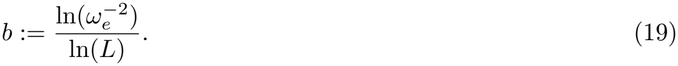

In this figure, it can be seen that *Lσ*^2^ is expected to converge to a step function of *b*, with a single value corresponding to 1 *< b <* 2 and discontinuities at *b* = 1 (weak selection regime) and *b* = 2 (strong selection regime). We also expect Δ\**L*^*b−*1^ to be independent of *b* when *b* ∈ (1, 2) (see Table 1).

#### Observable characteristics of the regimes and the stabilizing transition

The macroscopic properties of the three distinct regimes are illustrated in Fig. 4 and summarized in Table 1. Recall from (18) that the trait population variance *σ*^2^ is always of order 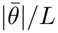 .

**Figure 4:**
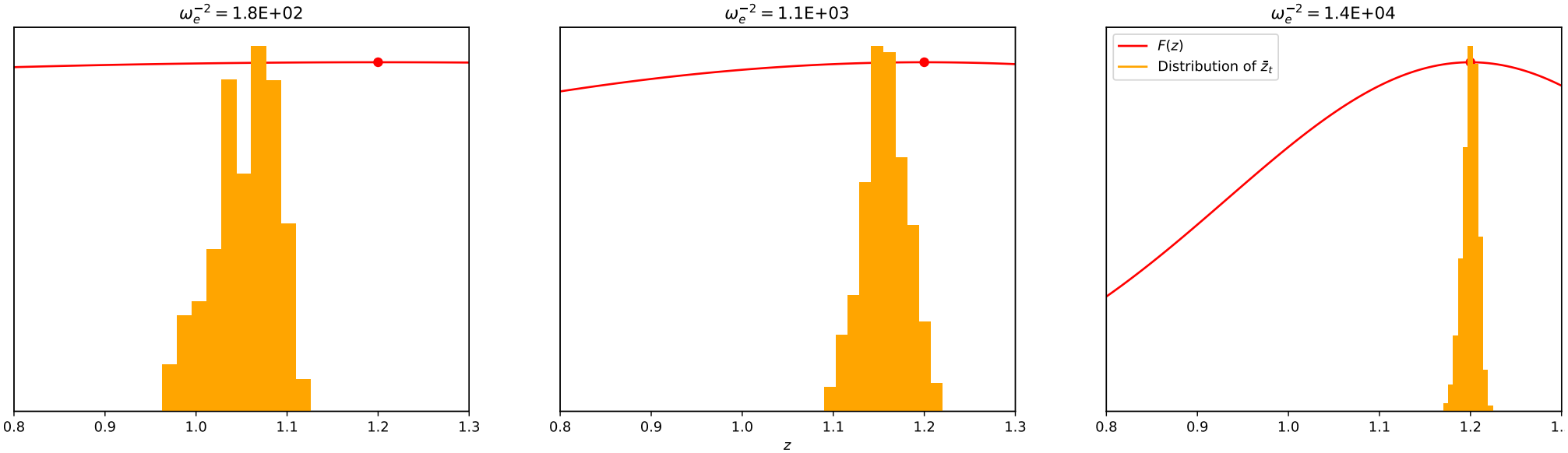
The histogram shows the empirical distribution over time of the population mean trait 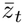 (not a snapshot of the trait distribution at a given time). For three different values of *ω*_*e*_ corresponding respectively to weak, moderate and strong selection, we show a histogram of 10, 000 values of 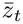 after a burn-in of 24, 000 generations. The parameters used are *N* = 100, *L* = 100, *η* = 1.2, *θ* = (0.1, 0.2), and the parameters (*α*_*l*_)_*l∈*[*L*]_ were sampled with distribution *Exponential*(*L*). We superimpose in red the fitness function *F* (*z*), and the dot corresponds to *η*. In particular, Δ^*\**^ is the distance between the mean of the orange distribution and the red dot, and *v* is the width of the orange distribution. As the strength of selection increases, both Δ^*\**^ and *v* decrease. Only in strong selection (right panel) should we see Δ^*\**^ and *v* being of the same order of magnitude. Furthermore, this distribution is expected to be Gaussian when (Δ_*t*_)_*t≥*0_ behaves as an Ornstein-Uhlenbeck process, which is not expected to hold for weak selection.

1. *Weak selection regime* 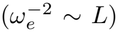 : In this regime, the population is concentrated away from the fitness optimum:

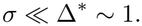
2. *Strong selection regime* (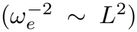 ∼ *L*^2^): Here, the population is tightly concentrated around the fitness optimum:

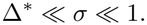

It is only in this regime that Robertson’s underdominant effect becomes significant, so that genes also experience positive density-dependent selection in addition to genic selection against mutational bias.
3. *Moderate selection regime* 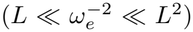 : This regime interpolates between the weak and the strong regime.

More precisely, Table 1 lets us identify an interesting transition: Δ* and *σ* are of the same order only when 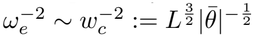. For weaker selection, Δ* dominates *σ*, and for stronger selection *σ* dominates Δ*.

This can be analyzed by looking at the fitness load: elementary computations (SI 5) show that the mean fitness within the population 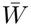 satisfies

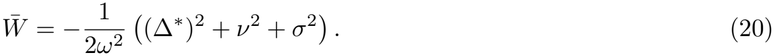

From Table 1, we always have that Δ* ≳ *v*, with the two only of the same order when 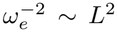 (strong selection), in which case *σ* ≫ Δ*. In particular the mutation load is determined by Δ* and *σ*. For selection strength below the critical value 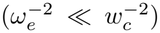 , the fitness load is determined by Δ*, whereas for greater values 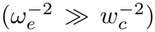 , the fitness load is determined by *σ*^2^. Therefore, it makes sense to qualify the regime below 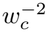 as the *directional dominant regime* and the one above 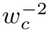 as the *stabilizing dominant regime*, and the critical value 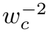 as the **stabilizing transition**. This should not be taken to mean that no directional selection occurs in the stabilizing dominant regime: as was discussed in the previous section, from a locus’s perspective the bias-correcting coefficient is always of order 1. The distinction between directional dominant and stabilizing dominant is based solely on what an ecologist measuring the trait observes (in one case, a population under apparent directional selection, in the other case, a population under apparent stabilizing selection). One important consequence of the stabilizing transition is that the strength of selection 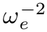 cannot be inferred when 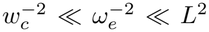 , neither from macroscopic data, because Δ* ≪ *σ*, nor from genomic data, because the distribution of *P*_*t*_ is independent of 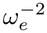. In terms of the selection power *b* from (19), this corresponds to 3*/*2 *< b <* 2 when 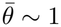. This is illustrated in Figure 5.

**Figure 5:**
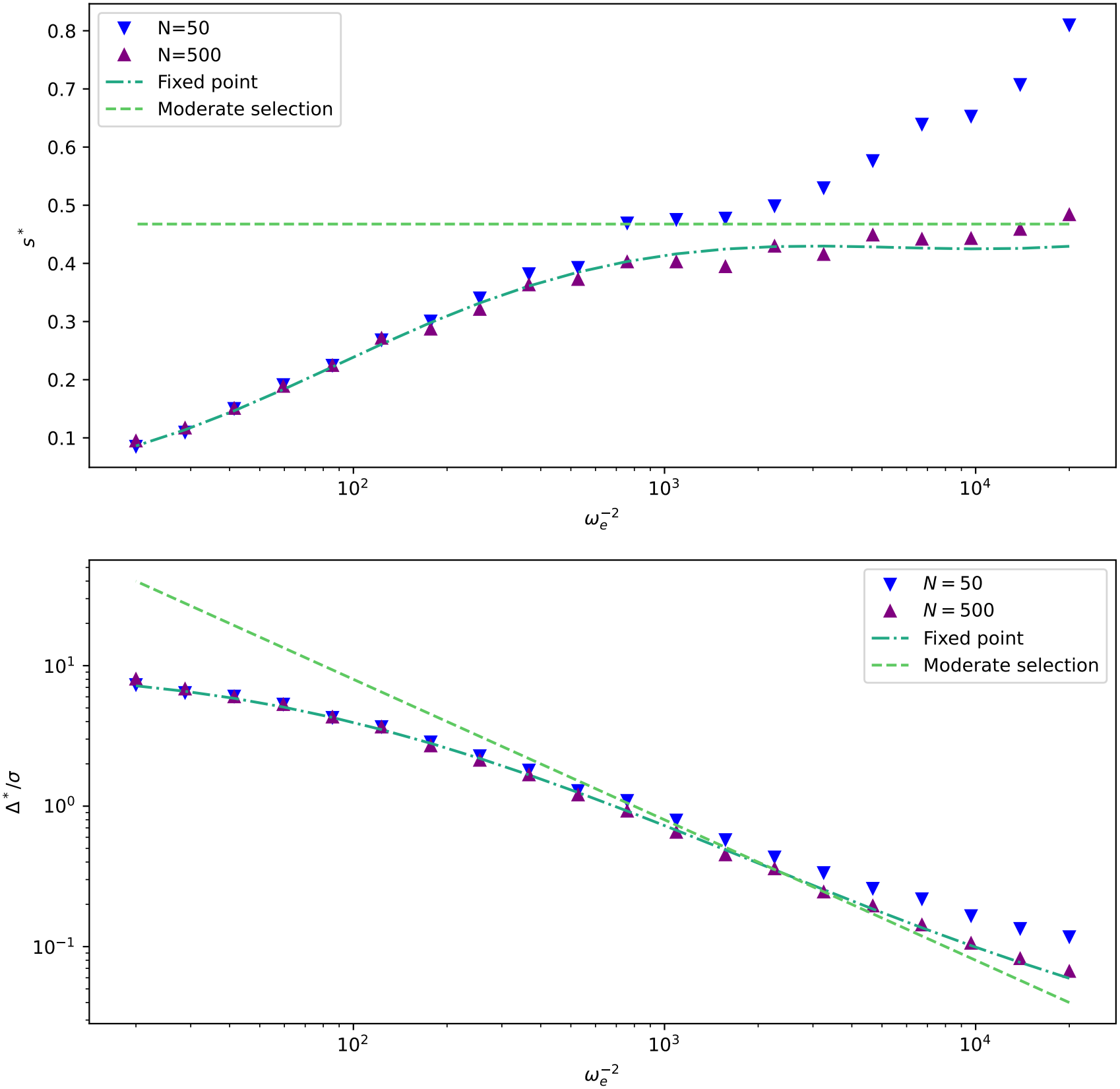
For the same simulations as in Fig. 2 (*L* = 100, *θ* = (.1, .2), *η* = 1.2), we plot the effective selection coefficient *s*^*\**^ and Δ^*\**^*/σ*. The fixed-point predictions are derived from (15) in SI 5.1. The moderate selection predictions are derived from the fixed-point equation under the assumption 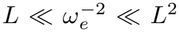 (see SI 5.2). As argued in Section 2.3, *s*^*\**^ is expected to converge to some constant independent of 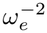 when selection is moderate 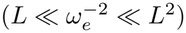. This prediction is more accurate for larger *N*. On the other hand, as shown in Table 1, Δ^*\**^*/σ* decreases for stronger selection, and in particular for sufficiently strong selection is so small that it cannot be reliably estimated. The coefficient *s*^*\**^ could be inferred from the distribution of (*α, P*_*t*_) obtained from GWAS data, and Δ^*\**^*/σ* could be inferred from phenotypic data.

#### Non-monotonicity of the variance

Another counterintuitive prediction of our model is that *σ* does not always decrease with higher selection pressure. In the absence of selection, when mutational bias is strong, allelic frequencies tend to accumulate close to 0 or to 1 depending on the sign of mutational bias. Increasing selection at the trait level (parameter *b* in Fig. 3, panels D, E, F) induces bias-correcting directional selection at the locus level that recenters the frequencies, thus increasing the population variance. We prove in SI 9 that when the (*α*_*l*_, *θ*_*l*_) are constant across loci, the criterion for some level of weak selection to increase *σ* is that going from the mutational equilibrium *z*_*M*_ to the selection optimum *η* brings the trait closer to the heterozygous trait value *z*_*H*_. That is: *η, z*_*H*_ *< z*_*M*_ or *η, z*_*H*_ *> z*_*M*_. To put it another way, starting from a situation with very weak selection, where 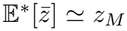 , increasing the strength of selection brings 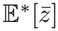 closer to *η*. If, in so doing, 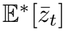 also gets closer to *z*_*H*_ , then the action of selection will lead some loci which the mutational bias kept at one boundary closer to 1*/*2, and *σ* will increase for entropic reasons.

This is important because in articles such as [18, 51], the neutral value of *σ*^2^ (the genetic variance if there were no selection) is often used as a proxy for its actual value, for instance to solve for Δ^*\**^. It is then claimed that this neutral estimate is an overestimate of the actual value of *σ*^2^, which as we see is not true.

#### Heterogeneity across loci

In Fig. 6 we plot the joint distribution of 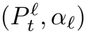, assuming that mutation rates are symmetric (*θ*^+^ = *θ*− = 1*/*2) and that the trait mutational bias *z*_*M*_ −*η* is negative, thereby selecting trait-increasing alleles.

**Figure 6:**
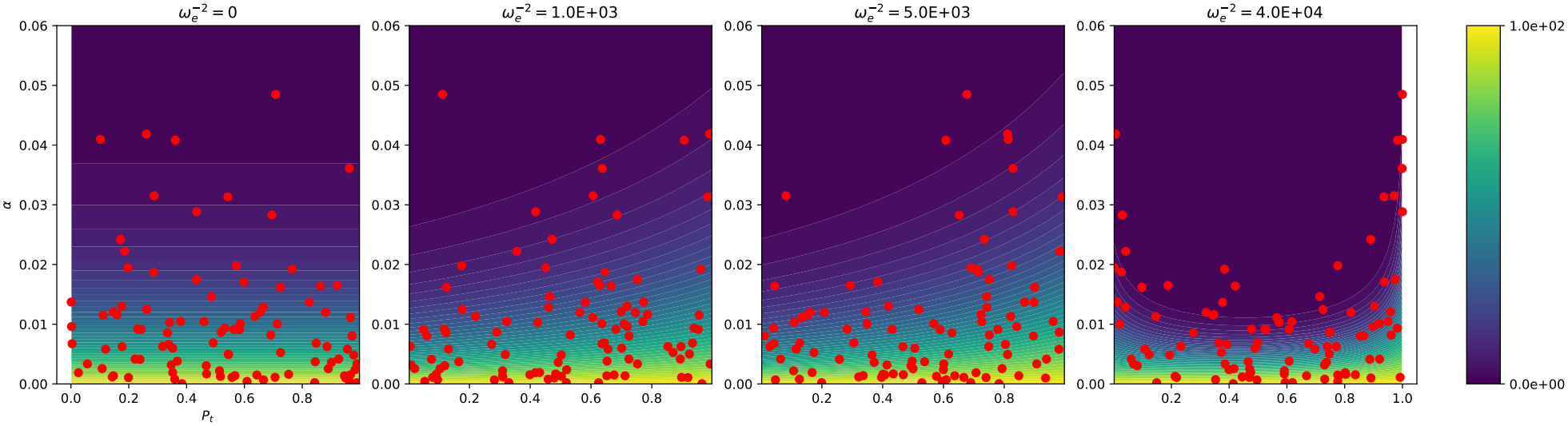
For no selection, weak, moderate and strong selection respectively, we plot the joint distribution of 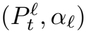 after a single run. We chose symmetric mutation rates *θ* = (0.5, 0.5) to make the transition between the selection regimes more apparent. The other parameters are *L* = 100, *N* = 500, *η* = 1.2 and *α*_*l*_ is distributed as *Exponential*(*L*). The colour plot correspond to the predicted densities in the corresponding regimes.

As expected, the distribution of allelic frequencies is observed to be approximately uniform in the absence of selection and to be biased to the right in the presence of substantial selection. In the moderate regime, the biascorrecting selection term indeed shifts the distribution to the right, effectively counteracting the asymmetry of mutation rates. In the strong regime, the positive frequency-dependent selection driven by Robertson’s underdominant effect (8) results in a U-shape distribution, increasing the weight on extreme allelic frequencies.

### 2.4 Trait dynamics

In the SI 4, we show that in the moderate and strong regimes, the fluctuations of the mean deviation Δ_*t*_ away from its statistical average Δ^*\**^ evolves according to an Ornstein–Uhlenbeck process:

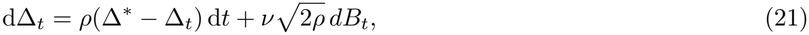

In this equation, *v* captures the magnitude of the stochastic fluctuations and *ρ* the autocorrelation parameter of 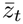 such that

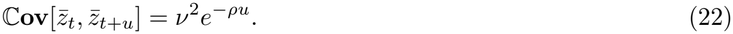

We find in SI 5.2 that

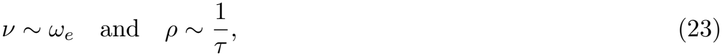

where we define the characteristic timescale of the trait 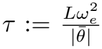 Under weak selection 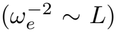, we show in SI 6.5 that (21) is no longer justified mathematically (see Fig. S7). We derive a correction for *v*^2^ and *ρ* which, though not mathematically rigorous, yields a good numerical approximation when mutation rates |*θ*_*l*_ |are constant across loci. See the right panels of Fig. 2 for a comparison between theoretical predictions and individual-based simulations for *ρ* and *v*. Increasing the strength of Gaussian selection decreases the magnitude of the fluctuations of 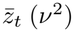, but accelerates them (increase in *ρ*). We plot in Fig. S7 the validity of equation (22) for the autocorrelation: it fits well even for weak selection, in which we expect this equation not to be valid.

This Ornstein-Uhlenbeck process was already obtained in [49], with the same correction to *v* as us (his equation (13c)), but an estimate of Δ^*\**^ which requires especially weak selection.

#### Justifying the mean-field approximation

In Section 2.1, the locus dynamics was obtained by formally replacing Δ_*t*_ with its average Δ^*\**^. How do we justify this approximation? From (6), we have

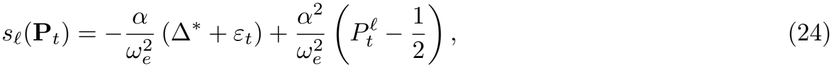

where *ε*_*t*_ := Δ_*t*_ − Δ^*\**^ is the fluctuation of the trait mean.

In the weak/moderate regimes 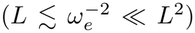 the fluctuations (*ε*_*t*_)_*t≥*0_ are negligible compared to the mean deviation Δ^*\**^ (*v* ≪ Δ^*\**^ in Table 1) and a classical mean-field approximation applies.

The strong regime 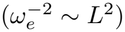 is more subtle and can only be justified by a *dynamical* mean-field approach. As already discussed, fluctuations cannot be ignored (*v*∼ Δ^*\**^, see Table 1) and at first sight, it seems that Δ_*t*_ is not approximately constant equal to Δ^*\**^ in (24). To address this issue, we adopt a *slow-fast approximation [37].* Since the trait mean Δ_*t*_ fluctuates on a much faster timescale than the locus-level dynamics, this separation of timescales leads to a *slow-fast averaging effect*, allowing us to effectively decouple macroscopic (trait-level) and microscopic (locus-level) dynamics. As a result, in (6), we may still, despite non-negligible fluctuations, approximate Δ_*t*_ by Δ^*\**^, now seen as a time average.

### 2.5 Compatibility with the infinitesimal model

The basic infinitesimal model (no dominance, no epistasis) describes the inheritance of trait values under an additive model in which the number of segregating loci is large (*L*≫ 1) and LD is small [4]. Under this model, the offspring of parents with respective traits *z*_1_ and *z*_2_ has trait value normally distributed around the midparent (*z*_1_ + *z*_2_)*/*2, with variance determined by a fixed parameter and independent of (*z*_1_, *z*_2_). It has been recognized that the infinitesimal model only emerges from a Mendelian model in which the number of loci *L* is large under some specific assumptions on the recombination mechanism: free recombination is acceptable, single-crossover is not (in a strict mathematical sense, though it may yield a good approximation). To see why single-crossover does not yield the infinitesimal model (as was discussed in [25], personal communication, in prep), one should consider that a linear genome is best described as *L→* +∞ as a random function (*g*_*l*_)_*l∈*[0,1]_ (see for instance [70]). In this setting (taking constant allelic effects), the offspring of genomes *g*^*a*^ and *g*^*b*^ under single-point crossover has trait value

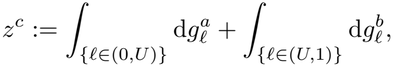

with *U* a uniform random variable on [0, 1]. If we take *g*^*a*^ and *g*^*b*^ to be independent Brownian motions, it can be seen that *z*^*c*^ is not normally distributed.

It is sometimes claimed that under the infinitesimal model, allele frequencies are not affected by selection (for instance, [4] claims that selection must “slightly perturb” the neutral distribution of alleles). If it were so, our model in which the stationary distribution of 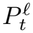is given by 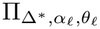 would never satisfy the infinitesimal model. This claim however is only true in the following sense: if we take a population with finite *L, N* and let *L*→ + ∞ *while keeping N constant*, then allele frequencies in the limit are not affected by selection. If we let *L, N* and 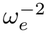 jointly go to infinity (with *N* sufficiently large with respect to *L*, see SI 2), as in our setting, then it is generally still true that the allele frequencies in the limit will not be altered by selection **over the course of a single generation**. The diffusion equation (5), which describes the evolution of allele frequencies, will however still be accurate and have a non-negligible selection term. The infinitesimal model is only used to describe short-term trait inheritance, and is accurate for 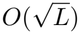 generations, whereas our diffusion approximation (5) measures time in units of 2*N* generations. In fact, as long as significant genetic variance is maintained by a sufficient mutational input, our description of the polygenic limit satisfies most usual assumptions of the infinitesimal model: large number of loci co-determining an additive trait with negligible LD. Depending on the recombination mechanism, trait inheritance within our population in the polygenic limit may or may not satisfy the infinitesimal model (even though allele frequencies are most definitely affected by selection). We illustrate this in Fig. 7, in which the trait distribution in the offspring of a randomly-sampled pair at stationarity is plotted.

**Figure 7:**
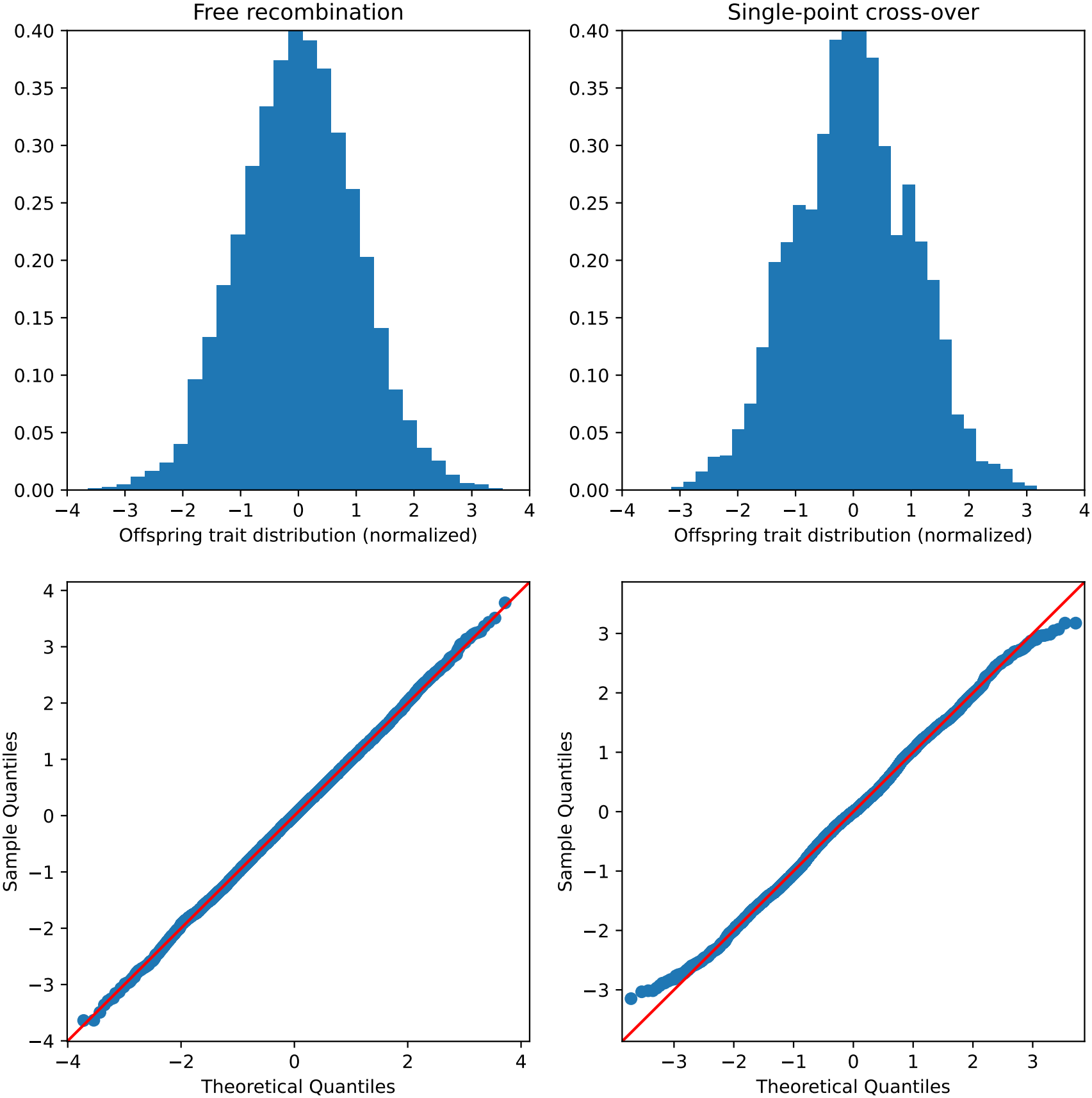
We evolved the population under strong selection for *T* = 5*N* generations. We sampled two organisms at random within the population. Above are represented the trait distribution of offspring of these same 2 organisms, either with free recombination or uniform crossover (rescaled and centered). Below are the qqplots. Under the infinitesimal model, the offspring trait distribution relative to the midparent is normal: this is asymptotically true for free recombination, not so for single crossover. Parameters are *θ* = (0.1, 0.2), *L* = 1, 000, *N* = 5, 000, *η* = 1.2, 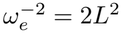 and (*α*_*l*_)_*l∈*[*L*]_ are exponentially distributed.

### 2.6 Breakdown

In SI 6, we discuss the breakdown of our results, which can come from several factors. First, in Section 6.2 we discuss how the assumption of HWLE may break down due to two different causes. The Bulmer effect [11] (discussed in Section 2.2 and SI 10) appears if the population size is too small with respect to the strength of selection. Under this effect, selection generates negative LD by favoring genomes in which genes are negatively correlated. The Hill-Robertson effect [34] similarly results in negative LD when the population size is too small, but may occur also when the trait mean is very far from the optimum, in which case selection is well described as directional. It results from the fact that as LD randomly fluctuates, when it is positive selection and recombination jointly act to deplete it, whereas when it is negative only recombination acts to restore LE (see Fig. S3 and S4).

Second, in Section 6.3 we argue that the mean-field approximation does not hold if the mutation rate is too small, specifically if 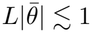. This assumption is crucial to ensure that many loci are segregating.

Third, in Section 6.5, we claim that the description of the fluctuations of (Δ_*t*_)_*t≥*0_ as an Ornstein-Uhlenbeck process in (21) breaks down if the error on the mean-field approximations of (21) is of the same order as the fluctuations of (Δ_*t*_)_*t≥*0_, which occurs for weak selection 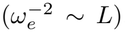. In this scaling, the fluctuations of the mean (Δ_*t*_)_*t≥*0_ depend on the fluctuations of the variance 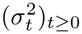, and can therefore no longer be seen as an autonomous Markov process. This breakdown can be corrected in a numerically satisfying way (as in Fig. 2B and D) but not in a mathematically rigorous way (see Fig. S7).

We also illustrate in Fig. S6 the need for the (*α*_*l*_)_*l∈*[*L*]_ to have well-behaved moments. Finite variance seems sufficient for practical purposes, but we recall [60] found rough estimates of tail exponents in additive effects between 1 and 2.5 (see also [77]).

Of note, we do not observe qualitative breakdown in which the population dynamics completely shift to genotype selection, described as clonal condensation in [58]. We also do not observe the appearance of sweeps at individual loci as was described [35]. It seems both these phenomena (which were to our knowledge not studied under Gaussian selection) can only occur in the “ultra-strong” selection regime 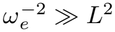, when selection completely overwhelms genetic drift, or in the out-of-equilibrium dynamics if the system is originally very far from the optimum [36].

### 2.7 Extensions

Our fixed-point approach can be extended considerably. In SI 10.1 we mention polyploidy, for which the results are essentially unchanged, replacing 2*N*_*e*_ with *kN*_*e*_ where *k* is the ploidy. In SI 10.2 we add pleiotropy: each allele influences *d* traits. Future work could study this behavior when *d*≫ 1, which should agree with the conclusions of [75, 82].

#### 2.7.1 Dominance

In SI 10.3 we add dominance: this means the trait function *Z* is now of the form

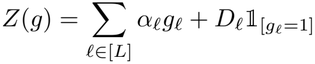

where *D*_*l*_ is the dominance coefficient at locus*l*. The selection coefficient on locus*l* is then 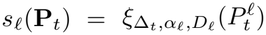 where for *δ* ∈ ℝ, *a* ∈ ℝ_+_, *D* ∈ ℝ we define

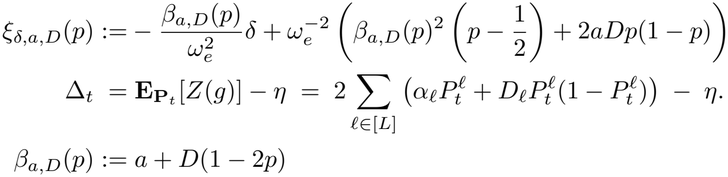

It is remarkable that whereas 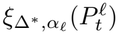 only was a polynomial of degree 1 in 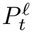 in the additive case above (8), in the dominance case 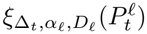 is a polynomial of degree 3 in 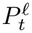.

Replacing Δ_*t*_ with Δ^*\**^ := E [Δ_*t*_], we find the following approximate equation 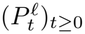

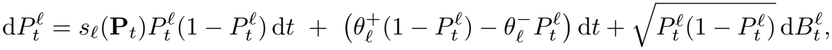

From there, we find the stationary distribution of 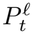to be 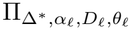 where we define

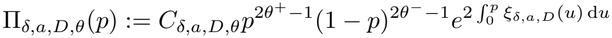

with *C*_*δ,a,D,θ*_ a normalization constant.

Finally, the definition of Δ^*\**^ lets us obtain the following fixed-point equation

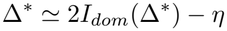

where

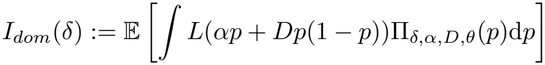

where the expectation is over (*α, D, θ*) taken to have the same law as 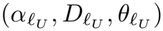 for*l*_*U*_ sampled uniformly at random over [*L*].

Overall, the philosophy is the same as in the additive case: we find the selection coefficient on locus*l* as a function of Δ_*t*_, approximate it with Δ^*\**^, use it to find the stationary distribution at a given locus and close the system by expressing Δ^*\**^ as a function of the stationary distribution at the typical locus. If *D* is of the same order as *α*∼ 1*/L*, then all the orders of magnitude from Table 1 remain accurate. The stationary distribution Π_*δ,a,D,θ*_ is much more complicated than previously (in particular, the exponential includes a polynomial of order 4 in *p*), but can be simplified under some circumstances. Most notably, if mutation rates are small 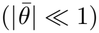 and selection is weak/moderate 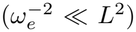 we actually recover the same stationary distribution as if there was no dominance and the same fixed-point equation. This means **dominance does not affect the stationary distribution, nor the distance to the optimum** (under small mutation rates and weak/moderate selection). Dominance will however impact the genetic variance.

#### 2.7.2 Epistasis

In SI 11 we discuss epistasis. In terms of model, this means the trait function *Z* may include interaction terms between locus*l* and locus*l*^*′*^ for*l*^*′*^ ≠*l*. We discuss two extreme cases for which our approach can be adapted to account for epistasis.

In the first case, the trait *Z*(*g*) is determined by many independent small modules. For instance, one could think of small independent gene regulatory networks such that mutations may affect local gene expression, but do not have long-range effect. Whereas in the additive case, stabilizing selection resulted in a fixed-point equation (15) emphasizing how a locus is influenced by the macroscopic observable Δ^*\**^, which itself emerges from the stationary distribution of the locus, here we have that each module is influenced by Δ^*\**^, which itself emerges from the multivariate distribution of allele frequencies within each module. So the general picture is that Δ^*\**^ is the only coupling factor between distinct regulatory networks.

In the second case, the trait *Z*(*g*) is determined by many independent interactions between pairs of loci. This is closer to physical models used in statistical physics such as the Sherrington-Kirkpatrick model [43]. The crucial idea is that each locus interacts loosely with all other loci to determine the trait. In this setting, due to the law of large numbers we may define some effective genetic effect at each locus, which depends on the law of the typical locus. The picture that emerges is then exactly the same as in the additive case, except that the genetic effect at a locus now also depends on the law of the typical locus. The fixed-point equation is then obtained by solving jointly for the effective effect and Δ^*\**^.

## 3 Discussion

### 3.1 The four features of the polygenic limit under Gaussian selection

We have discussed the polygenic limit *N, L*≫ 1 in a population under Gaussian selection. Under our assumptions (which guarantee HWLE and mean-field approximations) this population is then well described as follows

- sampling an organism at random at time *t*, its trait value is normally distributed with fixed variance *σ*^2^ and mean 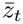.
- 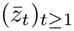 evolves as an Ornstein-Uhlenbeck process (21)
- a given locus*l* evolves independently of 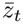 following the polygenic equation (12), parameterized by Δ^*\**^
- Δ^*\**^ is solution to the fixed point equation (15).

The first point (normality) has a long history in quantitative genetics, dating back to [66, 29]. It is deeply connected to the infinitesimal model [26, 4], which may or may not hold in our population (see Section 2.5). It has also played a crucial role in theoretical results, used as an assumption to derive predictions for the evolution of the population (e.g [6, 49, 75, 7]). Here, **we did not require this as an assumption**. Obviously, under our assumptions of HWLE, sufficient mutational input to maintain genetic variation and non-degenerate distribution of mutational effects, normality of a randomly sampled trait holds from the central limit theorem.

The second feature of the polygenic population (Ornstein-Uhlenbeck process for the trait mean) was famously suggested in [45], and its description in terms of genetic parameters was further developed in [18, 7, 49, 51] (in [7], what is obtained is close but distinct from an Ornstein-Uhlenbeck due to fluctuations of *N*). Our main contribution is to point out that under weak selection 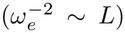, this description cannot hold in a strict mathematical sense (see Fig. S7 and SI 6.5), though it should be sufficient for most practical applications (provided the mutation rate is constant across loci).

The third and fourth points are our most important contributions. To the best of our knowledge, the fixed-point equation (15) had not previously been derived (though [18, 49, 51] came close under some extra assumptions such as constant allelic effects and mutational parameters across loci, and some restrictions on the strength of selection). Taking a different approach based on the stationary solution to the multi-locus

Wright-Fisher diffusion (5), [22] has recently found an explicit approximate expression for the solution Δ^*\**^ to the fixed-point equation (15) when the mutational bias is small (|*z*_*M*_ −*η*|≪ 1) and *θ*^+^ = *θ*−: this approximation is equivalent to computing the variance *V***ar**^*\**^[*P*_*t*_ |*α, θ*] by approximating the distribution of *P*_*t*_ with the neutral distribution Π_0,0,*θ*_ (the same idea is used in [18, 49, 51]). Using this idea coupled with the central limit theorem leads to the first-order perturbation to Π_0,0,*θ*_ due to selection.

### 3.2 The case for the gene-centric approach

A model to represent a polygenic trait in a large population is a complicated mathematical object, and analyzing its limit behavior necessarily requires some simplifying assumptions. Historically, a lot of emphasis has been put on the assumption that the trait remains normally distributed (e.g in [6, 49, 75, 7]). More generally, a very influential sequence of papers was [6, 5, 81], which set out a general framework to describe the evolution of the population by computing recursion equations for the cumulants. The goal of [6] is really to describe the evolution of macroscopic observables (the cumulants of the distribution of *Z*(*g*)) using recursion equations based on the cumulants of the distribution of allelic effects at a single locus (they use a continuum-of-alleles model which has since fallen from grace [73]). More generally, these papers write equations for the evolution of a cumulant of order *k* as a function of the cumulant of order *k* + 1. A crucial tool to do this is Wright’s formula [91], which relates the dynamics at a given locus to the gradient of the mean logfitness, thus expressing the effect of selection at the genetic level as a function of the trait distribution. To solve for equilibrium, it is then necessary to make a closure assumption: typically, one assumes cumulants of a certain order to be negligible (order 2 if the trait is normally distributed, sometimes order 3 to correct for skew).

On the other hand, the derivation we presented is more in line with gene-centric approaches such as deployed for instance in [12, 96, 79, 64, 57, 8] among others. At its core, the method can be summed up as follows:

- a diffusion approximation is used to describe the evolution of a typical locus which depends on microscopic parameters (the mutation probability *µ*, the allelic effect *α*) and also on an external variable, the bias-correcting coefficient *s*^*\**^
- a mean-field approximation is used to close the system by expressing Δ^*\**^ as a function of the microscopic observables, which determines the value of *s*^*\**^.

This leads to a fixed-point equation for the macroscopic observable, and the stationary distribution of the typical locus. We show in SI 7 that the gene’s-eye view and the traditional trait’s-eye view from [91] lead to the same equations. We discuss in SI 5.3-6 how to recover some classical approximations from our general equation (the stochastic house-of-cards approximation from [14, 38, 39], Lynch’s formula for Δ^*\**^ in [49]), showing that our approach is much more general and lets us recover these previous results as special cases under restrictive assumptions on the strength of selection 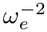 and the distribution of *α, θ*. We also note that [51] consider that the genetic variance *σ*^2^ is to a first order equal to its neutral value, (this assumption is also used locally in [18, 83]), which is only true strictly speaking in our setting under ultra-weak selection 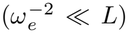.). This assumption would also hold if the mutational bias was small (*s*^*\**^ ≪1), which would require the mutational equilibrium *z*_*M*_ and the selection optimum *η* to be very close (this possibility is ruled out under our set of assumptions, see SI 2).

Our results can be further simplified if mutation rates are low 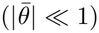. which corresponds to the infinite-sites model with reversible mutations in the terminology of [8]. We discuss this in SI 8, and in particular recover Charlesworth’s formula for *s*^*\**^ from [18].

This approach from the gene’s-eye view has seen increasing use in the past few years: for instance, [96] and [79] have used a similar approach to describe the stationary distribution of a population under directional selection and immigration. Another example is [9], in which the macroscopic observable is the genetic variance *σ*^2^ under fluctuating Gaussian selection, which determines the allelic frequencies, which in turn determine the genetic variance, thus yielding a fixed-point equation (their equation 16). In [8], a liability model is studied in which fitness is reduced by a fixed cost *C* when the polygenic trait *z* is greater than a threshold value: the selection coefficient at stationarity acting on an allele increasing liability by *α* then depends on the distribution of the population trait mean (their equation (8)), which is solved for in their equation (27).

The mathematical literature on mean-field approximations is vast and growing [16, 17, 37], and we believe that both the population genetics community and the probability community would benefit from a dialogue on this subject, for which we hope our contribution to be suited.

### 3.3 The three regimes of selection and the stabilizing transition

Our definition of three regimes of selection relies on the observation of the remarkable property that under moderate selection 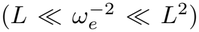, a single parameter *s*^*\**^ independent of 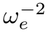 describes the law of 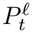(as was also seen in [18]). In this regime, a given locus only feels selection through the effect of the bias-correcting coefficient *s*^*\**^.

The regime of strong selection 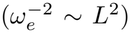 is the regime in which Robertson’s underdominant term kicks in, and selection actively depletes genetic variance. It is also the one in which the typical value of Δ_*t*_ (mean and fluctuations) is of the same order of magnitude as *α* (see Table 1), but the fluctuations of (Δ_*t*_)_*t≥*0_ are much faster. This presents an interesting mathematical challenge, since a rigorous analysis would have to prove convergence of 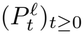 to the limit equation (12) not by neglecting the fluctuations of (Δ_*t*_)_*t≥*0_, as is typically the case in mean-field theory, but through a time-averaging theorem. We also recall in SI 5.5 how to recover the famous Stochastic House-of-Cards approximation [14, 38, 39] as a first-order expansion of *σ*^2^ in 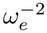, under the assumption that the (*α*_*l*_)_*l∈*[*L*]_ are distributed as the absolute value of a normal variable and that there is no mutational bias (*s*^*\**^ = 0). The original Stochastic House-of-Cards approximation is valid in a broader setting (it is derived from a continuum-of-alleles model in [14]), but its derivation in [38, 39] relies on a two-locus model similar to ours, and therefore only valid at the transition between moderate and strong selection.

Under weak selection, the system is maintained very far from the selection optimum (Δ^*\**^≫ *σ*, see Table 1), and the population only undergoes directional selection, effectively maintained at mutation-selection-drift bal-ance. The trait dynamics is not exactly given by an Ornstein-Uhlenbeck process, but can be approximated as such. In-between these two extremes, we identified the stabilizing transition, when the selection strength 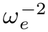 is of order 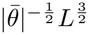 , as the only regime in which Δ and *σ* are of the same order. This transition corresponds to a switch between a regime in which the fitness load is mostly determined by directional selection (that is, Δ^*\**^) and a regime in which the fitness load is mostly determined by stabilizing selection (that is, *σ*). This transition is already informally discussed in [19] (after his equation (2b)). This transition is in particular important when considering the evolution of recombination, because in a population in which recombination rates are allowed to evolve, the directional component of the fitness load tends to favor more recombination whereas the stabilizing component of the fitness load tends to favor less recombination, all the way to asexuality [83].

### 3.4 On the importance of trait mutational bias

Trait mutational bias in a quantitative trait describes the situation in which the additive effect of mutations on the trait has a nonzero mean. This should not be confused with mutational bias in fitness. Mutational bias in fitness refers to the fact that if the population is close to the optimum, new mutations will tend to be deleterious (see for instance [83]). Even if there was no trait mutational bias (say *z*_*H*_ = *η* = *z*_*M*_), due to Robertson’s underdominant term in the selection coefficient (8), we would still have that, on average, a mutation with large effect *α* is deleterious. The impact of trait mutational bias on a polygenic trait has been described in [95] numerically, and [18] has provided a theoretical treatment which includes approximate solutions assuming low mutation rates 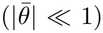 and constant (*θ*_*l*_, *α*_*l*_) across loci. [18] suggests two methods to approximate Δ^*\**^, one (Eq. A3.b) which coincides with our own (see our SI 5.3) and one which requires approximating E [*P*_*t*_(1 −*P*_*t*_) (*α, θ*)] with its neutral expectation (Eq. S3.3b). This second approximation was also used in [49, 51] to solve for Δ^*\**^, and in [83] to find the fitness load associated with mutational bias in sexual populations.

Apart from these exceptions, trait mutational bias is typically neglected in theoretical studies of polygenic traits under stabilizing selection. For instance, [67, 46, 12, 87, 10, 59, 75] all study a quantitative trait under stabilizing selection, and always assume that at equilibrium the mutation rate does not push the population in any specific direction away from the optimum ([75] consider anisotropic pleiotropic mutations as an extension of their base model, but still assume that the mean effect of any new mutation is zero). The models for quantitative traits from statistical physics such as [57, 59, 23] also typically assume symmetric mutation rates. This seems to be motivated not just by mathematical tractability, but by the idea that when the mutation rate is very low, the directional effect of mutations can be neglected. It would be unreasonable to argue that mutational bias is typically small, as this would imply that mutation favors the same trait values as selection, which would be a strange coincidence, in particular if this were satisfied across distinct environments. In fact, experimental data documenting mutational bias can be obtained from the evolution of morphological traits in mutation accumulation (MA) experiments. Morphological quantitative traits are typically assumed to be under stabilizing selection [73]. In MA lines, such traits generally evolve in a deterministic direction. For instance, table 2 of [30] reviews the case of drosophila, in which the proportion of trait-increasing mutations for sternopleural/abdominal bristle number and wing length range between 0.4 and 0.07. More recently, [53] has reported significant mutational bias in locomotion traits in *Caenorhabditis elegans*. Mutational bias could also account for cases in which directional selection and stabilizing selection are simultaneously observed on phenotypic data [71]. Population genetic analyses also suggest significant mutational bias for codon usage [74, 33, 94, 93]. Finally, [82] has established an empirical test of mutational bias for GWAS in humans, and successfully detected a bias for traits such as height, BMI, and educational attainment.

In the presence of mutational bias, which we expect to be pervasive, weak selection cannot be distinguished from directional selection [95], because the population is too far from the optimum 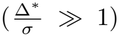. Such an equilibrium population would be in selection/mutation balance for a quantitative trait. While stable quantitative traits under directional selection are sometimes found in natural populations, mutational bias is however rarely mentioned as a putative explanation, presumably due to the relative strength of observed selection with respect to mutation (it is for instance never mentioned on reviews such as [42]). The possibility of mutations and genetic drift preventing a population from reaching the optimal trait value is discussed in the literature as the drift-barrier hypothesis. The typical quantitative trait for which this is deemed plausible is the mutation rate [78, 50] (but see also [48]). Our framework could be adapted to model this, following [21].

### 3.5 Example of a practical application: moderate selection and human height

One convenient aspect of our classification is that the qualitative behavior of the stationary system relies on parameters 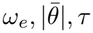 , which can be empirically measured. If we take the example of the human population, the effective population size is of order *N*_*e*_ 20, 000 [28]. For a trait like height, [76] estimate the mutational target to be of order 10^7^ ([92] obtained a saturated GWAS map for height from 12, 111 loci, so a certain lower bound for *L* is 10,000). The average effect of a locus is taken to be of order 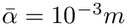, in line with previous estimates (the effect sizes vary between .03*cm* and 1*cm* according to [85]). In [71], the UK Biobank was used to estimate the parameter 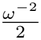, which was found to be of order 1.5 ×10^*−*2^ in units of *σ*^*−*2^, with *σ* of order 0.1*m*. Finally, we take the mean mutation rate 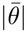 per locus per generation to be 10^*−*8^*N*_*e*_, where *N*_*e*_ is the effective population size [72]. If we neglect environmental variation for simplicity, we may compute

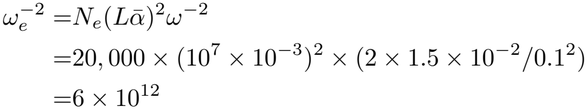

This is compatible with moderate selection, since 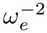 is much greater than *L* (of order 10^7^) but much smaller than *L*^2^ (of order 10^14^). Of course, applying this crude computation to human data assumes equilibrium and ignores population structure, linkage, the fluctuations of selection, the environmental contribution to height, pleiotropy and the fact that the allelic effects (*α*_*l*_)_*l∈*[*L*]_ have a heavy-tailed distribution.

### 3.6 Applicability to GWAS data

Our theoretical model yields a stationary distribution for the joint distribution of allelic frequency and effect (*P*_*t*_, *α*). This distribution is precisely what GWAS are measuring, and which the works of [82, 75, 76, 44] are interpreting from an evolutionary standpoint. Our contribution to their framework is the bias-correcting selection coefficient *s*^*\**^. From (8), we expect the distribution of *P*_*t*_ conditional on *α* to be proportional to Π[*ξ*_*α*_, *µ, N*_*e*_] where

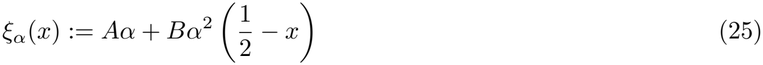

for some constants *A* and *B* for which we do have explicit theoretical expressions and which could be estimated empirically by maximum likelihood.

Empirically, [82] validates our core idea by detecting asymmetries in the distribution of (*P*_*t*_, *α*). More precisely, their model is concerned with (*Y*_*t*_, *β*) where *Y*_*t*_ is the frequency of the derived allele (as opposed to the ancestral allele) and *β* is its allelic effect. Compared to our model, *β* can be negative, but |*β* | = *α*. When *β <* 0, then *Y*_*t*_ = 1 −*P*_*t*_, otherwise *Y*_*t*_ = *P*_*t*_. They consistently reject the null hypothesis of unbiased mutations (same average value of *β* conditional on *Y*_*t*_ = *p* or on *Y*_*t*_ = 1 −*p*) in 6 out of 9 GWAS of human traits. One would expect this to mean our model (25) would fare well in practice.

It turns out that model (25) has been tested in the case of quantitative disease GWAS in [44]. In this preprint, the authors do not really justify model (25), but test it against the “traditional” model of [75] which corresponds to setting *A* = 0. Model (25) fares rather poorly, and for only 2 of 27 tested traits this model outperforms the *A* = 0 model in terms of restricted maximum likelihood (versus 21 traits that support the *A* = 0 model). We do note (their Fig. S13) that their model assuming only directional selection does yield consistent result, which suggests nevertheless an underlying asymmetry for many traits.

If it was confirmed that model (25) generally underperforms with respect to the model with *A* = 0, this could be interpreted in at least two different directions. First, we assume that genetic diversity is maintained by mutations. It can also be maintained by spatial structure [64]: if a very large population is subdivided into many small demes, then genetic diversity within each deme can be maintained by migration. In our model, if we assume a Levene migration model with identical fitness landscape in every deme [55] we could add this effect by replacing the mutation probabilities 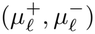 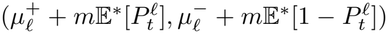 for some migration parameter 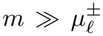. In such a setting, mutational bias should become negligible. Second, it could be seen as a signature of selection on the distribution (*α, θ*) itself. For instance, the mutation rate is highly variable across the genome [3], shaped by evolvable factors such as chromatin structure [52]. The distribution of *α* can be shaped through second-order selection favoring robust gene regulatory networks. A quantitative trait under Gaussian selection can evolve towards robustness in a stable environment, because selection favors genes which reduces the genetic variance in the trait [88]. The process by which Gaussian selection shapes the distribution of *α* is called genetic canalization [27] (see also [31]).

A framework for a systematic evolutionary analysis GWAS analysis was recently proposed [76] (currently in revision). The model considers that each locus has a highly pleiotropic effect, which determines its selection coefficient in Robertson’s underdominant term (the pleiotropic equivalent of the second term of (8), see SI 10.2). It is assumed that conditional on this selection coefficient, the additive effect of the alleles at this locus on a given focal trait has a symmetric distribution. This assumption posits that selection and mutation act identically on trait-increasing and trait-decreasing alleles, implying that *z*_*M*_ = *z*_*H*_ = *η* and that the distributions of (*α, P*_*t*_) and (*α*, 1 −*P*_*t*_) are identical. We would argue for a generalization of this approach including a possible bias in mutation and in allelic effects, for example through the addition of a hyperparameter.

### 3.7 Future directions

On top of all extensions discussed in Section 2.7, it will also be straightforward to add fluctuating environments as in [9] (provided the fluctuations are of the same order as *σ* or smaller), genes influencing plasticity (G× E interactions, which are studied from the trait’s-eye view in [84]), spatial structure as in [64], and more sophisticated selection (for instance if *W* (*z*) is skewed).

One could also consider would be insertions-deletions, which would let the number of loci *L* evolve [63, 62, 20, 65, 47]. One could also follow the spirit of [88], modeling canalization by adding modifier loci altering the genetic effects (*α*_*l*_)_*l∈*[*L*]_. On another level, there is growing evidence for the rôle of introgression and admixture [24] as an alternative to mutation for generating genetic diversity. Theoretical models such as [70] offer powerful descriptions of the long-term effect of an admixture event on a polygenic trait. These models allow for the representation of migration as an input in genetic variability under assumptions of strong recombination and weak selection [69]. Finally, one should account for the evolution of the effective population size *N*_*e*_, the fluctuations of which can be influenced directly by the effect of selection [7, 64, 80].

Future work will also describe the dynamics of the system out-of-equilibrium, extending the work of [32, 2, 54]. This will allow us to tackle questions such as the response of a population to a decrease in population size or a change in optimum, and the corresponding genetic load [19]. Overall, our framework lets us account for most biologically relevant features of polygenic systems (with the exception of LD) and provides efficient theoretical predictions as well as explicit criteria on the parameters for our predictions to hold.

## 4 Data availability

Our programs were run on Python 3.10.12. All of our code is available at the following adress: https://doi.org/10.5281/zenodo.18877448.

### Supporting Information

All of our supporting information is organised as a single file, which is structured as follows.

1. **Model and notation**
2. **Outline of the derivation**
3. **The diffusion approximation**
4. **The polygenic equation from the gene’s eye-view**
5. **Observables and scalings**
6. **Breakdown of the polygenic limit**
7. **Equivalence with the trait’s eye-view**
8. **When mutations are rare at a given locus**
9. **Derivation of the criterion for selection to increase genetic variance**.
10. **Some extensions**
11. **Epistasis**
12. **Numerical methods**

## 5 Acknowledgments

We hold deep gratitude towards one anonymous reviewer, Joshua Schraiber and John Welch for a most helpful review, for their comments and references. We thank Carsten Fortmann-Grote for data reviewing and helpful coding suggestions.

We thank Paul Jenkins and Henrique Teotónio for helpful comments and references. We thank Archana Devi, Mathilde André and François Ged for fruitful discussions. Thanks also to Céline Teplitsky who first introduced P. Courau to quantitative genetics.

## 8 Use of Large Language Models (LLM)

The authors declare no LLM was used at any point in the preparation of this manuscript.

