## Supplementary material for "Gaussian selection on a polygenic trait from the gene’s-eye view"

September 2026

1. Institut de Biologie de l'ENS (IBENS), École Normale Supérieure, Paris, France 75005

2. Centre Interdisciplinaire de Recherche en Biologie (CIRB), Collège de France, Paris, France 75005

3. Department of Statistics, University of Vienna, Vienna, Austria 1090

∗ Corresponding author: philibert.couraugmail.com.

#### 1 Model and notation

To make this appendix self-contained, we recall the model and notation from the main text.

##### 1.1 Miscellaneous notations

For $a, b \in \mathbb{R}_+$, the notation $a \lesssim b$ is taken to mean that we don't have $a \gg b$, and $a \sim b$ means we neither have $a \ll b$ nor $b \ll a$. We use the notation $[L] = \{1, \dots, L\}$.

##### 1.2 Individual-based model

The individual-based model as implemented in our numerical simulation is a classical diploid $L-$loci biallelic Wright-Fisher model with a population of size $N$. Each organism can be described by its **genome** $g = (g_\ell)_{\ell\in[L]} \in \{0, 1, 2\}^L$, with $g_\ell$ representing the number of trait-increasing allele at locus $\ell$. The (genetic) trait value of a genome is given by the **trait function**

$$Z : \begin{cases} \{0,1,2\}^L & \longrightarrow \mathbb{R} \\ g & \longmapsto \sum_{\ell\in[L]} \alpha_\ell g_\ell \end{cases}$$

where $\alpha_\ell \geq 0$ is the genetic coefficient at locus $\ell$. We take the $(\alpha_\ell)_{\ell\in[L]}$ to be an exchangeable vector of random variables such that

$$\sum_{\ell\in[L]} \alpha_\ell = 1. \tag{1}$$

In particular, unless their distribution is degenerate (see Figure 6) the typical value of $\alpha_\ell$ is of order $1/L$. The assumption of (1) is made to guarantee that $Z(g)$ is always between 0 and 2. It can be seen as a form of rescaling: we measure the trait in units such that the maximal possible trait value is 2 and the smallest one is 0.

The **fitness** of an organism is obtained with the **fitness function**

$$F(z) : \begin{cases} \mathbb{R} & \longrightarrow \mathbb{R} \\ z & \longmapsto \exp\left(-\frac{\omega^{-2}}{2}(z-\eta)^2\right) \end{cases}$$

We let $W := \ln F$ be the **log-fitness** function.

Reproduction occurs by sampling two parents with probability proportional to fitness, and for each parent picking a crossover position uniformly on $[L]$. Mutations at locus $\ell$ to (resp. from) the trait-increasing allele occur with probability $\mu_\ell^+$ (resp. $\mu_\ell^-$) per locus per generation per haploid genome. A formal definition of the model will be given in Section 3.

#### 1.3 Representation from allelic frequencies

For $t \geq 0$ we let $P_t^\ell$ be the frequency of the trait-increasing allele at generation $\lfloor 2Nt \rfloor$ at locus $\ell$.

For $\mathbf{p} = (p_\ell)_{\ell\in[L]} \in [0,1]^L$ we let $\mathbf{E_p}[\varphi(g)]$ be the expectation of $\varphi(g)$ when $g = (g_\ell)_{\ell\in[L]}$ is a vector of independent variables such that $g_\ell$ has law $Binomial(2, p_\ell)$. We write $\mathbf{Var_p}, \mathbf{Cov_p}$ for the variance and covariance associated to $\mathbf{E_p}$. For all other random variables the notation $\mathbb{E}, \mathbb{V}\mathbf{ar}$ and $\mathbb{C}\mathbf{ov}$ are used.

Consider $t \geq 0$. The population at time $t$ is in **Hardy-Weinberg Linkage Equilibrium (HWLE)** if conditional on $\mathbf{P}_t = (P_t^\ell)_{\ell\in[L]}$, the law of of a uniformly sampled genome is $\mathbf{E}_{\mathbf{P}_t}$.

#### 1.4 Typical locus and genetic architecture

For $\ell \in [L]$, we define the summary vector at locus $\ell$ at time $t$ as the vector

$$\vec{P}_t^\ell := (P_t^\ell, \alpha_\ell, \theta_\ell).$$

This vector contains all the information we need from locus $\ell$ at time $t$. Consider $\ell_U$ a uniform random variable on $[L]$. We call

$$\vec{P}_t = (P_t, \alpha, \theta) := \vec{P}_t^{\ell_U}$$

the **typical locus at time** $t$. The **genetic architecture** at time $t$ is the law of $\vec{P}_t$.

#### 1.5 Wright-Fisher diffusion

A process $(P_t)_{t\geq 0}$ is a standard Wright-Fisher diffusion with parameters $s > 0, \theta^+ > 0, \theta^- > 0$ if it satisfies the Stochastic Differential Equation (SDE)

$$\mathrm{d}P_t = sP_t(1-P_t)\mathrm{d}t + (\theta^+(1-P_t) - \theta^- P_t)\mathrm{d}t + \sqrt{P_t(1-P_t)}\mathrm{d}B_t \tag{2}$$

where $B_t$ is a Brownian motion.

Similarly, a process $(P_t)_{t\geq 0}$ is a frequency-dependent Wright-Fisher diffusion with parameters $\xi, \theta^+, \theta^-$ where $\xi$ is a continuous function on $[0,1]$ and $\theta^+ > 0, \theta^- > 0$ if it satisfies the SDE

$$\mathrm{d}P_t = \xi(P_t)P_t(1-P_t)\mathrm{d}t + (\theta^+(1-P_t) - \theta^- P_t)\mathrm{d}t + \sqrt{P_t(1-P_t)}\mathrm{d}B_t \tag{3}$$

### 2 Outline of the derivation

We outline the content of Sections 3-6.

#### 2.1 The polygenic equation from the gene's eye view

In Sections 3-4, we will derive an equation that jointly describes the evolution of the typical locus $P_t$ and the mean trait value within the population. This will be done in two successive steps.

##### 2.1.1 The diffusion equation

Our first step relies on deriving coupled diffusion equations for the evolutions of the allelic frequencies at various loci. Define $\Delta_t$ as the deviation from the optimum at time $t$

$$\Delta_t := \mathbf{E}_{\mathbf{P}_t}[Z(g)] - \eta = \sum_{\ell\in[L]} 2\alpha_\ell P_t^\ell \; - \eta. \tag{4}$$

In Section 3, we derive the following diffusion equation for $P_t^\ell$ on the time scale $N$

$$\mathrm{d}P_t^\ell = \xi_{\Delta_t,\alpha_\ell}(P_t^\ell)P_t^\ell(1-P_t^\ell)\,\mathrm{d}t + \;\left(\theta_\ell^+(1-P_t^\ell) - \theta_\ell^- P_t^\ell\right)\mathrm{d}t + \sqrt{P_t^\ell(1-P_t^\ell)}\,\mathrm{d}B_t^\ell, \tag{5}$$

where $(B^\ell)_{\ell\in[L]}$ are independent Brownian motions, $\theta_\ell^\pm = 2N\mu_\ell^\pm$, and the selection coefficient $\xi$ is such that for $a \in \mathbb{R}_+, \delta \in \mathbb{R}$:

$$\xi_{\delta,a}(p) := -\frac{a\delta}{\omega_e^2} + \frac{a^2}{\omega_e^2}\left(p - \frac{1}{2}\right). \tag{6}$$

where

$$\omega_e^2 := \frac{\omega^2}{2N}.$$

The derivation of of this result relies on the following hypotheses

**H 1.** *The population remains close to HWLE.*

**H 2.** *The law of $F(Z(g))$ when $g$ has law $\mathbf{E}_{\mathbf{P}_t}$ is very concentrated around $\hat{F}_t := e^{\bar{W}_t}$ where $\bar{W}_t = \mathbf{E}_{\mathbf{P}_t}[Z(g)]$, so that we may write*

$$F(Z(g)) \simeq \hat{F}_t(1 + W(Z(g)) - \bar{W}_t).$$

(H1) implies the $(P_t^\ell)_{\ell \in [L]}$ are sufficient to fully describe the population at time $t$. (H2) should be interpreted as: the typical fitness difference between two randomly sampled organisms is small.

###### 2.1.2 Mean-field approximation

From now on, we simplify the problem by considering that the system is at statistical equilibrium, and we denote by $\mathbb{P}^*$ the equilibrium distribution of the typical locus $\vec{P}_t$. The diffusion (5) is of dimension $L-1$, which makes it inconvenient to work with when $L \gg 1$. In Section 4, we derive a simplified system for the dynamics of the typical locus from (5) using mean-field approximations.

Mean-field approximations are an important tool in the study of interacting particles [10, 11]. In our setting, the mean-field approximation reads

**H 3** (Mean-field approximations)**.** *For any test function $f$,*

$$\frac{1}{L}\sum_{\ell \in [L]} f(\vec{P}_t^\ell) \simeq \mathbb{E}^*[f(\vec{P}_t)] = \mathbb{E}^*[f(\vec{P}_0)].$$

*where the last equality follows from the fact that the system is assumed to be at equilibrium.*

The left hand side is a random quantity, whereas the right-hand side is deterministic and is obtained by looking at the expected value of an observable of a typical locus at time 0 (under the equilibrium distribution $\mathbb{P}^*$).

Intuitively, in our system, the loci are coupled by $\Delta_t$ in the selection coefficient (6), and $\Delta_t$ is determined by all loci. But this dependence is diffuse and evenly spread across all loci, as each locus only has a very small influence on $\Delta_t$. So the dependence between different loci is weak, and we may expect (H3) to hold. This phenomenon, known as **propagation of chaos** [10], has been rigorously proven to occur in our system when $\omega_e^{-2} \sim L$ in [14].

In Section 4, we formally derive the limit equation for the distance of the trait mean to the optimum and a typical locus $P_t$ at stationarity, and the fluctuations of the mean trait value $\varepsilon_t := \Delta_t - \Delta^*$.

$$\mathrm{d}P_t = \xi_{\Delta^*,\alpha}(P_t)P_t(1-P_t)\,\mathrm{d}t + \left(\theta^+(1-P_t) - \theta^- P_t\right)\mathrm{d}t + \sqrt{P_t(1-P_t)}\,\mathrm{d}B_t^P \tag{7}$$
$$\Delta^* := \mathbb{E}^*[\Delta_0] = \mathbb{E}^*[\Delta_t] = 2L\mathbb{E}^*[\alpha P_t] - \eta \tag{8}$$
$$\mathrm{d}\varepsilon_t = -\rho\varepsilon_t\,\mathrm{d}t + \omega_e\sqrt{2\rho}\,\mathrm{d}B_t^\Delta \tag{9}$$

where $B^P, B^\Delta$ are Brownian motions. In particular, $(\varepsilon_t)_{t\geq 0}$ is an Ornstein-Uhlenbeck process with **autocorrelation parameter** $\rho$ defined as

$$\rho := \frac{1}{\tau} \times \frac{\mathbb{E}^*[2(L\alpha)^2 P_t(1-P_t)]}{|\bar{\theta}|} \tag{10}$$

where the **characteristic timescale of the trait** $\tau$ are

$$\tau := \frac{L\omega_e^2}{|\bar{\theta}|}. \tag{11}$$

where $|\theta| := \theta^+ + \theta^-$ and $\bar{\theta} = (\mathbb{E}[\theta^+], \mathbb{E}[\theta^-])$. This system is much more convenient than the SDE (5), and in particular it has a simple stationary distribution (see Section 2.1.3).

The derivation will be based on the mean-field hypothesis (H3) and either one of the following two hypotheses

**H 4.** *We have $\omega_e^{-2} \ll L^2$.*

**H 4'.** *We have*

*a)* $\omega_e^{-2} \sim L^2$

*b)* $\mathbb{E}^*\left[(L\alpha)^2 P_t(1-P_t)\right] \sim |\bar{\theta}|$

*c)* $L\omega_e^2 \ll |\bar{\theta}|$

*d) the timescale on which $(P_t)_{t\geq 0}$ evolves is of order 1 or greater*

(H4) corresponds to what we call the weak/moderate selection regime (see main text). (H4'a) corresponds to the strong selection regime.

(H4'b) is a classical result of population genetics. If we take $\hat{P}_t$ a neutral Wright-Fisher diffusion with mutation rates $\theta$, then at stationarity $\hat{P}_t$ has distribution $Beta(2\theta^+, 2\theta^-)$, and in particular

$$\mathbb{E}^*[\hat{P}_t(1-\hat{P}_t)] = \frac{\theta^+\theta^-}{|\theta|\left(\frac{1}{2}+|\theta|\right)}$$

In particular, provided $\theta^+ \sim \theta^- \lesssim 1$, we have

$$\mathbb{E}^*[\hat{P}_t(1-\hat{P}_t)] \sim |\theta|$$

The same holds if $\hat{P}_t$ is a frequency-dependent Wright-Fisher diffusion (3) with selection coefficient of order 1. In light of this, we may expect (H4'b) to hold as soon as $\alpha \sim 1/L$ and the mutational bias is small. See Section 5 for further details.

(H4'c) can be rewritten $\tau \ll 1$. (H4'd) is verified for any Wright-Fisher diffusion in which genetic drift plays a significant rôle.

##### 2.1.3 Stationary distribution

Using the decoupling of $P_t$ and $\Delta_t$, we can rewrite (7) as

$$\mathrm{d}P_t^\ell = \xi_{\Delta^*,\alpha_\ell}(P_t^\ell)P_t^\ell(1-P_t^\ell)\,\mathrm{d}t + \left(\theta_\ell^+(1-P_t) - \theta_\ell^- P_t\right)\mathrm{d}t + \sqrt{P_t^\ell(1-P_t^\ell)}\,\mathrm{d}B_t^\ell \tag{12}$$

This means that conditional on $(\alpha_\ell, \theta_\ell)$, $(P_t^\ell)_{t\geq 0}$ behaves as a frequency-dependent Wright-Fisher diffusion (3). In particular, its equilibrium distribution [25] is $\Pi_{\Delta^*,\alpha_\ell,\theta_\ell}$ where for $\delta \in \mathbb{R}, a \in \mathbb{R}_+, \theta \in (0,+\infty)^2$ we define

$$\Pi_{\delta,a,\theta}(p) := C_{\delta,a,\theta}\, p^{2\theta^+-1}(1-p)^{2\theta^--1}e^{2\int_0^p \xi_{\delta,a}(u)\,\mathrm{d}u} = C_{\delta,a,\theta}\, p^{2\theta^+-1}(1-p)^{2\theta^--1}e^{-2-\frac{\Delta^*}{\omega_e^2}ap-\frac{a^2}{\omega_e^2}p(1-p)}, \tag{13}$$

with $C_{\delta,a,\theta}$ a normalization constant. Furthermore, the distribution of a typical locus $\vec{P}_t$ is given by

$$\mathbb{E}^*[f(\vec{P}_t)] = \mathbb{E}\left[\int f(p,\alpha,\theta)\Pi_{\Delta^*,\alpha,\theta}(p)\mathrm{d}p\right]$$

Since we know the distribution of a typical locus, we can compute

$$\Delta^* \simeq 2L\mathbb{E}^*[\alpha P_t] - \eta.$$

This can be rewritten

$$\Delta^* \simeq 2\, I(\Delta^*) - \eta \tag{14}$$

with

$$I(\delta) := \mathbb{E}\left[\int L\alpha p \Pi_{\delta,\alpha,\theta}(p)\mathrm{d}p\right]. \tag{15}$$

With words, $I(\delta)$ is the expectation of $L\alpha\hat{P}_t^\delta$, where $\hat{P}_t^\delta$ is a stationary frequency-dependent Wright-Fisher diffusion with parameters $(\xi_{\delta,\alpha}, \theta)$ ((3)).

In mathematical terms, (14) is what is known as a fixed point equation: we obtain $\Delta^*$ as a function of $\Delta^*$. The fact that this equation has a unique solution can be seen from the fact that $I$ is continuous, non-increasing and bounded by 1 on $\mathbb{R}$.

#### 2.2 Simplifying assumptions on the parameters

In Section 5, we consider the stationary solution of the system given by Eq. (7-9), and we derive the order of magnitude of macroscopic observables based on the following assumptions.

**A 1** (Uniform boundedness). *There is a constant* $C \sim 1$ *such that for any* $\ell \in [L]$, $|\theta_\ell| \leq C|\bar{\theta}|$ *and* $\alpha_\ell \leq \frac{C}{L}$.

**A 2** (Mutations smaller than genetic drift). $|\bar{\theta}| \lesssim 1$.

**A 3** (Mutational bias not too extreme). *There is a constant* $C \sim 1$ *such that for any* $\ell$, $\theta_\ell^-/C \leq \theta_\ell^+ \leq C\theta_\ell^-$.

**A 4** (Accessibility of the selection optimum). $\eta \in (0,2)$ *satisfies* $\eta(2-\eta) \sim 1$.

**A 5** (Weak/moderate/strong selection). $L \lesssim \omega_e^{-2} \lesssim L^2$.

**A 6** (Distance between the selection and the mutation optimum). *We have*

$$|2\, I(0) - \eta| \sim 1.$$

(A1) is a strong assumption that guarantees no single locus disproportionally contributes to the genetic variance. It makes computations much more tractable. Future work should relax this assumption (see also Fig. 6). (A2-3) ensure mutations rates are not too large or too asymmetric. In particular, we allow for small mutation rates ($|\bar{\theta}| \ll 1$), which corresponds to the infinite-sites model as used in [6]. (A4) ensures that there are many different genotypes that can satisfy $Z(g) = \eta$. (A5) rules out ultra-weak ($\omega_e^{-2} \ll L$) selection, under which the system is close to neutrality (see Section 5.4), and ultra-strong $\omega_e^{-2} \gg L^2$, under which natural selection strongly depletes the genetic variability. (A6) is quite technical, but when $\omega_e^{-2} \ll L^2$ (weak/moderate selection) it can be rewritten $|z_M - \eta| \sim 1$ where $z_M := 2L\mathbb{E}\left[\alpha \frac{\theta^+}{\theta^+ + \theta^-}\right]$ is the mutational equilibrium (the mean trait value if there was no selection). In such a setting, (A6) precludes coincidental situations in which the mutational equilibrium would be very close to the selection optimum. Under strong selection ($\omega_e^{-2} \sim L^2$), the condition is more complex, but it similarly rules out a very coincidental situation.

##### 2.3 Necessary assumptions for consistency

In Section 6, we will show that the following assumptions are necessary for (A1-6) to be consistent with (H1-4').

**N 1** (Sufficiently large population). $2N \gg |\bar{\theta}| \frac{\ln(L)}{L\omega_e^2} + L\sqrt{|\bar{\theta}|}$.

**N 2** (Minimal mutational input every generation). $|\bar{\theta}| L \gg 1$.

Furthermore, the following assumption (stronger than (N2)) is needed for (9) to describe the fluctuations of the distance to the optimum $(\varepsilon_t)_{t\geq 0}$.

**N 3** (Sufficient mutational input). $\frac{|\bar{\theta}|}{L\omega_e^2} \gg 1$.

Condition (N2) guarantees that sufficient genetic variability within the population is maintained by mutations. (N1) ensures HWLE (H1) and that the fitness variance is small (H2). Finally, (N3) guarantees that selection is the force stabilizing the fluctuations of $(\varepsilon_t)_{t\geq 0}$ (see Section 6 for details), it can be rewritten as $\tau \ll 1$ with $\tau$ the timescale of the trait from (11).

#### 3 The diffusion approximation

In this section, we aim at obtaining the SDE (5), assuming $N \gg 1$, $\theta_\ell^\pm = \mu_\ell^\pm 2N \sim 1$, HWLE (H1) and that fitness is very concentrated (H2). Equation (5) is well-known to be the limit of the individual-based model under the hypothesis of HWLE as $N \to +\infty$, when selection and mutation are weak (see [1]), and the selection coefficient (6) was obtained by Wright in [47]. We recall the derivation of these equations in an effort to be self-contained.

We will obtain the diffusion equation by computing the first and second moments. Specifically, we must show

$$\mathbb{E}\left[P^\ell_{t+\frac{1}{2N}} - P^\ell_t \,\middle|\, \mathbf{P}_t\right] \simeq \frac{1}{2N}\left(\xi_{\Delta_t,\alpha_\ell}(P^\ell_t)P^\ell_t(1-P^\ell_t) + \theta^+_\ell(1-P^\ell_t) - \theta^-_\ell P^\ell_t\right) \tag{16}$$

$$\mathbb{V}\mathbf{ar}\left[P^\ell_{t+\frac{1}{2N}} \,\middle|\, \mathbf{P}_t\right] \simeq \frac{1}{2N} P^\ell_t(1-P^\ell_t) \tag{17}$$

$$\left|\mathbb{C}\mathbf{ov}\left[P^{\ell_1}_{t+\frac{1}{2N}}, P^{\ell_2}_{t+\frac{1}{2N}} \,\middle|\, \mathbf{P}_t\right]\right| \ll \frac{1}{2N} \tag{18}$$

##### 3.1 First moment computation

We write $P^{*\ell}_t$ for the frequency of the trait-increasing allele at generation $\lfloor 2Nt \rfloor$ after reproduction but before mutation. Assume the population at time $2Nt$ is in HWLE (H1). Because a genome $g$ from generation $\lfloor 2Nt \rfloor$ has an average number of offspring proportional to $F(Z(g))$, and because $P^{*\ell}_t$ is half the expectation of $g_\ell$ post-reproduction, we find

$$\mathbb{E}\left[P^{*\ell}_t \,|\mathbf{P}_t\right] = \frac{\mathbf{E}_{\mathbf{P}_t}[\frac{g_\ell}{2}F(Z(g))]}{\mathbf{E}_{\mathbf{P}_t}[F(Z(g))]}.$$

It follows from $P^\ell_t = \frac{1}{2}\mathbf{E}_{\mathbf{P}_t}[g_\ell]$

$$\mathbb{E}\left[P^{*\ell}_t - P^\ell_t \,|\mathbf{P}_t\right] = \frac{\mathbf{Cov}_{\mathbf{P}_t}[F(Z(g)), g_\ell]}{2\mathbf{E}_{\mathbf{P}_t}[F(Z(g))]}.$$

As a sidenote, this can also be seen as an application of the Price equation [37] to the trait $\frac{g_\ell}{2}$. Writing $\hat{F}_t = e^{\bar{W}_t}$ we get from (H2)

$$\frac{\mathbf{Cov}_{\mathbf{P}_t}[F(Z(g)), g_\ell]}{2\mathbf{E}_{\mathbf{P}_t}[F(Z(g))]} \simeq \frac{\mathbf{Cov}_{\mathbf{P}_t}[\hat{F}_t(1 + W(Z(g)) - \bar{W}_t), g_\ell]}{2\hat{F}_t} = \frac{\mathbf{Cov}_{\mathbf{P}_t}[W(Z(g)), g_\ell]}{2}$$

Using $\mathbf{Var}_{\mathbf{P}_t}[g_\ell] = 2P_t^\ell(1 - P_t^\ell)$ we find

$$\mathbb{E}\left[P_t^{*\ell} - P_t^\ell \,|\mathbf{P}_t\right] \simeq \frac{s_\ell(\mathbf{P}_t)}{2N} P_t^\ell(1 - P_t^\ell)$$

where $s_\ell$ is the selection coefficient at locus $\ell$

$$s_\ell : \begin{cases} [0,1]^L & \longrightarrow \mathbb{R} \\ \mathbf{p} & \longmapsto 2N\frac{\mathbf{Cov}_{\mathbf{p}}[W(Z(g)), g_\ell]}{\mathbf{Var}_{\mathbf{p}}[g_\ell]} \end{cases} \tag{19}$$

In particular, $\frac{s_\ell}{2N}$ can be seen as the linear regression of $W(Z(g))$ on $g_\ell$ (see Fig. 1).

Taking into account the effect of mutation we find

$$\mathbb{E}\left[P^\ell_{t+\frac{1}{2N}} - P_t^\ell \middle|\, \mathbf{P}_t\right] \simeq \frac{1}{2N}\left(s_\ell(\mathbf{P}_t)P_t^\ell(1 - P_t^\ell) + \theta_\ell^+(1 - P_t^\ell) - \theta_\ell^- P_t^\ell\right)$$

To get (16), we must show $s_\ell(\mathbf{P}_t) = \xi_{\Delta_t,\alpha_\ell}(P_t^\ell)$.

#### 3.2 Selection coefficient

We find from the definition of $s_\ell$ and $W$

$$\begin{aligned}
\frac{s_\ell(\mathbf{p})}{2N} =& \frac{\mathbf{Cov}_{\mathbf{p}}\left[-\frac{1}{2\omega^2}\left(\sum_{\ell'\in[L]} \alpha_{\ell'} g_{\ell'} - \eta\right)^2, g_\ell\right]}{2p_\ell(1 - p_\ell)} \\
=& -\frac{1}{4\omega^2 p_\ell(1 - p_\ell)}\left(\mathbf{Cov}_{\mathbf{p}}\left[\sum_{\ell_1,\ell_2\in[L]} \alpha_{\ell_1} g_{\ell_1} \alpha_{\ell_2} g_{\ell_2}, g_\ell\right] - 2\eta\alpha_\ell \mathbf{Var}_{\mathbf{p}}\left[g_\ell\right]\right) \\
=& -\frac{1}{4\omega^2 p_\ell(1 - p_\ell)}\left(\mathbf{Cov}_{\mathbf{p}}\left[\sum_{\substack{\ell'\in[L] \\ \ell'\neq\ell}} 2\alpha_{\ell'} g_{\ell'} \alpha_\ell g_\ell, g_\ell\right] + \mathbf{Cov}_{\mathbf{p}}\left[(\alpha_\ell g_\ell)^2, g_\ell\right] - 4\eta\alpha_\ell p_\ell(1 - p_\ell)\right)
\end{aligned}$$

where in the last equality we used (H1) which guarantees the independence of $g_{\ell_1} g_{\ell_2}$ and $g_\ell$ when $\ell \notin \{\ell_1, \ell_2\}$. This independence further yields for $\ell' \neq \ell$

$$\begin{aligned}
\mathbf{Cov}_{\mathbf{p}}\left[\alpha_{\ell'} g_{\ell'} \alpha_\ell g_\ell, g_\ell\right] =& \alpha_{\ell'} \mathbf{E}_{\mathbf{p}}\left[g_{\ell'}\right] \alpha_\ell \mathbf{Var}_{\mathbf{p}}[g_\ell] \\
=& \alpha_{\ell'} \times 2p_{\ell'} \times \alpha_\ell \times 2p_\ell(1 - p_\ell)
\end{aligned}$$

Similarly we can compute

$$\mathbf{Cov}_{\mathbf{p}}\left[\left(\alpha_\ell g_\ell\right)^2, g_\ell\right] = \alpha_\ell^2 \left(2 + 4p_\ell\right) p_\ell(1 - p_\ell)$$

From there it is elementary to obtain

$$\frac{s_\ell(\mathbf{p})}{2N} = \frac{\alpha_\ell}{\omega^2}\left(\eta - 2\sum_{\ell'\in[L]} \alpha_{\ell'}\, p_{\ell'}\right) + \frac{\alpha_\ell^2}{\omega^2}\left(p_\ell - \frac{1}{2}\right)$$

We thus get (6) using (4).

#### 3.3 Second moment

Here we compute $\mathbb{V}\mathbf{ar}\left[P^\ell_{t+\frac{1}{2N}} \,\middle|\, \mathbf{P}_t\right]$ and $\mathbb{C}\mathbf{ov}\left[P^{\ell_1}_{t+\frac{1}{2N}}, P^{\ell_2}_{t+\frac{1}{2N}} \,\middle|\, \mathbf{P}_t\right]$ for $\ell_1 \neq \ell_2$. For this, it will be useful to give a more formal definition of the individual-based model.

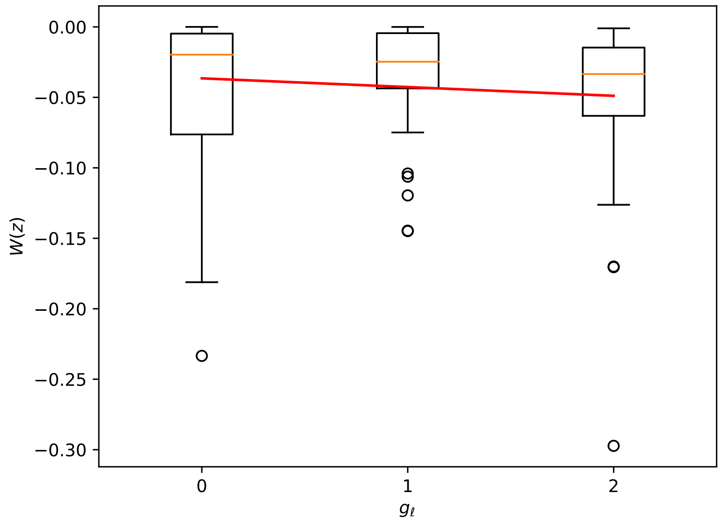


Figure 1: A population of $N = 100$ individuals with $L = 100$ biallelic loci, was evolved for $T = 100$ generations under Gaussian selection with parameters $\eta = 1.2, \omega_e^{-2} = 10^3$ and $\theta = (0.1, 0.2)$. The logfitness of the organisms within the population is plotted as a function of $g_\ell$ for $\ell = 3$. The selection coefficient $s_\ell(\mathbf{P}_t)$ at generation $t$ at locus $\ell$ is given by the linear regression coefficient of logfitness $W(Z(g))$ on $g_\ell$ (see Section 1A). This corresponds to the slope of the red line.

##### 3.3.1 Formal definition of the individual-based model.

The population is described with an array $(G^{n,i}_{\ell,(j)})_{\ell\in[L],j\in[2],i\in[N]}$ in which

- $n \in \mathbb{N}$ is a time-coordinate, it denotes the generation under consideration
- $i \in [N]$ is the label of the organism under consideration
- $j \in [2]$ is the label of the chromosome
- $\ell \in [L]$ is the label of the locus.

In particular, we have $G^{n,i}_{\ell,(j)} = 1$ (resp. 0) if the $i-$th organism at generation $n$ has the trait-increasing (resp. decreasing) allele on its $j-$th chromosome, at locus $\ell$. Then $g^{n,i} = (g^{n,i}_\ell)_{\ell\in[L]} := (G^{n,i}_{\ell,(1)} + G^{n,i}_{\ell,(2)})_{\ell\in[L]}$ is the genome of organism $i$ at generation $n$.

The generation $n+1$ is generated from generation $n$ as follows. For $i \in [N], j \in [2]$, the $j-$th chromosome of the $i-$th organism $(G^{n+1,i}_{\ell,(j)})_{\ell\in[L]}$ is independently generated in two steps

- *(Reproduction).* we sample a parental genome $I_{i,j}$ with probability proportional to $F(Z(g^{n,I_{i,j}}))$. A crossover positions $\ell_U^{(j)}$ is uniformly sampled on $[L]$, and a *Bernoulli*$(1/2)$ variable $b_j$. Then we set

$$\forall \ell \in [L], \qquad G^{n+1,i}_{\ell,(j)} = \begin{cases} G^{n,I_{i,j}}_{\ell,(b_j)} & \text{if } \ell < \ell_U^{(j)} \\ G^{n,I_{i,j}}_{\ell,(1-b_j)} & \text{otherwise} \end{cases}$$

- *(Mutation).* With probability $|\mu_\ell| := \mu_\ell^+ + \mu_\ell^-$, $G^{n,i}_{\ell,(j)}$ mutates and is replaced by an independently sampled variable with law *Bernoulli*$(\mu_\ell^+/(\mu_\ell^+ + \mu_\ell^-))$.

##### 3.3.2 Second moment: diagonal coefficients.

From the formal model, we see that each $(G^{\lfloor 2Nt\rfloor+1,i}_{\ell,(j)})_{i\in[N],j\in[2]}$ is independently generated with the same procedure. We therefore find

$$\mathbb{V}\mathrm{ar}\left[P^\ell_{t+\frac{1}{2N}} \,\middle|\, \mathbf{P}_t\right] = \frac{1}{(2N)^2} \sum_{i\in[N],j\in[2]} \mathbb{V}\mathrm{ar}\left[G^{\lfloor 2Nt\rfloor+1,i}_{\ell,(j)} \,\middle|\, \mathbf{P}_t\right] = \frac{1}{2N} \mathbb{V}\mathrm{ar}\left[G^{\lfloor 2Nt\rfloor+1,1}_{\ell,(1)} \,\middle|\, \mathbf{P}_t\right].$$

Because $G_{\ell,(1)}^{\lfloor 2Nt \rfloor+1,1}$ is a Bernoulli variable we find

$$\mathbb{V}\mathbf{ar}\left[P^{\ell}_{t+\frac{1}{2N}} \,|\, \mathbf{P}_t\right] = \frac{1}{2N}\mathbb{E}\left[G_{\ell,(1)}^{\lfloor 2Nt \rfloor+1,1} \,|\, \mathbf{P}_t\right]\mathbb{E}\left[1 - G_{\ell,(1)}^{\lfloor 2Nt \rfloor+1,1} \,|\, \mathbf{P}_t\right].$$

Since conditional on $P^{\ell}_{t+\frac{1}{2N}}$, $G_{\ell,(1)}^{\lfloor 2Nt \rfloor+1,1}$ has law *Bernoulli*$(P^{\ell}_{t+\frac{1}{2N}})$, we find

$$\mathbb{V}\mathbf{ar}\left[P^{\ell}_{t+\frac{1}{2N}} \,|\, \mathbf{P}_t\right] = \frac{1}{2N}\mathbb{E}\left[P^{\ell}_{t+\frac{1}{2N}} \,|\, \mathbf{P}_t\right]\mathbb{E}\left[1 - P^{\ell}_{t+\frac{1}{2N}} \,|\, \mathbf{P}_t\right]. \tag{20}$$

Using the first-order approximation $\mathbb{E}\left[P^{\ell}_{t+\frac{1}{2N}} \,|\, \mathbf{P}_t\right] \simeq P^{\ell}_t$ yields (17).

##### 3.3.3 Second moment: cross coefficients

We can use again the fact that $G_{\ell_1,(j_1)}^{\lfloor 2Nt \rfloor+1,i_1}$ and $G_{\ell_1,(j_2)}^{\lfloor 2Nt \rfloor+1,i_2}$ are independently generated whenever $j_1 \neq j_2$ or $i_1 \neq i_2$ to find

$$\mathbb{C}\mathbf{ov}\left[P^{\ell_1}_{t+\frac{1}{2N}}, P^{\ell_2}_{t+\frac{1}{2N}} \,|\, \mathbf{P}_t\right] = \frac{1}{2N}\mathbb{C}\mathbf{ov}\left[G_{\ell_1,(1)}^{\lfloor 2Nt \rfloor+1,1}, G_{\ell_2,(1)}^{\lfloor 2Nt \rfloor+1,1} \,|\, \mathbf{P}_t\right].$$

If the population at time $t + \frac{1}{2N}$ is in HWLE (H1), then this last term is zero, which yields (18).

### 4 The polygenic equation from the gene's eye-view

We now assume that the system is at statistical equilibrium, writing $\mathbb{P}^*$ for the corresponding probability. We use mean-field approximations to obtain the system Eq. (7-9) for $(P_t, \Delta^*, \varepsilon_t)$ where $\varepsilon_t := \Delta_t - \Delta^*$.

This section is structured as follows

- In Section 4.1, we apply the mean-field hypothesis (H3) to $(\Delta_t)_{t\geq 0}$.
- In Section 4.2, we discuss the decoupling of $(\Delta_t)_{t\geq 0}$ and $\vec{P}_t$ under (H4) and (H4'), which lets us obtain (7) by replacing $\Delta_t$ in (5) with its mean value $\Delta^*$.
- In Section 4.3, we recover (9) for $(\varepsilon_t)_{t\geq 0}$.

#### 4.1 Dynamics of the trait mean

Here we will derive the following equation for $\Delta_t$

$$\begin{aligned}\mathrm{d}\Delta_t = \frac{1}{\tau}\Bigg( &- \Delta_t \times \frac{1}{|\bar\theta|} \times \mathbb{E}^*\left[2(L\alpha)^2 P_t(1-P_t)\right] \;+\; \frac{1}{|\bar\theta|} \times \mathbb{E}^*\left[2L^2\alpha^3\left(P_t - \frac{1}{2}\right)P_t(1-P_t)\right] \\ &+ \frac{L\omega_e^2}{|\bar\theta|} \times \mathbb{E}^*\left[2L\alpha\left(\theta^+(1-P_t) - \theta^- P_t\right)\right]\Bigg)\mathrm{d}t \;+\; \frac{1}{\sqrt{\tau}} \times \sqrt{\frac{\omega_e^2}{|\bar\theta|}\mathbb{E}^*\left[(2L\alpha)^2 P_t(1-P_t)\right]}\mathrm{d}B_t^{\Delta}\end{aligned} \tag{21}$$

The derivation is as follows. From (4), the dynamics of $\Delta_t$ are given by

$$\begin{aligned}\mathrm{d}\Delta_t =& \sum_{\ell=1,\dots,L} \alpha_\ell 2\mathrm{d}P^{\ell}_t \\ =& 2\sum_{\ell\in[L]} \alpha_\ell\left(\xi_{\Delta_t,\alpha_\ell}(P^\ell_t)P^\ell_t(1-P^\ell_t)P^\ell_t(1-P^\ell_t)\mathrm{d}t + (\theta^+_\ell(1-P^\ell_t) - \theta^-_\ell P^\ell_t)\mathrm{d}t + \sqrt{P^\ell_t(1-P^\ell_t)}\mathrm{d}B^\ell_t\right)\end{aligned}$$

We find from (6)

$$\begin{aligned}\mathrm{d}\Delta_t = 2\sum_{\ell\in[L]} -\Delta_t\frac{\alpha_\ell^2}{\omega_e^2}P^\ell_t(1-P^\ell_t)\mathrm{d}t \;+\; 2\sum_{\ell\in[L]}\frac{\alpha_\ell^3}{\omega_e^2}\left(P^\ell_t - \frac{1}{2}\right)P^\ell_t(1-P^\ell_t)\mathrm{d}t \\ + 2\sum_{\ell\in[L]}\alpha_\ell\left(\theta^+_\ell(1-P^\ell_t) - \theta^-_\ell P^\ell_t\right)\mathrm{d}t \;+\; 2\sum_{\ell\in[L]}\alpha_\ell\sqrt{P^\ell_t(1-P^\ell_t)}\mathrm{d}B^\ell_t\end{aligned}$$

We now express this as a function of $\tau$ from (11)

$$
\mathrm{d}\Delta_t = \frac{1}{\tau}\Bigg( \frac{2L\omega_e^2}{|\bar\theta|} \sum_{\ell\in[L]} -\Delta_t \frac{\alpha_\ell^2}{\omega_e^2} P_t^\ell(1-P_t^\ell) \quad + \quad \frac{2L\omega_e^2}{|\bar\theta|} \sum_{\ell\in[L]} \frac{\alpha_\ell^3}{\omega_e^2}\left(P_t^\ell - \frac{1}{2}\right) P_t^\ell(1-P_t^\ell)
$$
$$
+ \frac{2L\omega_e^2}{|\bar\theta|} \sum_{\ell\in[L]} \alpha_\ell\left(\theta_\ell^+(1-P_t^\ell) - \theta_\ell^- P_t^\ell\right) \Bigg)\mathrm{d}t \quad + \quad \frac{1}{\sqrt{\tau}} \sum_{\ell\in[L]} 2\alpha_\ell \sqrt{\frac{L\omega_e^2}{|\bar\theta|} P_t^\ell(1-P_t^\ell)}\mathrm{d}B_t^\ell
$$

which we rewrite

$$
\mathrm{d}\Delta_t = \frac{1}{\tau}\Bigg( -\frac{2}{|\bar\theta|}\Delta_t \times \frac{1}{L}\sum_{\ell\in[L]} (L\alpha_\ell)^2 P_t^\ell(1-P_t^\ell) \quad + \quad \frac{2}{|\bar\theta|} \times \frac{1}{L} \sum_{\ell\in[L]} L^2\alpha_\ell^3 \left(P_t^\ell - \frac{1}{2}\right) P_t^\ell(1-P_t^\ell)
$$
$$
+ \frac{L\omega_e^2}{|\bar\theta|} \times \frac{1}{L} \sum_{\ell\in[L]} 2L\alpha_\ell\left(\theta_\ell^+(1-P_t^\ell) - \theta_\ell^- P_t^\ell\right) \Bigg)\mathrm{d}t \quad + \quad \frac{1}{\sqrt{\tau L}} \sum_{\ell\in[L]} 2L\alpha_\ell \sqrt{\frac{\omega_e^2}{|\bar\theta|} P_t^\ell(1-P_t^\ell)}\mathrm{d}B_t^\ell. \quad (22)
$$

We now use (H3) on the first, second and third term

$$
\frac{1}{L}\sum_{\ell\in[L]} (L\alpha_\ell)^2 P_t^\ell(1-P_t^\ell) \simeq \mathbb{E}^*\left[(L\alpha)^2 P_t(1-P_t)\right] \quad (23)
$$
$$
\frac{1}{L}\sum_{\ell\in[L]} L^2\alpha_\ell^3\left(P_t^\ell - \frac{1}{2}\right)P_t^\ell(1-P_t^\ell) \simeq \mathbb{E}^*\left[L^2\alpha^3\left(P_t - \frac{1}{2}\right)P_t(1-P_t)\right] \quad (24)
$$
$$
\frac{1}{L}\sum_{\ell\in[L]} 2L\alpha_\ell\left(\theta_\ell^+(1-P_t^\ell) - \theta_\ell^- P_t^\ell\right) \simeq \mathbb{E}^*\left[2L\alpha\left(\theta^+(1-P_t) - \theta^- P_t\right)\right] \quad (25)
$$

The last term of (22) is a Brownian term with quadratic variation

$$
\frac{1}{\tau L}\sum_{\ell\in[L]} (2L\alpha_\ell)^2 \times \frac{\omega_e^2}{|\bar\theta|} P_t^\ell(1-P_t^\ell) \simeq \frac{1}{\tau} \times \frac{\omega_e^2}{|\bar\theta|}\mathbb{E}^*\left[(2L\alpha)^2 P_t(1-P_t)\right] \quad (26)
$$

where we used again (H3). We thus obtain (21) with $B^\Delta$ a Brownian motion such that for any $\ell \in [L]$,

$$
\frac{\mathrm{d}}{\mathrm{d}t}\left\langle \Delta, P^\ell \right\rangle_t = 2\alpha_\ell \frac{\mathrm{d}}{\mathrm{d}t}\left\langle P^\ell \right\rangle_t = 2\alpha_\ell \times P_t^\ell(1-P_t^\ell) \sim \frac{1}{L} \quad (27)
$$

where $\langle \cdot, \cdot \rangle$ denotes quadratic variation.

#### 4.2 Decoupling of the trait and the typical locus

We obtain the polygenic system Eq. (7-9) from the SDE for $(P_t^\ell)_{t\geq 0}$ in (5) and that for $(\Delta_t)_{t\geq 0}$ (21), by showing that we may replace $\xi_{\Delta_t,\alpha_\ell}$ with $\xi_{\Delta^*,\alpha_\ell}$ in the SDE for $P_t^\ell$, (5).

Crucially, our derivation assumes either (H4) or (H4').

*Derivation under (H4).* At stationarity, equation (21) is the equation of an Ornstein-Uhlenbeck process with fluctuations of order $\omega_e$. It follows that

$$
\xi_{\Delta_t,\alpha_\ell} = \xi_{\Delta^*,\alpha_\ell} + O(1/(L\omega_e))
$$

In particular, because $\omega_e^{-2} \ll L^2$, we get that $\xi_{\Delta_t,\alpha_\ell} - \xi_{\Delta^*,\alpha_\ell} = o(1)$. □

*Derivation under (H4').* We invoke the principle of **time-averaging** which we illustrate in Fig. 2.

This crucially relies on the fact that $(\Delta_t)_{t\geq 0}$ evolves on a timescale of $\tau \ll 1$. This can be seen as follows

- from (21) we know that $(\Delta_t)_{t\geq 0}$ is an Ornstein-Uhlenbeck process with autocorrelation $\rho$, and in particular it evolves on a characteristic timescale of $\frac{1}{\rho}$
- (H4'b) and (10) implies $\rho \sim \frac{1}{\tau}$
- (H4'c) implies $\tau \ll 1$.

Let us now recall the principle of time-averaging. Suppose we know $(P_0, \Delta_0)$, where $\Delta_0 = \Delta^* + O(\omega_e)$. Let us consider what the first-order increments of $P_t$ are, where $t$ is chosen such that $\tau \ll t \ll 1$. The SDE for $P_t$ can be written from (5) as

$$P_t = P_0 + \int_0^t \left(\xi_{\Delta_u,\alpha}(P_u)P_u(1-P_u) + \left(\theta^+(1-P_u) - \theta^- P_u\right)\right) \mathrm{d}u + \int_0^t \sqrt{P_u(1-P_u)}\, \mathrm{d}B_u^P$$

for some Brownian motion $B^P$. We write from (6)

$$\xi_{\Delta_u,\alpha} = -\alpha \frac{\Delta_u - \Delta^*}{\omega_e^2} + \xi_{\Delta^*,\alpha}.$$

Therefore, to replace $\xi_{\Delta_u,\alpha}$ by $\xi_{\Delta^*,\alpha}$ in the equation for $P_t$, we must show

$$\left| \int_0^t -\alpha \frac{\Delta_u - \Delta^*}{\omega_e^2} P_u(1-P_u)\mathrm{d}u \right| \ll t$$

From (H4'd), on the interval $[0,t]$ the typical locus $P_t$ scarcely evolves, and we may therefore suppose for $u \in [0,t]$

$$P_u \simeq P_0.$$

Therefore, we must show

$$\left| \int_0^t -\alpha \frac{\Delta_u - \Delta^*}{\omega_e^2} \mathrm{d}u \right| \ll t$$

Because the fluctuations of $(\Delta_u)_{u \le t}$ are of order $\omega_e$ and since (H4'a) implies $\frac{\alpha}{\omega_e} \sim 1$, we only have to show that

$$\frac{1}{t}\int_0^t \Delta_u \mathrm{d}u \simeq \Delta^*$$

This ergodic theorem (see for instance [32]) crucially hinges on (H4'b), which tells us that the characteristic timescale for the evolution of $(\Delta_u)_{u\ge 0}$ is $\tau$.

As we have done throughout this appendix, we defer to future work a rigorous proof of this time-averaging, but we do note that a result close to the one needed here has already been obtained by [21]. A rigorous proof in our system will face difficulties which are not tackled in [21], in particular the fact that $B^\Delta$ and $B^P$ are not independent (see (27)).

□

#### 4.3 Recovering the Ornstein-Uhlenbeck process

Here we recover the Ornstein-Uhlenbeck SDE (9) for $(\varepsilon_t)_{t\ge 0}$ from (21). Defining $\varepsilon_t := \Delta_t - \Delta^*$, we get from (21)

$$\begin{aligned}\mathrm{d}\varepsilon_t = -\varepsilon_t \times \frac{1}{\tau|\bar\theta|} \times \mathbb{E}^*\left[2(L\alpha)^2 P_t(1-P_t)\right] \;+\; \frac{1}{\sqrt{\tau}} \times \sqrt{\frac{\omega_e^2}{|\bar\theta|}\mathbb{E}^*\left[(2L\alpha)^2 P_t(1-P_t)\right]}\mathrm{d}B_t^\Delta \\ + \frac{1}{\tau}\left( \frac{1}{|\bar\theta|} \times \mathbb{E}^*\left[2L^2\alpha^3\left(P_t - \frac{1}{2}\right)P_t(1-P_t)\right] + \frac{L\omega_e^2}{|\bar\theta|} \times \mathbb{E}^*\left[2L\alpha\left(\theta^+(1-P_t) - \theta^- P_t\right)\right] \right. \\ \left. - \Delta^* \times \frac{1}{|\bar\theta|} \times \mathbb{E}^*\left[2(L\alpha)^2 P_t(1-P_t)\right] \right)\mathrm{d}t\end{aligned}$$

It remains to show

$$\Delta^* = \frac{\mathbb{E}^*\left[2L^2\alpha^3\left(P_t - \frac{1}{2}\right)P_t(1-P_t)\right] + L\omega_e^2\mathbb{E}^*\left[2L\alpha\left(\theta^+(1-P_t) - \theta^- P_t\right)\right]}{\mathbb{E}^*\left[2(L\alpha)^2 P_t(1-P_t)\right]}. \tag{28}$$

where

$$\Delta^* = 2L\mathbb{E}^*[\alpha P_t] - \eta.$$

We start by noticing that because the system is at stationarity, we have

$$\frac{\mathrm{d}}{\mathrm{d}t}\mathbb{E}^*[P_t \mid (\alpha,\theta)] = 0.$$

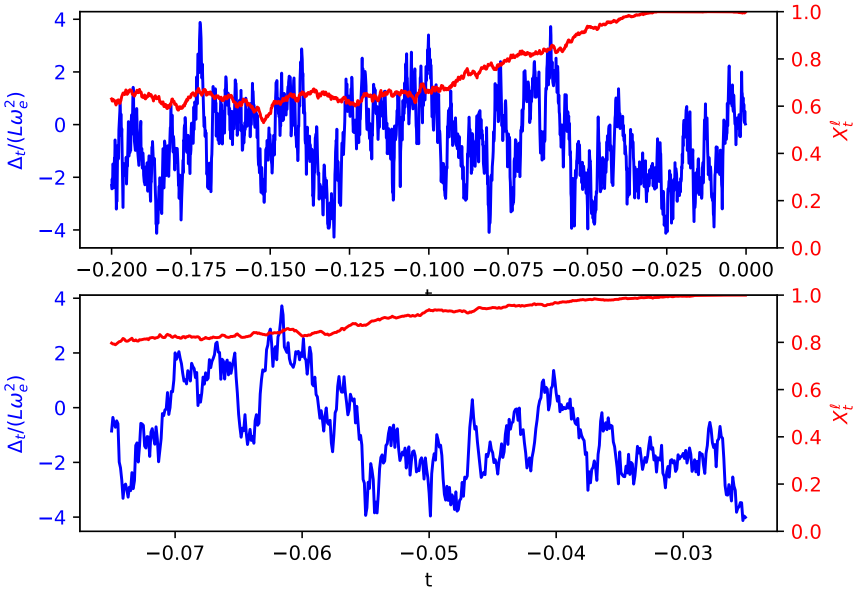


Figure 2: The slow/fast principle applied in strong selection. The population was evolved with $\omega_e^{-2} = 2L^2$ (strong selection) and $L = 1{,}000$, $N = 5{,}000, \eta = 1.2, \theta = (0.1, 0.2)$, and $\alpha_\ell$ has law $Exponential(L)$. In the left figure, we see that $(\Delta_t)_{t\geq 0}$ evolves with very short excursions away from its mean value before returning there. Meanwhile, $P_t^\ell$ only explores the segment $[0.6, 0.9]$, a small portion of the state space of $P_t$. On the right-hand side, we zoom in on a time window in which $P_t^\ell$ stays effectively constant, whereas $\Delta_t$ evolves very quickly. In particular, the fluctuations of $\Delta_t$ do not impact $P_t$. The time-averaging principle is then to consider a small time interval $dt$ with $\tau \ll dt \ll 1$, to replace $\Delta_t$ in the equation of $P_t^\ell$ with its time average over $[t, t+dt]$, and consider that the law of $P_t^\ell$ does not evolve on $[t, t+dt]$.

Applying (7) to $\mathbb{E}^*[P_t \mid (\alpha, \theta)]$, we find

$$\frac{\mathrm{d}}{\mathrm{d}t}\mathbb{E}^*[P_t \mid (\alpha,\theta)] = \mathbb{E}^*\left[\xi_{\Delta^*,\alpha}(P_t)P_t(1-P_t) + \theta^+(1-P_t) - \theta^- P_t \;\middle|\; (\alpha,\theta)\right] = 0.$$

From the definition of $\xi_{\Delta^*,\alpha}$ in (6) we find

$$\mathbb{E}^*\left[\left(-\frac{\alpha}{\omega_e^2}\Delta^* + \frac{\alpha^2}{\omega_e^2}\left(P_t - \frac{1}{2}\right)\right)P_t(1-P_t) + \left(\theta^+(1-P_t) - \theta^- P_t\right) \;\middle|\; (\alpha,\theta)\right] = 0.$$

We rewrite this, multiplying by $2L^2\alpha\omega_e^2$

$$\Delta^*\mathbb{E}^*\left[2(L\alpha)^2 P_t(1-P_t) \mid (\alpha,\theta)\right]$$
$$= \mathbb{E}^*\left[2L^2\alpha^3\left(P_t - \frac{1}{2}\right)P_t(1-P_t) + L\omega_e^2 \times 2L\alpha\left(\theta^+(1-P_t) - \theta^- P_t\right) \;\middle|\; (\alpha,\theta)\right]$$

Taking the expectation with respect to $(\alpha, \theta)$, this yields (28).

### 5 Observables and scalings

Here, we consider that the polygenic limit holds, that is, that the system Eq. (7-8) holds for $(P_t, \Delta^*)$. We do not assume that (9) for $(\varepsilon_t)_{t\geq 0}$ holds unless stated otherwise.

In Section 5.1, we show how the macroscopic observables of the system can be computed from the stationary distribution of $\vec{P}_t$. In section 5.4 we briefly discuss the ultra-weak selection regime ($\omega_e^{-2} \ll L$). In section 5.2, we discuss how macroscopic observables scale under (A1-6). In section 5.3, we introduce the bias-correcting selection coefficient $s^*$ and discuss its behavior in moderate selection ($L \ll \omega_e^{-2} \ll L^2$). In section 5.5, we recover the traditional Stochastic House-of-Cards approximation as a first-order expansion of the genetic variance when $\alpha$ has absolute-normal distribution.

#### 5.1 Observables at stationarity

With the fixed point equation (14), we showed how to compute $\Delta^* = \mathbb{E}^*[\Delta_t]$ and the distribution of the typical locus $\vec{P}_t$. In this subsection we detail how to compute the theoretical predictions for other observables. We assume Eq. (7-8) and HWLE (H1) hold.

We define the following observables at stationarity

$$\sigma_t^2 := \mathbf{Var}_{\mathbf{P}_t}[Z(g)] \qquad \bar{W}_t := \mathbf{E}_{\mathbf{P}_t}[W(Z(g))] \qquad V_t := \mathbf{Var}_{\mathbf{P}_t}[W(Z(g))] \tag{29}$$

$$\nu^2 := \mathbb{V}\mathbf{ar}[\Delta_t] \qquad \rho_u := -\ln\left(\frac{\mathbb{C}\mathbf{ov}\left[\Delta_t, \Delta_{t+u}\right]}{\mathbb{V}\mathbf{ar}[\Delta_t]}\right) \tag{30}$$

where $t, u \geq 0$. They are respectively the trait variance, the fitness load, the fitness variance, the fluctuations of the trait mean and its log-autocorrelation function.

We argue

$$\sigma_t^2 \simeq 2L\mathbb{E}^*[\alpha^2 P_t(1-P_t)] \tag{31}$$

$$\bar{W}_t = -\frac{1}{2\omega^2}\left(\sigma_t^2 + \Delta_t^2\right) \tag{32}$$

$$V_t \simeq \frac{\sigma_t^2(\sigma_t^2 + 2\Delta_t^2)}{2\omega^4} \tag{33}$$

Furthermore if (9) holds then

$$\nu = \omega_e \qquad \forall u \geq 0, \qquad \rho_u = u\rho \tag{34}$$

where $\rho$ was defined in (10).

**Remark 1.** *Because $\mathbb{V}\mathbf{ar}^*[\sigma_t^2] \ll \mathbb{E}^*[\sigma_t^2]^2$, we will consider $\sigma_t^2$ as a constant in the rest of this work rather than a fluctuating quantity, writing $\sigma^2 = \sigma_t^2$.*

**Remark 2.** *The formula for $\bar{W}_t$ was well-known, for instance in [13].*

Let us start by computing a first-order approximation for the genetic variance $\sigma_t^2$. We have

$$\begin{aligned}
\sigma_t^2 =& \mathbf{Var}_{\mathbf{P}_t}\left[\sum_{\ell \in [L]} \alpha_\ell g_\ell\right] \\
\simeq& \sum_{\ell \in [L]} 2\alpha_\ell^2 P_t^\ell (1 - P_t^\ell) \\
=& L \times \frac{1}{L} \sum_{\ell \in [L]} 2\alpha_\ell^2 P_t^\ell (1 - P_t^\ell)
\end{aligned}$$

where we used the HWLE hypothesis (H1) to neglect cross-correlations. Using a mean-field approximation (H3) we get (31).

We turn to $\bar{W}_t$

$$\begin{aligned}
\bar{W}_t =& -\frac{1}{2\omega^2}\mathbf{E}_{\mathbf{P}_t}[(Z(g) - \eta)^2] \\
=& -\frac{1}{2\omega^2}\left(\mathbf{Var}_{\mathbf{P}_t}[Z(g)] + \mathbf{E}_{\mathbf{P}_t}[Z(g) - \eta]^2\right)
\end{aligned}$$

We thus get (32).

Third, we compute $V_t$

$$\begin{aligned}
V_t =& \frac{1}{4\omega^4}\mathbf{Var}_{\mathbf{P}_t}\left[(Z(g) - \eta)^2\right] \\
=& \frac{1}{4\omega^4}\mathbf{Var}_{\mathbf{P}_t}\left[(Z(g) - \bar{z}_t)^2 - 2(Z(g) - \bar{z}_t)\Delta_t\right] \\
=& \frac{1}{4\omega^4}\left(\mathbf{Var}_{\mathbf{P}_t}\left[(Z(g) - \bar{z}_t)^2\right] - 4\mathbf{Cov}_{\mathbf{P}_t}\left[(Z(g) - \bar{z}_t)^2, Z(g) - \bar{z}_t\right]\Delta_t + 4\sigma_t^2\Delta_t^2\right)
\end{aligned}$$

where in the second line we used $\bar{z}_t - \eta = \Delta_t$ and in the third line $\mathbf{Var}_{\mathbf{P}_t}[Z(g)] = \sigma_t^2$. We simplify the computations of $V_t$ by approximating the law of $Z(g)$ under $\mathbf{E}_{\mathbf{P}_t}$ with a $\mathcal{N}(\bar{z}_t, \sigma_t^2)$ distribution (this is a consequence of the Central Limit Theorem under HWLE (H1), provided there is sufficient genetic variability). Under this approximation, the covariance between $(Z(g) - \bar{z}_t)^2$ and $Z(g) - \bar{z}_t$ is zero and

$$\mathbf{Var}_{\mathbf{P}_t}[(Z(g) - \bar{z}_t)^2] \simeq \mathbf{E}_{\mathbf{P}_t}[(Z(g) - \bar{z}_t)^4] - \sigma_t^4 \simeq 2\sigma_t^4$$

We thus obtain

$$V_t \simeq \frac{1}{4\omega^4}\left(2\sigma_t^4 + 4\sigma_t^2\Delta_t^2\right).$$

This yields the result.

The last two equalities are obtained from (9). When $\omega_e^{-2} \gg L$, (9) implies that $\varepsilon_t$ is an Ornstein-Uhlenbeck. Furthermore, standard properties of Ornstein-Uhlenbeck processes, yield that $\Delta_t$ has autocorrelation structure

$$\forall t_1 < t_2, \qquad \mathbb{C}\mathbf{ov}[\varepsilon_{t_1}, \varepsilon_{t_2}] = \omega_e^2 e^{-\rho(t_2 - t_1)}$$

This yields (34).

#### 5.2 Scaling of observables

We now assume (A2-5) hold. We claim

$$|\Delta^*| \sim L\omega_e^2 \qquad\qquad \sigma \sim \sqrt{\frac{|\bar{\theta}|}{L}} \tag{35}$$

If furthermore (9) holds for $(\varepsilon_t)_{t\geq 0}$ then

$$\rho \sim \frac{1}{\tau} \sim \frac{|\bar{\theta}|}{L\omega_e^2} \qquad \mathbb{E}^*[\bar{W}_t] \sim -\frac{1}{2N}\left(\frac{|\bar{\theta}|}{L\omega_e^2} + L^2\omega_e^2\right) \qquad \mathbb{E}^*[V_t] \sim \frac{|\bar{\theta}|}{(2N)^2}\left(\frac{|\bar{\theta}|}{(L\omega_e^2)^2} + L\right). \tag{36}$$

Let us detail the computations.

*Magnitude of* $\Delta^*$. We must show $|\Delta^*| \sim L\omega_e^2$. Specifically, we will show that if $\Delta^*$ satisfies the fixed-point equation (14), then $x^* := \frac{\Delta^*}{L\omega_e^2}$ satisfies a non-degenerate equation and in particular has order 1. Without loss of generality, we assume $2I(0) - \eta > 0$, so that in particular $\Delta^* > 0$.

We rewrite (14) as

$$\Delta^* = 2\mathbb{E}^*\left[(L\alpha)\int p\Pi_{\Delta^*,\alpha,\theta}(p)\mathrm{d}p\right] - \eta.$$

From the definition of $\Pi_{\delta,a,\theta}$ in (13) we have

$$\Pi_{\Delta^*,\alpha,\theta}(p) = \frac{\Pi_{0,\alpha,\theta}(p)}{\int \Pi_{0,\alpha,\theta}(p')e^{-2\frac{\Delta^*}{\omega_e^2}\alpha p'}\mathrm{d}p'}e^{-2\frac{\Delta^*}{\omega_e^2}\alpha p}$$

In particular, the fixed point equation becomes

$$\Delta^* = 2\mathbb{E}^*\left[(L\alpha)\frac{\int p\Pi_{0,\alpha,\theta}(p)e^{-2\frac{\Delta^*}{\omega_e^2}\alpha p}\mathrm{d}p}{\int \Pi_{0,\alpha,\theta}(p)e^{-2\frac{\Delta^*}{\omega_e^2}\alpha p}\mathrm{d}p}\right] - \eta.$$

Rewriting this in terms of $x^*$, we get

$$x^* L\omega_e^2 = 2\mathbb{E}^*\left[(L\alpha)\frac{\int p\Pi_{0,\alpha,\theta}(p)e^{-2(L\alpha)x^*p}\mathrm{d}p}{\int \Pi_{0,\alpha,\theta}(p)e^{-2(L\alpha)x^*p}\mathrm{d}p}\right] - \eta. \tag{37}$$

Let us show that $x^*$ necessarily satisfies $x^* \sim 1$, by showing that we neither have $x^* \ll 1$ nor $x^* \gg 1$.

If $x^*$ satisfied $x^* \ll 1$, from (A5) we have $x^* L\omega_e^2 \ll 1$ and therefore (37) yields

$$|2I(0) - \eta| \ll 1$$

which is impossible from (A6). Let us now show that we cannot have $x^* \gg 1$. It can be checked that the mass of $\Pi_{0,\alpha,\theta}$ between 0 and $\varepsilon < 1/2$ is of order $\frac{\theta^-}{\theta^+ + \theta^-}\varepsilon^{2\theta^+}$. For $\alpha \sim 1/L$ and $\theta$ such that $\theta^+ \sim \theta^- \lesssim 1$, it follows that

$$\frac{\Pi_{0,\alpha,\theta}(p)}{\int \Pi_{0,\alpha,\theta}(p')e^{-2(L\alpha)xp'}\mathrm{d}p'}e^{-2(L\alpha)xp}$$

is close to a Dirac mass on 0 whenever $x \gg 1$. In particular,

$$\frac{\int p\Pi_{0,\alpha,\theta}(p)e^{-2(L\alpha)xp}\mathrm{d}p}{\int \Pi_{0,\alpha,\theta}(p)e^{-2(L\alpha)xp}\mathrm{d}p} \ll 1$$

If the solution to the fixed-point equation (37) satisfied $x^* \gg 1$, then we would have

$$x^* L\omega_e^2 \simeq -\eta$$

The left-hand side is non-negative whereas the right-hand side is negative of order 1 from (A4), which yields a contradiction. □

*Magnitude of $\sigma$.* Recall from (6) and (13)

$$\begin{aligned}\xi_{\delta,a}(p) :=& -\frac{a\delta}{\omega_e^2} + \frac{a^2}{\omega_e^2}\left(p - \frac{1}{2}\right)\\ \Pi_{\delta,a,\theta}(p) :=& C_{\delta,a,\theta}\, p^{2\theta^+ -1}(1-p)^{2\theta^- -1} e^{2\int_0^p \xi_{\delta,a}(u)\,\mathrm{d}u}.\end{aligned}$$

with $C_{\delta,a,\theta}$ a normalization constant. Then if $\alpha \sim \frac{1}{L}$ we find from $\Delta^* \sim L\omega_e^2$ and (5) that $\xi_{\Delta^*,\alpha} \sim 1$ and therefore

$$\int p(1-p)\Pi_{\Delta^*\alpha,\theta}(p)\mathrm{d}p \sim \frac{\int p^{2\theta^+}(1-p)^{2\theta^-}\mathrm{d}p}{\int p^{2\theta^+-1}(1-p)^{2\theta^--1}\mathrm{d}p} = \frac{Beta(2\theta^+ + 1, 2\theta^- + 1)}{Beta(2\theta^+, 2\theta^-)}$$

where *Beta* is the Beta function. In particular

$$\int p(1-p)\Pi_{\Delta^*\alpha,\theta}(p)\mathrm{d}p \sim \frac{\theta^+\theta^-}{|\theta|}$$

where $|\theta| = \theta^+ + \theta^-$. As a consequence, if we consider a typical locus $\vec{P}_t$ we have

$$\mathbb{E}^*[(L\alpha)^2 P_t(1-P_t)] \sim \mathbb{E}\left[(L\alpha)^2\frac{\theta^+\theta^-}{|\theta|}\right] \sim |\bar{\theta}|$$

where we used (A3). We thus find

$$\sigma^2 = L\mathbb{E}^*[\alpha^2 P_t(1-P_t)] \sim \frac{|\bar{\theta}|}{L}.$$

□

*Derivation of Eq. (36).* We obtain from (10) and $\sigma \sim \sqrt{|\bar{\theta}|/L}$ that $\rho \sim \frac{1}{\tau} = \frac{|\bar{\theta}|}{L\omega_e^2}$ from the definition of $\tau$ in (11). Finally, (32) implies

$$\mathbb{E}^*[\bar{W}_t] \sim -\frac{1}{\omega^2}\left(\sigma^2 + (\Delta^*)^2 + \omega_e^2\right)$$

using $\mathbb{E}^*[\Delta_t^2] = (\Delta^*)^2 + \omega_e^2$ from (34). Using $\omega^2 = 2N\omega_e^2$ and the previous estimates for $\sigma^2, \Delta^*$ we find

$$\begin{aligned}\mathbb{E}^*[\bar{W}_t] \sim& -\frac{1}{2N\omega_e^2}\left(\frac{|\bar{\theta}|}{L} + (L\omega_e^2)^2 + \omega_e^2\right)\\ \sim& -\frac{1}{2N}\left(\frac{|\bar{\theta}|}{L\omega_e^2} + L^2\omega_e^2 + 1\right).\end{aligned}$$

The result follows from the fact that under (A5), we have $L^2\omega_e^2 \gtrsim 1$, and therefore the last term can always be absorbed into the second term. Similarly (33) yields

$$\begin{aligned}\mathbb{E}^*[V_t] \sim& \frac{\sigma^2}{\omega^4}\left(\sigma^2 + (\Delta^*)^2 + \omega_e^2\right)\\ \sim& \frac{|\bar{\theta}|}{(2N)^2\omega_e^4 L}\left(\frac{|\bar{\theta}|}{L} + (L\omega_e^2)^2 + \omega_e^2\right)\\ \sim& \frac{|\bar{\theta}|}{(2N)^2}\left(\frac{|\bar{\theta}|}{(L\omega_e^2)^2} + L + \frac{1}{L\omega_e^2}\right)\end{aligned}$$

and the last term can be absorbed in $L$. □

#### 5.3 The bias-correcting coefficient

Define the bias-correcting coefficient $s^*$ as

$$s^* := -\frac{\Delta^*}{L\omega_e^2} \tag{38}$$

From (35), we know that $s^*$ is always of order 1. In this subsection we claim that for $L \ll \omega_e^{-2} \ll L^2$ (moderate selection), $s^*$ is solution to

$$2L\mathbb{E}\left[\alpha\frac{\theta^+}{|\theta|} \times \frac{{}_1F_1(2\theta^+ + 1\ ;\ 2|\theta| + 1\ ;\ 2s^*L\alpha)}{{}_1F_1(2\theta^+\ ;\ 2|\theta|\ ;\ 2s^*L\alpha)}\right] \simeq \eta,$$

where ${}_1F_1$ is the confluent hypergeometric function. In particular, $s^*$ is independent of $\omega_e^{-2}$.

The derivation is as follows.

In moderate selection, we find from (35) that $|\Delta^*| \sim L\omega_e^2 \ll 1 \sim \eta$. It follows that (14) can be rewritten

$$2I(\Delta^*) \simeq \eta. \tag{39}$$

On the other hand, we find from (6) that for a typical $\alpha$,

$$\xi_{\Delta^*,\alpha}(p) \simeq -s^*L\alpha \sim 1$$

where we neglected $\frac{\alpha^2}{\omega_e^2} \sim \frac{1}{L^2\omega_e^2} \ll 1$. It follows

$$\Pi_{\Delta^*,\alpha,\theta}(p) \simeq \frac{p^{2\theta^+-1}(1-p)^{2\theta^--1}e^{2s^*L\alpha p}}{\int y^{2\theta^+-1}(1-y)^{2\theta^--1}e^{2s^*L\alpha y}\mathrm{d}y}$$

From the definition of $I$ in (15) we get

$$I(\Delta^*) \simeq \mathbb{E}\left[L\alpha\frac{\int p^{2\theta^+}(1-p)^{2\theta^--1}e^{2s^*L\alpha p}\mathrm{d}p}{\int p^{2\theta^+-1}(1-p)^{2\theta^--1}e^{2s^*L\alpha p}\mathrm{d}p}\right] = \mathbb{E}\left[L\alpha\frac{\theta^+}{|\theta|} \times \frac{{}_1F_1(2\theta^+ + 1\ ;\ 2|\theta| + 1\ ;\ 2s^*L\alpha)}{{}_1F_1(2\theta^+\ ;\ 2|\theta|\ ;\ 2s^*L\alpha)}\right]$$

where we recall the notation $|\theta| := \theta^+ + \theta^-$. Carrying this into (39), we get the first result.

#### 5.4 Ultra-weak selection regime

Let us briefly discuss the case of ultra-weak-selection regime ($\omega_e^{-2} \ll L$).

Because of the definition of $\Delta_t$ and (1), we necessarily have $|\Delta_t| \leq 2$ as long as $\eta \in [0, 2]$. It follows from the definition of $\xi$ (6) that when $\omega_e^{-2} \ll L$, $|\xi_{\Delta_t,\alpha}| \ll 1$, and therefore $P_t^\ell$ evolves as a neutral Wright-Fisher diffusion. In this regime, an individual locus is not affected by selection in a detectable way. In particular, the macroscopic observables $\Delta^*, \sigma$ can be computed from the neutral distribution $\Pi_{0,0,\theta}$.

We also note that both [31] in his equation (B4a) (which extended the analysis of [30]) and [12] (for instance following equation (S3.2) in his Appendix) suggest that $\sigma^2$ can reasonably be replaced by its neutral expectation. As can easily be seen from the stationary distribution of $P_t$ in (13) and the fact that $s^*$ is of order 1 when $\omega_e^{-2} \gtrsim L$, this only holds strictly under ultra-weak selection ($\omega_e^{-2} \ll L$).

#### 5.5 The Stochastic House-of-Cards approximation

The Stochastic House-of-Cards approximation (SHoC) was simultaneously derived independently in [9, 22, 24]. These derivations rely on the infinite-allele model used to derive the basic House-of-Cards model in [42]. We can rederive this result in our setting, under the following assumptions

- $|\bar{\theta}| \ll 1$
- $\eta = z_M$ (with $z_M$ the mutational equilibrium from Section 2.2), so that $s^* = 0$ (no mutational bias)
- $L \ll \omega_e^{-2} \ll L^2$ (this corresponds to the moderate selection regime, but we specifically look for a first-order Taylor expansion of $\sigma^2$ in $(L\omega_e)^{-2}$)
- $\alpha$ conditioned on $\theta$ is distributed as the absolute value of an $\mathcal{N}(0, v)$ variable, with fixed variance $v$. This assumption violates our simplifying assumption (A 1) of uniform boundedness of the $(\alpha_\ell)_{\ell\in[L]}$, but we still expect the polygenic equation (7) to hold even when this distribution is very degenerate (see Fig. S6).

We will show the following first-order expansion

$$\sigma^2 \simeq \frac{2Lv\mathbb{E}\left[\frac{2\theta^+\theta^-}{\theta^++\theta^-}\right]}{1+\frac{v}{2\omega_e^2}}. \tag{40}$$

We will see that this result crucially hinges on the assumption that $\alpha$ is normally distributed, and is therefore of questionable biological relevance.

*Derivation of* (40). Applying Itô's formula on $2L\alpha^2 P_t(1-P_t)$, and taking expectation we find a contribution of selection equal to

$$-\mathbb{E}^*\left[L\frac{\alpha^4}{\omega_e^2}P_t(1-P_t)(1-2P_t)^2\right] = -\mathbb{E}^*\left[L\frac{\alpha^4}{\omega_e^2}\int_0^1 p(1-p)(1-2p)^2\Pi_{0,\alpha,\theta}(p)\mathrm{d}p\right],$$

using $\Pi$ from (13).

We then write

$$\int_0^1 p(1-p)\Pi_{0,\alpha,\theta}(p)\mathrm{d}p = \frac{\int_0^1 p^{2\theta^+}(1-p)^{2\theta^-}e^{-\frac{\alpha^2}{\omega_e^2}p(1-p)}\mathrm{d}p}{\int_0^1 p^{2\theta^+-1}(1-p)^{2\theta^--1}e^{-\frac{\alpha^2}{\omega_e^2}p(1-p)}\mathrm{d}p}.$$

Our assumption $\omega_e^{-2} \ll L^2$ lets us expand the exponential, and we find up to a first order in $(L\omega_e)^{-2}$

$$\int_0^1 p(1-p)\Pi_{0,\alpha,\theta}(p)\mathrm{d}p \simeq \frac{Beta(2\theta^++1,2\theta^-+1)-\frac{\alpha^2}{\omega^2}Beta(2\theta^++2,2\theta^-+2)}{Beta(2\theta^+,2\theta^-)-\frac{\alpha^2}{\omega^2}Beta(2\theta^++1,2\theta^-+1)},$$

with *Beta* the Beta function. This can be rearranged using $\theta^+ + \theta^- \ll 1$ and properties of *Beta* to find

$$\int_0^1 p(1-p)\Pi_{0,\alpha,\theta}(p)\mathrm{d}p \simeq \frac{2\theta^+\theta^-}{\theta^++\theta^-}\times\left(1-\frac{\alpha^2}{6\omega^2}\right).$$

Carrying this into (31) we find

$$\sigma^2 \simeq 2L\mathbb{E}\left[\alpha^2\frac{2\theta^+\theta^-}{\theta^++\theta^-}\right] - 2L\mathbb{E}\left[\frac{\alpha^4}{6\omega_e^2}\frac{2\theta^+\theta^-}{\theta^++\theta^-}\right].$$

Because we assumed $\alpha$ conditioned on $\theta$ has normal distribution with variance $v$, we find $\mathbb{E}\left[\alpha^4|\theta\right] = 3v^2$ and in particular

$$\sigma^2 \simeq 2Lv\mathbb{E}\left[\frac{2\theta^+\theta^-}{\theta^++\theta^-}\right]\left(1-\frac{v}{2\omega_e^2}\right).$$

We get the result from $1-u \simeq 1/(1+u)$ with $u = v/(2\omega_e^2)$. □

##### 5.6 Lynch's formula

Here, we recover Lynch's formula for $\Delta^*$ (formula (13b) in [30]). This formula only holds under the following assumptions

- selection is weak or moderate ($\omega_e^{-2} \ll L^2$)
- the mutation rate $|\theta|$ is constant across loci (but the ratio $|\theta^+/\theta^-|$ may vary across loci).

We will show the following formula for the expected trait value at equilibrium

$$\mathbb{E}^*[\bar{z}_t] = \frac{\omega_e^2}{\omega_e^2+\frac{\sigma^2}{|\theta|}}z_M + \frac{\sigma^2}{|\theta|}\times\frac{1}{\omega_e^2+\frac{\sigma^2}{|\theta|}}\eta. \tag{41}$$

This formula is not very convenient because it gives $\mathbb{E}^*[\bar{z}_t]$ as a function of $\sigma^2$, which is not itself known. In [31], this difficulty is bypassed by replacing $\sigma^2$ with its neutral expectation, which is only valid when mutational bias is small (unlike what we assume here).

Our derivation will show among other things that this formula cannot be expected to hold when $|\theta|$ is not constant. We take this opportunity to note that [30] goes on to simplify this expression assuming $\omega_e^2 \ll \sigma^2/|\theta|$, which in view of (35) is equivalent to $\omega_e^{-2} \gg L$ (moderate selection). His simplified equation is

$$\mathbb{E}^*[\bar{z}_t] \simeq \frac{\omega_e^2|\theta|}{\sigma^2}z_M + \eta.$$

But this correction is obtained by neglecting terms of order $\omega_e^2/\sigma^2$, so the first term of this new equation should also be neglected, in which case it becomes simply $\mathbb{E}^*[\bar{z}_t] \simeq \eta$, which matches our analysis of moderate selection in Section 5.3.

*Derivation of 41.* We start by rewriting (7) as

$$\mathrm{d}P_t = -\frac{\alpha\Delta^*}{\omega_e^2}P_t(1-P_t)\,\mathrm{d}t + \left(\theta^+(1-P_t) - \theta^- P_t\right)\mathrm{d}t + \sqrt{P_t(1-P_t)}\,\mathrm{d}B_t^P,$$

where we neglected the second term in the selection coefficient (6) due to $\omega_e^{-2} \ll L^2$. We rewrite this

$$\mathrm{d}P_t = -\frac{\Delta^*}{\omega_e^2}\alpha P_t(1-P_t)\,\mathrm{d}t + \left(\theta^+ - |\theta|\frac{\eta}{2L} - |\theta|\left(P_t - \frac{\eta}{2L}\right)\right)\mathrm{d}t + \sqrt{P_t(1-P_t)}\,\mathrm{d}B_t^P.$$

Multiplying by $2L\alpha/|\theta|$ and taking expectation, we find

$$0 = -\frac{\Delta^*}{\omega_e^2|\theta|}2L\mathbb{E}^*[\alpha^2 P_t(1-P_t)] + \mathbb{E}^*\left[2L\alpha\frac{\theta^+}{|\theta|} - \alpha\eta\right] - 2L\mathbb{E}^*[\alpha P_t] + \eta,$$

which we rewrite using (31), (8) and $\mathbb{E}[\alpha] = 1$ as

$$0 = -\frac{\Delta^*}{|\theta|\omega_e^2}\sigma^2 + 2L\mathbb{E}^*\left[\alpha\frac{\theta^+}{|\theta|}\right] - \eta - \Delta^*.$$

Crucially, the previous rewriting would not have been possible had we not assumed $|\theta|$ to be constant. Finally, rearranging we get

$$\Delta^* = \frac{z_M - \eta}{1 + \frac{\sigma^2}{|\theta|\omega_e^2}}$$

where we recall $z_M$ the mutational equilibrium from Section 2.2. We thus get the result from $\Delta^* = \mathbb{E}^*[\bar{z}_t] - \eta$. □

### 6 Breakdown of the polygenic limit

Here, we discuss (H1-4') in light of Section 5. We assume (A1-6) hold and the polygenic equation for $(P_t, \Delta^*)_{t\geq 0}$ in Eq. (7-9) hold. In particular, the orders of magnitude derived in Section 5 let us characterize the parameter values for which (A1-6) are consistent with (H1-4'). We argue that if (N1-2) is not satisfied, then (35) is incompatible with (H1-4'), and in particular that the polygenic system in Eq. (7-8) breaks down. In Section 6.5, we argue that if (N3) is not satisfied, then (9) cannot provide an accurate description of the fluctuations of $(\varepsilon_t)_{t\geq 0}$.

#### 6.1 Discussion of (H2)

We illustrate our method by discussing (H2), which assumes that the fitness $F(Z(g))$ is very concentrated around its mean value under $\mathbf{E}_{\mathbf{P}_t}$. In particular, this requires the fitness variance $V_t$ to be very small. In light of (36), this requires

$$\left(\frac{|\bar{\theta}|}{2NL\omega_e^2}\right)^2 \ll 1 \qquad\qquad \frac{|\bar{\theta}|L}{(2N)^2} \ll 1$$

which we rewrite

$$2N \gg \frac{|\bar{\theta}|}{L\omega_e^2} \qquad\qquad 2N \gg \sqrt{|\bar{\theta}|L}.$$

Both these conditions are satisfied under (N1).

Let us be more precise. In Section 3, (H2) was used to write

$$\mathbf{Cov}_{\mathbf{P}_t}[F(Z(g)), g_\ell] \simeq \mathbf{Cov}_{\mathbf{P}_t}[\hat{F}_t(W(Z(g)) - \bar{W}_t), g_\ell] \qquad\qquad \mathbf{E}_{\mathbf{P}_t}[F(Z(g))] \simeq \hat{F}_t$$

By Taylor expansion, this is justified if

$$\sum_{k>1}\frac{1}{k!}\left|\mathbf{Cov}_{\mathbf{P}_t}[(W(Z(g)) - \bar{W}_t)^k, g_\ell]\right| \ll \left|\mathbf{Cov}_{\mathbf{P}_t}[(W(Z(g)) - \bar{W}_t), g_\ell]\right|$$

$$\sum_{k\geq 2}\frac{1}{k!}\mathbf{E}_{\mathbf{P}_t}[|W(Z(g)) - \bar{W}_t|^k] \ll 1$$

Such estimates can be obtained if we assume that $Z(g)$ is normally distributed under $\mathbf{E}_{\mathbf{P}_t}$ conditioned on $g_\ell$. The trait's eye-view, which will be presented in Section 7, yields an alternative derivation of (5) which relies precisely on this assumption.

#### 6.2 Discussion of HWLE (H1)

HWLE (H1) is undoubtedly the most delicate assumption above. We will discuss it by referring to the rich literature on the subject.

Hypothesis (H1) was discussed in [14] (preprint). In this article, a Wright-Fisher diffusion is considered in which LE is not assumed. In this setting, the recombination rate $\rho$ is defined as the number of recombination events for a given lineage over $2N$ generations. In our setting, in which recombination occurs every generation, this corresponds to $\rho = 2N$. The effect of recombination is to force the population close to the Wright manifold, on which the population is at LE. The main result of [14] adapted to our setting is that if

$$2N \gg \omega_e^{-4} \ln(L)^2 \tag{42}$$

then we can effectively assume LE when computing the dynamics of $P_t^\ell$. That is, we may neglect the effect of linked selection on $P_t^\ell$. (42) is admittedly biologically unrealistic, and it is likely not optimal, but it is derived entirely *a priori*. In our simulations (Fig. 2 of main text), $2N = 1,000$, $L = 100$ and $\omega_e^{-4} \in [L^2, L^4]$, and the criterion (42) reads $1,000 \gg [2 \times 10^5, 2 \times 10^9]$. In particular, (42) is assuredly not optimal. We can try and get a better idea of the conditions for the breakdown of (H1) using the Quasi-Linkage Equilibrium approach from statistical genetics [34]. This approach assumes *a priori* that the first-order effect of LD on gene dynamics and macroscopic observables can be described exclusively with two-loci correlations (neglecting cumulants of order 3 or greater).

##### 6.2.1 Bulmer effect

When selection is sufficiently strong, we have $\Delta^* \ll \sigma$. Bulmer suggested in [8] that for an additive trait under Gaussian selection, LD can be neglected as long as $\ln(L)\frac{\sigma_t^2}{\omega^2} \ll 1$ where the $\ln(L)$ factor comes from the choice of single-point uniform crossover as a recombination mechanism (see Eq. (10) of [8], see also for greater clarity Eq. (26) of [33] and the paragraph under). This criterion can be derived from the Quasi-Linkage Equilibrium approach [34], considering that LD only appears through selection picking advantageous combinations of genes, which when the population is close to the optimum results in negative LD (for Gaussian selection). The criterion can be rewritten

$$2N \gg \sigma_t^2 \ln(L)\omega_e^{-2}. \tag{43}$$

In light of (35), this can be rewritten

$$2N \gg |\bar{\theta}|\frac{\ln(L)}{L\omega_e^2}$$

which is satisfied under (N1). If this equation is not satisfied, then we expect to enter the Quasi-Linkage Equilibrium regime [26, 34], in which LD appears between loci. This would mainly manifest itself in the decrease of the genetic variance in the trait $\sigma^2$ relative to the theoretical prediction due to negative LD [33].

In main text Fig. 2C, we see this effect appear around $\omega_e^{-2} \sim 200$ and $\omega_e^{-2} \sim 2000$, respectively for $2N = 100$ and $2N = 1000$. Translating this in terms of (43), we get that the critical value of $\omega_e^{-2}$ at which LD ceases to be negligible satisfies

$$|\bar{\theta}|\frac{\ln(L)}{2NL\omega_e^2} \sim 0.03.$$

We discuss in more details the Bulmer effect in SI 10, and in particular obtain a first-order correction and test it against simulations (Fig. S8).

##### 6.2.2 The Hill-Robertson effect

When selection is sufficiently weak that $\Delta^* \gg \sigma$, the population remains far from the optimum and can be described using a linear selection model, in which the logfitness of an organism is approximated by linear selection

$$\tilde{W}(z) := -\frac{\Delta^*}{\omega^2}(z - \bar{z})$$

where $\bar{z} = 2L\mathbb{E}^*[\alpha P_t] = \Delta^* + \eta$.

In such a setting, the appearance of negative LD for small population sizes is known as Hill-Robertson effect [20]. The rationale is as follows: LD leads the genetic variance $\sigma_t^2$ to oscillates randomly and quickly with respect to the genic variance $\sigma_{G,t}^2$ defined with

$$\sigma_{G,t}^2 := \sum_{\ell \in [L]} 2\alpha_\ell^2 P_t^\ell (2 - P_t^\ell).$$

We illustrate this oscillation in Fig. 3 in an extreme scenario in which $L = 1,000$ and $N = 20$, for weak Gaussian selection and directional selection of equivalent magnitude. When LD is positive, then there is a lot of genetic

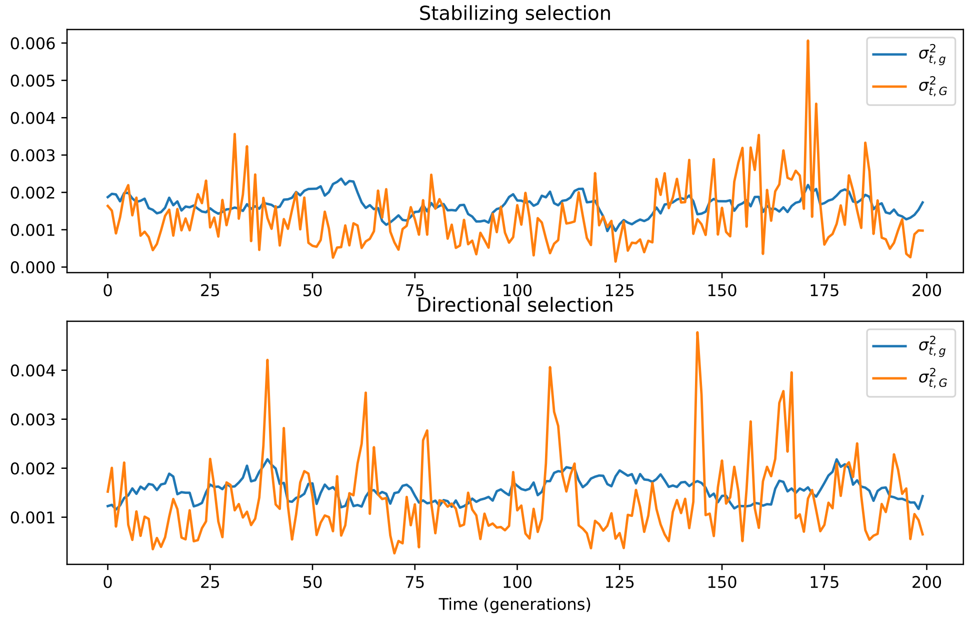


Figure 3: The Hill-Robertson effect. The population was evolved with $N = 10, L = 200, \theta = (0.1, 0.2)$, $\eta = 1.2$, for $T = 1000N$ generations, either under weak Gaussian selection ($\eta = 1.2, \omega_e^{-2} = 2/L$) or on the corresponding directional selection regime. The $(\alpha_\ell)_{\ell\in[L]}$ have law $Exponential(L)$. The **genetic trait variance** $\sigma_{t,G}$ is the variance of $Z(g)$ for $g$ sampled within the population at random, whereas the **genic trait variance** corresponds to the variance the population would have if it was in HWLE, labelled $\sigma^2_{t,g}$ in the figure. When blue is on top of orange, the population displays negative LD, whereas when orange is on top of blue, the population displays positive LD. The Hill-Robertson effect corresponds to the fact that positive LD only exists in very short bursts (for example around generation 105 on the top figure), whereas negative LD exists on longer excursions (for instance after generation 130 on the top figure).

variance, in which case selection will act efficiently to decrease the genetic variance (and in particular LD). This explains why positive LD only manifests itself in very short excursions. On the other hand, when LD is negative, then the genetic variance $\sigma_t^2$ is reduced with respect to the genic variance $\sigma^2_{G,t}$, and selection is inefficient. In this setting, only recombination will, on a longer timescale, bring LD back to zero.

We illustrate the Hill-Robertson effect and the corresponding breakdown of the polygenic limit of Eq. (7-8) in Fig. 4. We see that this breakdown is quite limited, since even with as few as $N = 10$ organisms with $L = 100$ loci each, LD only decreases the genetic variance by about 5%.

Using the theory of Quasi-Linkage Equilibrium, we can try and obtain a quantitative criterion for the appearance of the Hill-Robertson effect. In Eq. (44) of Section VI.B of [34], it is claimed that the Hill-Robertson effect at a locus $\ell \in [L]$ can be neglected as soon as

$$\sum_{\ell'\neq\ell} P_t^{\ell'}(1-P_t^{\ell'})\left(\frac{1}{r_{\ell,\ell'}}\xi_{\Delta^*,\alpha_{\ell'}}(P_t^{\ell'})\right)^2 \ll 1$$

where $r_{\ell_1\ell_2}$ is the rate of recombination between $\ell_1$ and $\ell_2$, in our system $r_{\ell_1\ell_2} = \frac{|\ell_1-\ell_2|}{L}2N$. This criterion was originally obtained in [2] assuming $|\bar{\theta}| \ll 1$, but should be taken with caution as it is known that the usual Quasi-Linkage Equilibrium approach does not describe well the equilibrium distribution of allelic frequencies (see e.g [35, 17, 15]). Future work on this aspect should instead consider approaches such as suggested in [5], which has not yet been adapted to recombining genomes.

Taking the expectation, and using from (38) that $\xi_{\Delta^*,\alpha_{\ell'}}$ is typically of order 1, we obtain the criterion

$$\sum_{\ell'\in[L]\setminus\{\ell\}} \mathbb{E}^*[P_t(1-P_t)] \times \left(\frac{L}{|\ell-\ell'|2N}\right)^2 \ll 1$$

which yields from (35)

$$|\bar{\theta}|\frac{L^2}{(2N)^2} \ll 1$$

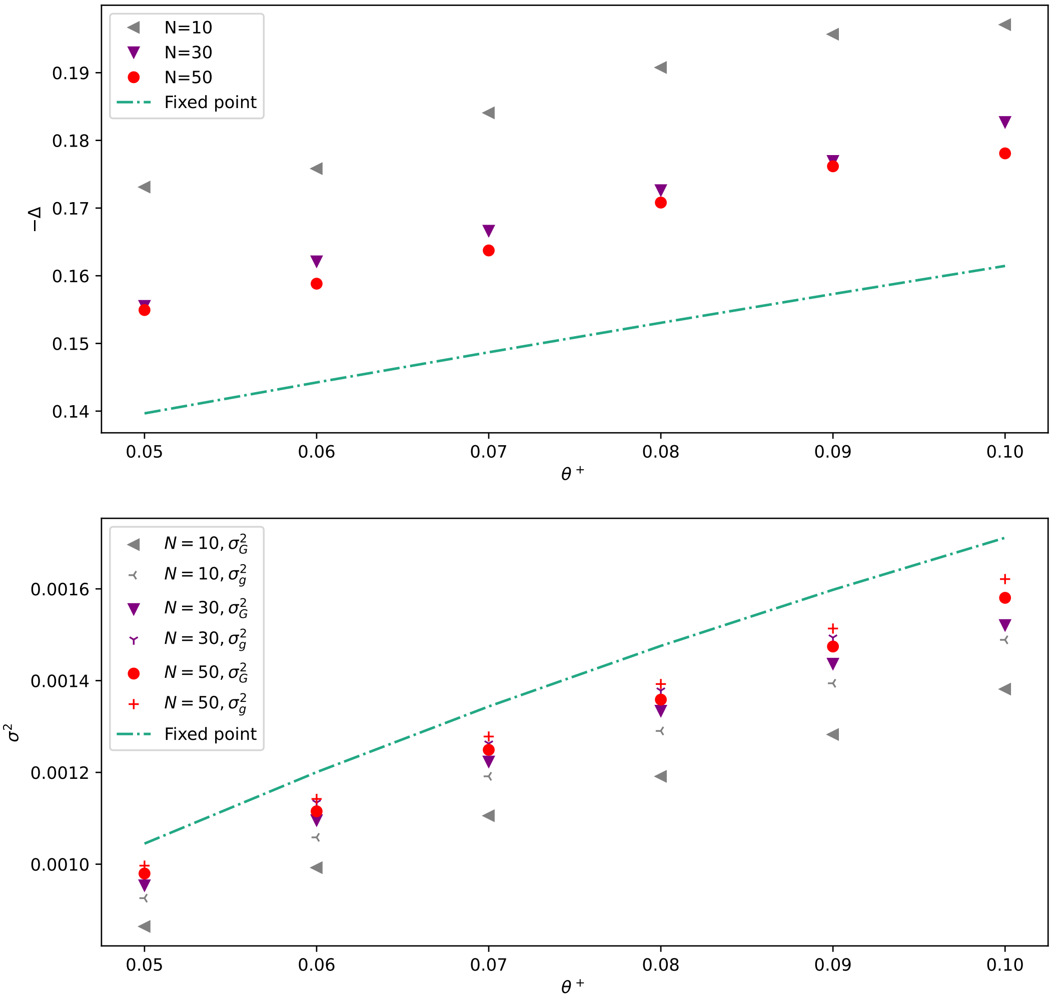


Figure 4: The Hill-Robertson effect. The population was evolved at various population sizes with $L = 200, \theta^- = 2\theta^+$, $\eta = 1.2$, for $T = 4,000N$ generations, half of which were used as burn-in. The population was under weak Gaussian selection ($\omega_e^{-2} = 2/L$ and $\eta = 1.2$). Below, the plot distinguishes between genetic variance (the variance in the trait) and the genic variance, corresponding to the label "LE". The genic variance is the variance of the trait if the allelic frequencies were kept unchanged but linkage equilibrium was enforced. The difference between the genetic and the genic variance is a measure of LD.

which we rewrite

$$2N \gg L\sqrt{|\bar{\theta}|}.$$

This is satisfied under (N1).

#### 6.3 Discussion of the mean-field hypothesis (H3)

We will now show that our mean field approximation breaks down when mutation rates are small. As is usual with mean-field approximations [11], let us consider that the error of the mean-field hypothesis (H3) has variance of order $\frac{1}{L}$, that is

$$\mathbb{V}\mathbf{ar}\left[\frac{1}{L}\sum_{\ell\in[L]} f(\vec{P}_t^\ell)\right] \sim \frac{\mathbb{V}\mathbf{ar}[f(\vec{P}_t)]}{L} \tag{44}$$

We consider the mean-field approximation (H3) to be valid as long as this error is much smaller than the mean of $f(\vec{P}_t)$

$$\sqrt{\frac{\mathbb{V}\mathbf{ar}[f(\vec{P}_t)]}{L}} \ll \mathbb{E}^*[f(\vec{P}_t)]$$

Let us now check the validity of the mean-field approximations of Eq. (23-26). From (35), we find

$$\mathbb{E}^*\left[(L\alpha)^2 P_t(1-P_t)\right] \sim|\bar{\theta}| \quad \mathbb{E}^*\left[L^2\alpha^3\left(\frac{1}{2}-P_t\right)P_t(1-P_t)\right] \lesssim\frac{|\bar{\theta}|}{L} \quad \mathbb{E}^*\left[2L\alpha(\theta^+(1-P_t)-\theta^-P_t)\right] \sim|\bar{\theta}|$$

In Section 6.3.1 we argue

$$\mathbb{V}\mathbf{ar}^*\left[(L\alpha)^2 P_t(1-P_t)\right] \sim|\bar{\theta}| \quad \mathbb{V}\mathbf{ar}^*\left[L^2\alpha^3\left(\frac{1}{2}-P_t\right)P_t(1-P_t)\right] \lesssim\frac{|\bar{\theta}|}{L^2} \quad \mathbb{V}\mathbf{ar}^*\left[2L\alpha(\theta^+(1-P_t)-\theta^-P_t)\right] \sim|\bar{\theta}|^2 \tag{45}$$

It follows that the mean-field approximations Eq. (23-26) are valid iff

$$\sqrt{\frac{|\bar{\theta}|}{L}} \ll|\bar{\theta}| \qquad \sqrt{\frac{|\bar{\theta}|}{L^2}} \ll\frac{|\bar{\theta}|}{L} \qquad \sqrt{\frac{|\bar{\theta}|^2}{L}} \ll|\bar{\theta}|$$

This is equivalent to (N2). In Fig. 5, we illustrate the breakdown of the polygenic limit if (N2) is not satisfied.

##### 6.3.1 Computing the errors on the mean-field approximationss

We start with the first equation of (45). Recall from (13) that conditional on $(\alpha,\theta)$, $P_t$ has distribution $\Pi_{\Delta^*,\alpha,\theta}$ which is equivalent to a $Beta(2\theta^+,2\theta^-)$ distribution when $\Delta^* \sim L\omega_e^2$ and $\alpha \sim 1/L$. If $\hat{P}_t$ has law $Beta(2\theta^+,2\theta^-)$ for some $\theta$, standard properties of the Beta distribution yield

$$\begin{aligned}
\mathbb{E}^*[\hat{P}_t^2(1-\hat{P}_t)^2] =& \frac{2\theta^+(2\theta^++1)\theta^-(2\theta^-+1)}{(\theta^++\theta^-)(2\theta^++2\theta^-+1)(2\theta^++2\theta^-+2)} \sim |\theta| \\
\mathbb{E}^*[\hat{P}_t(1-\hat{P}_t)]^2 =& \left(\frac{2\theta^+\theta^-}{\theta^++\theta^-}\right)^2 \sim |\theta|^2 \\
\mathbb{V}\mathbf{ar}^*[\hat{P}_t(1-\hat{P}_t)] =& \mathbb{E}^*[\hat{P}_t^2(1-\hat{P}_t)^2] - \mathbb{E}^*[\hat{P}_t(1-\hat{P}_t)]^2 \sim |\theta|
\end{aligned}$$

if $\theta^+ \sim \theta^- \lesssim 1$. Similarly, conditional on $(\alpha,\theta)$, we have

$$\mathbb{V}\mathbf{ar}^*[P_t(1-P_t) \mid (\alpha,\theta)] \sim |\theta|$$

Using the decomposition

$$\mathbb{V}\mathbf{ar}^*\left[(L\alpha)^2P_t(1-P_t)\right] = \mathbb{E}^*\left[\mathbb{V}\mathbf{ar}^*\left[(L\alpha)^2P_t(1-P_t)\,\middle|(\alpha,\theta)\right]\right] + \mathbb{V}\mathbf{ar}^*\left[\mathbb{E}^*\left[(L\alpha)^2P_t(1-P_t)\,\middle|(\alpha,\theta)\right]\right]$$

we get under (A1-3)

$$\mathbb{V}\mathbf{ar}^*\left[(L\alpha)^2P_t(1-P_t)\right] \sim |\bar{\theta}|.$$

as requested.

The second equation of (45) is obtained similarly. Let us turn to the third one. As above, if $\hat{P}_t$ has law $Beta(2\theta^+,2\theta^-)$ for some $\theta$ with $\theta^+ \sim \theta^- \lesssim 1$ then

$$\mathbb{V}\mathbf{ar}^*[\hat{P}_t] = \frac{\theta^+\theta^-}{|\bar{\theta}|^2(2|\bar{\theta}|+1)} \sim 1.$$

Consider $\tilde{P}_t$ with law $\Pi_{\Delta^*,\alpha,\theta}$ with $\alpha \sim 1/L$. Let us show that we also have $\mathbb{V}\mathbf{ar}[\tilde{P}_t] \sim 1$. Because $\Pi_{\Delta^*,\alpha,\theta}$ is equivalent to a $Beta(2\theta^+, 2\theta^-)$ distribution, we have

$$\begin{aligned}\mathbb{V}\mathbf{ar}^*[\tilde{P}_t] =& \mathbb{E}^*\left[\left(\tilde{P}_t - \mathbb{E}^*[\tilde{P}_t]\right)^2\right]\\ \sim& \mathbb{E}^*\left[\left(\hat{P}_t - \mathbb{E}^*[\tilde{P}_t]\right)^2\right]\\ =& \mathbb{V}\mathbf{ar}^*[\hat{P}_t] + \left(\mathbb{E}^*[\hat{P}_t] - \mathbb{E}^*[\tilde{P}_t]\right)^2\\ \sim& 1\end{aligned}$$

From this, we may conclude that conditional on $(\alpha, \theta)$ we have

$$\mathbb{V}\mathbf{ar}^*\left[P_t \mid (\alpha,\theta)\right] \sim 1$$

Writing the decomposition

$$\begin{aligned}&\mathbb{V}\mathbf{ar}^*\left[2\alpha(\theta^+(1-P_t) - \theta^- P_t)\right]\\ &\qquad = \mathbb{E}^*\left[\mathbb{V}\mathbf{ar}^*\left[2\alpha(\theta^+(1-P_t) - \theta^- P_t)\ \middle|(\alpha,\theta)\right]\right] + \mathbb{V}\mathbf{ar}^*\left[2\mathbb{E}^*\left[\alpha(\theta^+(1-P_t) - \theta^- P_t)\ \middle|(\alpha,\theta)\right]\right]\end{aligned}$$

we get under (A1-3)

$$\mathbb{V}\mathbf{ar}^*\left[2L\alpha(\theta^+(1-P_t) - \theta^- P_t)\right] \sim |\bar{\theta}|^2$$

as requested.

##### 6.3.2 Breakdown of the mean-field hypothesis under small mutation rates

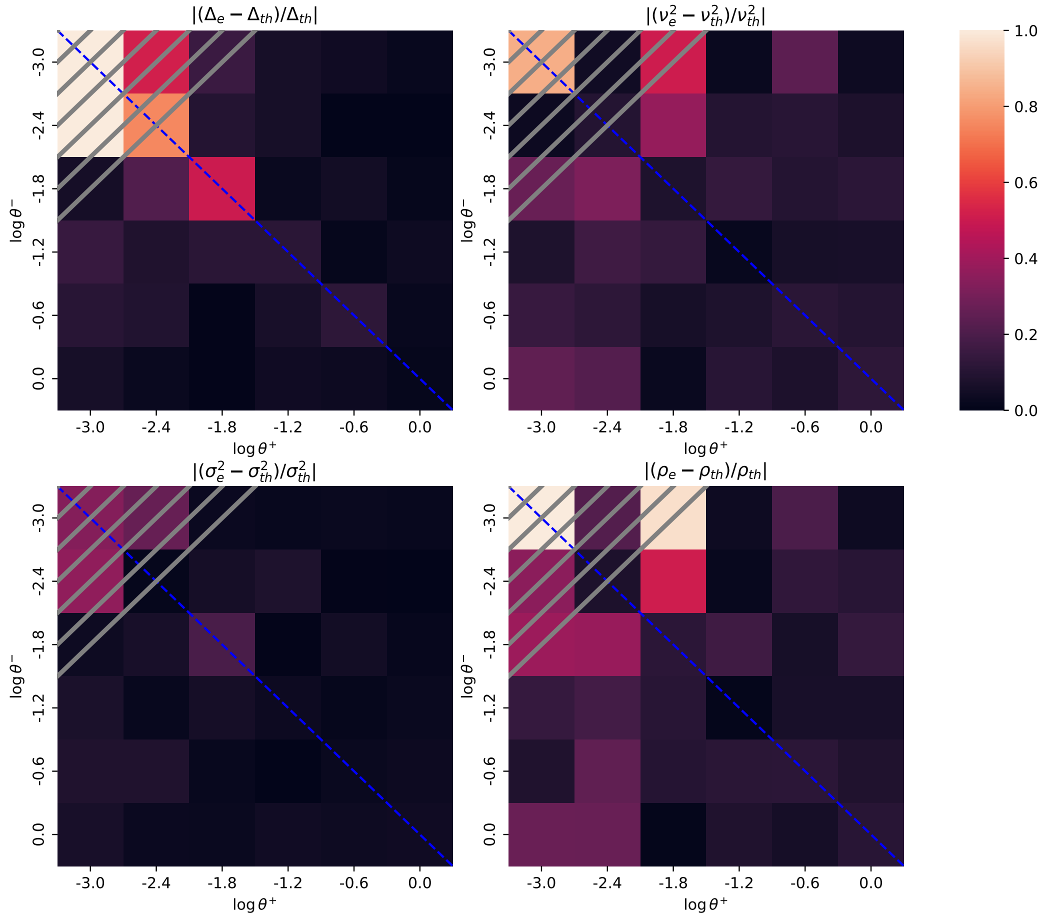


Figure 5: Breakdown of our approximation when hypotheses (N2) and (A3) fail. The population was evolved with $\omega_e^{-2} = 10^3$ (strong selection), $N = 500, L = 100, \eta = 1.2$ under different mutation rates for $T = 500N$ generations, including a burn-in of $250N$ generations. We took the same $(\alpha_\ell)_{\ell\in[L]}$ as in Fig. 2 of the main text. We plot the empirical value of $\Delta$ from simulations ($\Delta_e$) vs our theoretical prediction ($\Delta_{th}$), and similarly for $\nu, \sigma, \rho$. As in Fig. 2 of the main text, the predictions were obtained conditional on the values of $(\alpha_\ell)_{\ell\in[L]}$. In this parameterization, assumption (N2) reads $\theta^+ + \theta^- \gg 10^{-2}$ (non-hashed zone) and (A3) reads $\theta^+ \sim \theta^-$ (blue dashed line).

##### 6.3.3 Breakdown of the mean-field hypothesis when the allele effects have heavy tails.

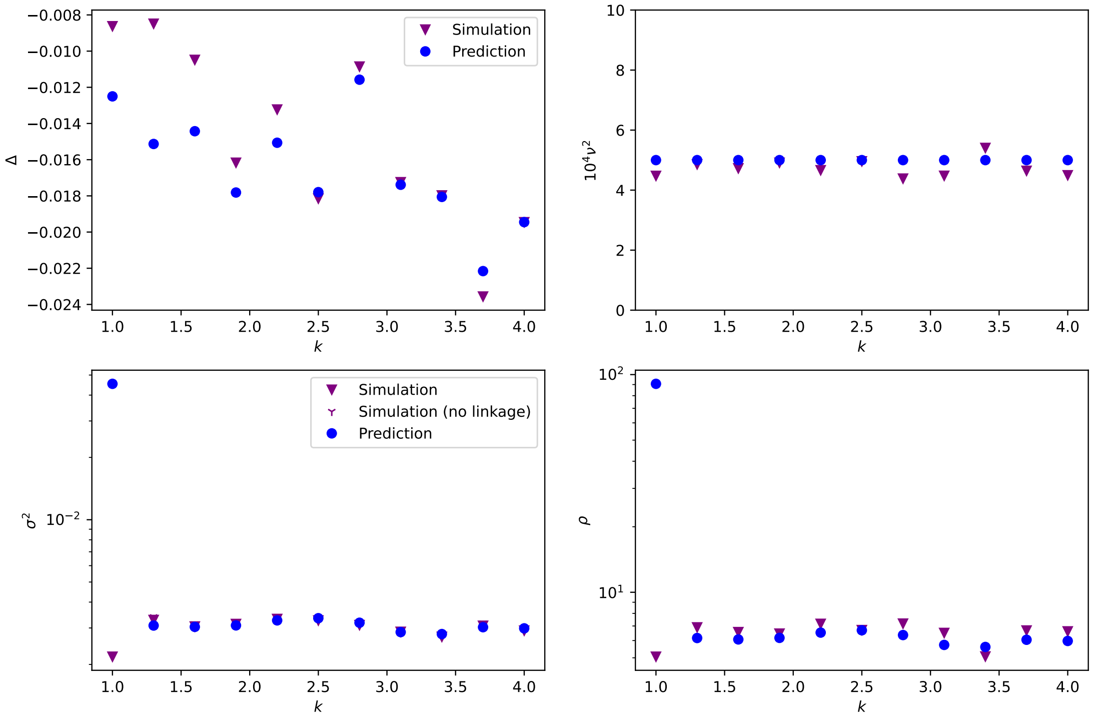


Figure 6: Breakdown of the approximation when $(\alpha_\ell)_{\ell\in[L]}$ has heavy tails. We simulated the system at stationarity for $N = 500, L = 100, \eta = 1.2, \omega_e^{-2} = 10^3, \theta = (0.1, 0.2)$. We sampled $\hat{\alpha}_\ell$ using the $Pareto(k)$ distribution for $k$ between 1 and 4, and set the allelic effect as $\alpha_\ell := \hat{\alpha}_\ell/(\sum_\ell \hat{\alpha}_\ell) = 1$. In particular, the smaller $k$, the heavier the tail of $\alpha$. The prediction was obtained using the same method as in Fig. 5, using the empirical distribution of $(\alpha_\ell)_{\ell\in[L]}$ as the law of $\alpha$. When $k = 1$, a single locus can have a very large effect, $\alpha_\ell \sim 1$, and the mean-field approximation cannot be expected to hold. The prediction seems to hold quite well for $k > 1.5$. This is particularly surprising because the theoretical prediction for $\sigma^2$ and $\rho$ are obtained using the expectation $\mathbb{E}^*[\alpha^2 P_t(1-P_t)]$ and equation (8) for $\Delta^*$ involves $\mathbb{E}^*[\alpha^3 P_t(1-P_t)(1-2P_t)]$, whereas $\mathbb{E}\left[\alpha^2\right]$ and $\mathbb{E}\left[\alpha^3\right]$ are ill-behaved when $k \leq 2$. We defer to future work a theoretical characterization of this breakdown.

#### 6.4 Discussion of (H4-4')

In Section 4.2, (H4-4') were used to argue that $\xi_{\Delta_t,\alpha_\ell}$ can be replaced with $\xi_{\Delta^*,\alpha_\ell}$ in (5). Let us check that under (A5) and (N2), (H4) or (H4') are satisfied.

Under (A5), we have either $\omega_e^{-2} \ll L^2$ or $\omega_e^{-2} \sim L^2$ (strong selection). The former situation corresponds to (H4), while the latter corresponds to (H4'a). Furthermore, (H4'b) is satisfied from (35), and (H4'c) is satisfied under (H4'a) and (N2).

#### 6.5 Breakdown of the equation for the dynamics of the trait mean

It can be seen from Fig. 2B,D in the main text that the theoretical predictions for $(\nu, \rho)$ from (36) fail for weak selection ($\omega_e^{-2} \sim L$), whereas those for $\sigma^2, \Delta^*$ (Fig. 2A,C) still hold. This stems from the fact that the domain of validity of (9) is smaller than that of Eq. (7-9). Here we discuss the breakdown of (9) for the fluctuations of the trait mean $(\varepsilon_t)_{t\geq 0}$ when (N3) is not satisfied, and suggest a proxy equation when mutation rates are constant across loci.

##### 6.5.1 Breakdown of (9) for weak selection

Here, we show why (N3) is necessary for the Ornstein-Uhlenbeck SDE (9) to hold.

Let us start by rewriting (21) as follows

$$\mathrm{d}\Delta_t = \frac{1}{\tau}\Bigg( -\Delta_t \times \frac{1}{|\bar{\theta}|} \times \mathbb{E}^*\left[2(L\alpha)^2 P_t(1-P_t)\right] \; + \; \frac{1}{|\bar{\theta}|} \times \mathbb{E}^*\left[2L^2\alpha^3\left(P_t - \frac{1}{2}\right)P_t(1-P_t)\right] + \frac{L\omega_e^2}{|\bar{\theta}|} \times \mathbb{E}^*\left[2L\alpha\left(\theta^+(1-P_t) - \theta^- P_t\right)\right]\Bigg)\mathrm{d}t \; + \; \frac{1}{\sqrt{\tau}} \times \sqrt{\frac{\omega_e^2}{|\bar{\theta}|}\mathbb{E}^*\left[(2L\alpha)^2 P_t(1-P_t)\right]}\mathrm{d}B_t^{\Delta} + \mathrm{d}E_t$$

where we added the term $E_t$, which is the **error term of the mean-field approximations** Eq. (23-26). Specifically, we define $E_t := E_t^1 + E_t^2 + E_t^3 + E_t^4$ where

$$\mathrm{d}E_t^1 := -\frac{1}{\tau} \times \Delta_t \times \frac{1}{|\bar{\theta}|} \times \left(\frac{1}{L}\sum_{\ell\in[L]} 2(L\alpha_\ell)^2 P_t^\ell(1-P_t^\ell) \; - \mathbb{E}^*\left[2(L\alpha)^2 P_t(1-P_t)\right]\right)\mathrm{d}t$$

$$\mathrm{d}E_t^2 := \frac{1}{\tau|\bar{\theta}|} \times \left(\frac{1}{L}\sum_{\ell\in[L]} 2L^2\alpha_\ell^3\left(P_t^\ell - \frac{1}{2}\right)P_t^\ell(1-P_t^\ell) \; - \mathbb{E}^*\left[2L^2\alpha^3\left(P_t - \frac{1}{2}\right)P_t(1-P_t)\right]\right)\mathrm{d}t$$

$$\mathrm{d}E_t^3 := \frac{L\omega_e^2}{\tau|\bar{\theta}|} \times \left(\frac{1}{L}\sum_{\ell\in[L]} 2L\alpha_\ell\left(\theta_\ell^+(1-P_t^\ell) - \theta_\ell^- P_t^\ell\right) \; - \; \mathbb{E}^*\left[2L\alpha\left(\theta^+(1-P_t) - \theta^- P_t\right)\right]\right)\mathrm{d}t$$

$$\mathrm{d}E_t^4 := \sqrt{\frac{\omega_e^2}{\tau|\bar{\theta}|}}\left(\sqrt{\frac{1}{L}\sum_{\ell\in[L]}(2L\alpha_\ell)^2 P_t^\ell(1-P_t^\ell)} - \sqrt{\mathbb{E}^*\left[(2L\alpha)^2 P_t(1-P_t)\right]}\right)\mathrm{d}B_t^{\Delta}$$

Using $\varepsilon_t := \Delta_t - \Delta^*$ and Section 4.3, (21) can be rewritten

$$\mathrm{d}\varepsilon_t = -\rho\varepsilon_t + \omega_e\sqrt{2\rho}\mathrm{d}B_t^{\Delta} + \mathrm{d}E_t$$

We consider (9) for the description of $(\varepsilon_t)_{t\geq 0}$ to be valid as long as the error term $E_t$ is negligible with respect to the other terms. Since (9) is an Ornstein-Uhlenbeck process with variance $\omega_e^2$ and autocorrelation parameter $\rho \sim \frac{1}{\tau}$ (see (36)), this the contribution of the terms of (9) is of order $\omega$ over a timescale of order $\tau$. Therefore, (9) is valid provided

$$\omega_e \gg \left|\tau\frac{\mathrm{d}}{\mathrm{d}t}E_t^1\right| + \left|\tau\frac{\mathrm{d}}{\mathrm{d}t}E_t^2\right| + \left|\tau\frac{\mathrm{d}}{\mathrm{d}t}E_t^3\right| + \sqrt{\tau\frac{\mathrm{d}}{\mathrm{d}t}\langle E^4\rangle_t}$$

which, in light of (44) we rewrite

$$\omega_e \gg \Delta_t \times \frac{1}{|\bar{\theta}|} \times \sqrt{\frac{\mathbb{V}\mathbf{ar}^*\left[2(L\alpha)^2 P_t(1-P_t)\right]}{L}} + \frac{1}{|\bar{\theta}|}\sqrt{\frac{\mathbb{V}\mathbf{ar}^*\left[2L^2\alpha^3\left(P_t - \frac{1}{2}\right)P_t(1-P_t)\right]}{L}} + \frac{L\omega_e^2}{|\bar{\theta}|} \times \sqrt{\frac{\mathbb{V}\mathbf{ar}^*\left[2L\alpha\left(\theta^+(1-P_t) - \theta^- P_t\right)\right]}{L}} + \sqrt{\frac{\omega_e^2}{|\bar{\theta}|}} \times \sqrt{\frac{\mathbb{V}\mathbf{ar}^*\left[(2L\alpha)^2 P_t(1-P_t)\right]}{L}}$$

In light of (45), this becomes

$$\omega_e \gg \frac{\Delta_t}{|\bar{\theta}|} \times \sqrt{\frac{|\bar{\theta}|}{L}} + \frac{1}{|\bar{\theta}|} \times \sqrt{\frac{|\bar{\theta}|}{L^3}} + \frac{L\omega_e^2}{|\bar{\theta}|} \times \sqrt{\frac{|\bar{\theta}|^2}{L}} + \sqrt{\frac{\omega_e^2}{|\bar{\theta}|}} \times \sqrt{\frac{|\bar{\theta}|}{L}}$$

which yields

$$\omega_e \gg \frac{\Delta_t}{\sqrt{|\bar{\theta}|L}} + \frac{1}{\sqrt{|\bar{\theta}|L^3}} + \omega_e^2\sqrt{L} + \frac{\omega_e}{\sqrt{L}}.$$

We can ignore the second term using (A5)+(N2) and the fourth term on the right-hand side using $L \gg 1$, and use that $\Delta_t \sim L\omega_e^2$ from (35) to get

$$\omega_e \gg \omega_e^2\sqrt{\frac{L}{|\bar{\theta}|}} + \omega_e^2\sqrt{L}.$$

Because $|\bar{\theta}| \lesssim 1$ (A2), the second term is smaller than the first one and we get

$$\omega_e \gg \omega_e^2\sqrt{\frac{L}{|\bar{\theta}|}}$$

which can be rewritten

$$1 \gg \frac{L\omega_e^2}{|\bar{\theta}|}.$$

This is precisely (N3).

##### 6.5.2 A proxy equation for the fluctuations under weak selection

Assumption (N3) is not satisfied in many circumstances, for instance under weak selection ($\omega_e^{-2} \sim L$). From the previous section, this means the Ornstein-Uhlenbeck equation (9) cannot give a good description of the fluctuations of $(\Delta_t)_{t\geq 0}$ for weak selection. Here, we suggest a proxy equation for this regime. We will derive the proxy equation under the assumption

$$|\theta_\ell| = \theta_\ell^+ + \theta_\ell^- = |\bar{\theta}| \text{ is constant across loci.}$$

We will obtain the following proxy SDE for $(\varepsilon_t)_{t\geq 0}$

$$\mathrm{d}\varepsilon_t = -\tilde{\rho}\varepsilon_t\,\mathrm{d}t + \tilde{\nu}\sqrt{2\tilde{\rho}}\,\mathrm{d}B_t^\Delta \tag{46}$$

with

$$\tilde{\rho} := \frac{1}{\tau}\left(\frac{L\sigma^2}{|\bar{\theta}|} + L\omega_e^2\right) \tag{47}$$

$$\frac{1}{\tilde{\nu}^2} := \frac{1}{\omega_e^2} + \frac{|\bar{\theta}|}{\sigma^2}, \tag{48}$$

where we recall $\tau$ from (11). By proxy equation, we mean that (46) is not the correct mathematical object to describe the limit (it unduly neglects the fluctuations of the genetic variance $(\sigma_t^2)_{t\geq 0}$ corresponding to $(E_t^1)_{t\geq 0}$), but can nevertheless yield a sufficient approximation for practical purposes (see Fig. 7). In particular, this approximation is used to compute the theoretical predictions under weak selection for $\nu^2$ and $\rho$ in the main text Fig. 2B and D. Notice how under moderate/strong selection ($L \ll \omega_e^{-2} \lesssim L^2$), we have from (35) $\omega_e^2 \ll \frac{\sigma^2}{|\bar{\theta}|}$ and therefore $\tilde{\rho} \simeq \rho, \tilde{\nu} \simeq \omega_e^2$, thus recovering (36). On the other hand, if $\omega_e^{-2} \ll L$ (ultra-weak selection), then we recover an Ornstein-Uhlenbeck process dominated by mutations.

The corrected formula for $\tilde{nu}^2$ was obtained in [30] (his equation (13C)), using a trait-based approach.

*Derivation of Eq. (46).* Our proxy equation will account for the term $E_t^3$ from Section 6.5.1, but not for the term $E_t^1$. The missing term $E_t^1$ is precisely why this equation is not exact, and corresponds to the fluctuations of the trait variance $(\sigma_t^2)_{t\geq 0}$. In practice, this term slightly inflates the variance $\nu^2 = \mathbb{V}\mathbf{ar}[\Delta_t]$ in Fig. 2 of the main text, and leads the log-autocorrelation function $(\rho_u)_{u\geq 0}$ to depart from linearity (Figure 7).

Using that $|\theta_\ell|$ is constant across loci, we write

$$\begin{aligned}\sum_{\ell\in[L]} 2L\alpha_\ell(\theta_\ell^+(1-P_t^\ell) - \theta_\ell^- P_t^\ell) &= \sum_{\ell\in[L]} 2L\alpha_\ell\theta_\ell^+ \; - \; |\bar{\theta}|L\sum_{\ell\in[L]} 2\alpha_\ell P_t^\ell \\ &= \sum_{\ell\in[L]} 2L\alpha_\ell\theta_\ell^+ \; - \; |\bar{\theta}|L(\Delta_t + \eta)\end{aligned}$$

from the definition of $\Delta_t$ in (4). Using a mean-field approximation we get

$$\sum_{\ell\in[L]} 2L\alpha_\ell(\theta_\ell^+(1-P_t^\ell)-\theta_\ell^- P_t^\ell) \simeq L\mathbb{E}^*[2L\alpha\theta^+] - |\bar{\theta}|L(\Delta_t+\eta).$$

It can be checked that the error of this mean-field approximation is always negligible under (A1). It follows

$$\begin{aligned}\mathrm{d}E_t^3 =& \frac{L\omega_e^2}{\tau|\bar{\theta}|}\left(\mathbb{E}^*[2L\alpha\theta^+] - |\bar{\theta}|\Delta_t + |\bar{\theta}|\eta - \mathbb{E}^*\left[2L\alpha\left(\theta^+(1-P_t)-\theta^-P_t\right)\right]\right)\mathrm{d}t\\ =& \frac{L\omega_e^2}{\tau|\bar{\theta}|}\left(-|\bar{\theta}|\Delta_t + |\bar{\theta}|\eta - \mathbb{E}^*\left[2L\alpha|\theta|P_t\right]\right)\mathrm{d}t\end{aligned}$$

Therefore, accounting for $E_t^3$, (21) becomes

$$\begin{aligned}\mathrm{d}\Delta_t = \frac{1}{\tau}\Bigg(& -\Delta_t \times \frac{1}{|\bar{\theta}|} \times \mathbb{E}^*\left[2(L\alpha)^2P_t(1-P_t)\right] \;+\; \frac{1}{|\bar{\theta}|}\times\mathbb{E}^*\left[2L^2\alpha^3\left(P_t-\frac{1}{2}\right)P_t(1-P_t)\right]\\ &+\frac{L\omega_e^2}{|\bar{\theta}|}\times\mathbb{E}^*\left[2L\alpha\left(\theta^+(1-P_t)-\theta^-P_t\right)\right]\Bigg)\mathrm{d}t \;+\; \frac{1}{\sqrt{\tau}}\times\sqrt{\frac{\omega_e^2}{|\bar{\theta}|}\mathbb{E}^*\left[(2L\alpha)^2P_t(1-P_t)\right]}\mathrm{d}B_t^\Delta\\ &+\frac{L\omega_e^2}{\tau|\bar{\theta}|}\left(-|\bar{\theta}|\Delta_t+|\bar{\theta}|\eta-\mathbb{E}^*\left[2L\alpha|\theta|P_t\right]\right)\mathrm{d}t\end{aligned}$$

This can be rewritten

$$\begin{aligned}\mathrm{d}\Delta_t = \frac{1}{\tau}\Bigg(& -\left(\frac{\mathbb{E}^*\left[2(L\alpha)^2P_t(1-P_t)\right]}{|\bar{\theta}|}+L\omega_e^2\right)\Delta_t \;+\; \frac{1}{|\bar{\theta}|}\times\mathbb{E}^*\left[2L^2\alpha^3\left(P_t-\frac{1}{2}\right)P_t(1-P_t)\right]\\ &+L\omega_e^2\left(-\eta+\frac{\mathbb{E}^*\left[2L\alpha\theta^+\right]}{|\bar{\theta}|}\right)\Bigg)\mathrm{d}t \;+\; \frac{1}{\sqrt{\tau}}\times\sqrt{\frac{\omega_e^2}{|\bar{\theta}|}\mathbb{E}^*\left[(2L\alpha)^2P_t(1-P_t)\right]}\mathrm{d}B_t^\Delta \qquad (49)\end{aligned}$$

This yields

$$\mathrm{d}\Delta_t = -\tilde{\rho}(\tilde{\Delta}^*-\Delta_t)\mathrm{d}t + \tilde{\nu}\sqrt{2\tilde{\rho}}\mathrm{d}B_t^\Delta$$

where

$$\begin{aligned}\tilde{\Delta}^* :=& \frac{1}{\tau\tilde{\rho}}\times\left(\frac{1}{|\bar{\theta}|}\times\mathbb{E}^*\left[2L^2\alpha^3\left(P_t-\frac{1}{2}\right)P_t(1-P_t)\right]+L\omega_e^2\left(-\eta+\frac{\mathbb{E}^*\left[2L\alpha\theta^+\right]}{|\bar{\theta}|}\right)\right)\\ \tilde{\rho} :=& \frac{1}{\tau}\left(\frac{\mathbb{E}^*\left[2(L\alpha)^2P_t(1-P_t)\right]}{|\bar{\theta}|}+L\omega_e^2\right)\\ \tilde{\nu}^2 =& \frac{\frac{\omega_e^2}{|\bar{\theta}|}\mathbb{E}^*\left[(2L\alpha)^2P_t(1-P_t)\right]}{2\left(\frac{\mathbb{E}^*\left[2(L\alpha)^2P_t(1-P_t)\right]}{|\bar{\theta}|}+L\omega_e^2\right)}\end{aligned}$$

To obtain Eq. (46-48), it remains to show that $\tilde{\Delta}^*=\Delta^*$.

From (28) we know

$$\Delta^* = \frac{1}{\tau|\bar{\theta}|\rho}\left(\mathbb{E}^*\left[2L^2\alpha^3\left(P_t-\frac{1}{2}\right)P_t(1-P_t)\right]+L\omega_e^2\mathbb{E}^*\left[2L\alpha\left(\theta^+(1-P_t)-\theta^-P_t\right)\right]\right).$$

It follows

$$\tilde{\Delta}^* = \frac{1}{\tau\tilde{\rho}}\left(\tau\rho\Delta^* - \frac{L\omega_e^2}{|\bar{\theta}|}\mathbb{E}^*\left[2L\alpha\left(\theta^+(1-P_t)-\theta^-P_t\right)\right]+L\omega_e^2\left(-\eta+\frac{\mathbb{E}^*\left[2L\alpha\theta^+\right]}{|\bar{\theta}|}\right)\right)$$

which we rewrite using the definition of $\tau = \frac{L\omega_e^2}{|\bar{\theta}|}$

$$\tilde{\Delta}^* = \frac{1}{\tilde{\rho}}\left(\rho\Delta^* - \mathbb{E}^*\left[2L\alpha\left(\theta^+(1-P_t)-\theta^-P_t\right)\right] - |\bar{\theta}|\eta + \mathbb{E}^*\left[2L\alpha\theta^+\right]\right).$$

This can be rewritten

$$\tilde{\Delta}^* = \frac{1}{\tilde{\rho}}\left(\rho\Delta^* + |\bar{\theta}|\mathbb{E}^*\left[2L\alpha P_t\right] - |\bar{\theta}|\eta\right).$$

Using $\Delta^* = \mathbb{E}^*[2L\alpha P_t] - \eta$, we get

$$\tilde{\Delta}^* = \frac{1}{\tilde{\rho}} \left( \rho \Delta^* + |\bar{\theta}| \Delta^* \right).$$

Because $\tilde{\rho} = \rho + |\bar{\theta}|$, we obtain as claimed

$$\tilde{\Delta}^* = \Delta^*.$$

□

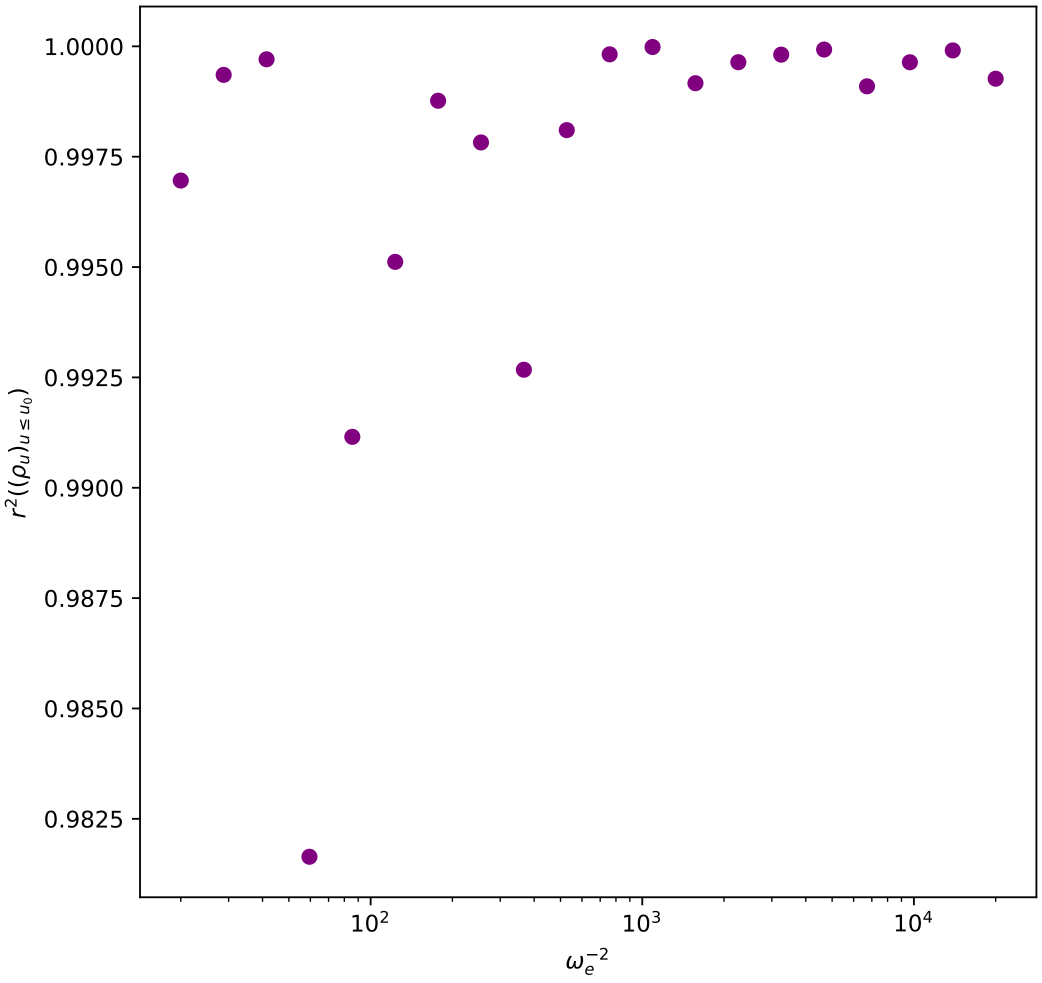


Figure 7: Here we plot the square of Pearson's correlation coefficient $r^2$ for the log-autocorrelation function $(\rho_u)_{u\leq u_0}$ defined in (30) as a function of the selection strength $\omega_e^{-2}$. For each value of $\omega_e^{-2}$, $\rho_u$ is computed for times $u \leq u_0$ with $u_0 = \ln(2)/\rho_{1/(2N)}$, and $r^2$ was computed on $(\rho_u)_{u\leq u_0}$. The time $u_0$ was chosen such that we may expect $e^{-\rho_{u_0}} \simeq 1/2$ for an Ornstein-Uhlenbeck process.
For an Ornstein-Uhlenbeck process, we expect $r^2 = 1$. As expected, the deviation from the expectation of an Ornstein-Uhlenbeck process appears for weak selection.
The simulations were carried out with $L = 100, N = 500$, the same mutation probabilities for all loci $\theta = (0.1, 0.2)$, and allelic effects $(\alpha_\ell)_{\ell\in[L]}$ exponentially distributed with parameter $L$ (in particular $\bar{\alpha} = 1/L$). The selection optimum is $\eta = 1.2$. The same $(\alpha_\ell)_{\ell\in[L]}$ were used in all simulations. The simulations were run for $T = 1000N$ generations, and each observable was measured as an average over the last $500N$ generations.

### 7 Equivalence with the trait's eye-view

Here we recover $\xi_{\Delta_t,\alpha_\ell}$ the selection coefficient of the diffusion equation (5) from the trait's eye-view, as originally proposed by Wright in [47]. This method has seen much use since (see for instance the supplementary data of [39, 43]). We will show that

$$s^0_\ell(\mathbf{P}_t) := 2N\frac{\mathbf{Cov}_{\mathbf{P}_t}[F(Z(g)), g_\ell]}{\mathbf{Var}_{\mathbf{p}}[g_\ell]}$$

satisfies

$$s^0_\ell(\mathbf{P}_t) \simeq \xi_{\Delta_t,\alpha_\ell}(P^\ell_t)$$

under the following hypotheses

(T1) We may write $Z(g) \simeq \mathcal{Z}_{\ell,t} + \alpha_\ell g_\ell - 2\alpha_\ell P^\ell_t$ where under $\mathbf{E}_{\mathbf{P}_t}$, the law of $\mathcal{Z}_{\ell,t}$ is well approximated by a normal distribution $\mathcal{N}(\bar{z}_t, \sigma^2_{\ell,t})$ for some parameters $\bar{z}_t, \sigma^2_{\ell,t}$.

(T2) We have $\sigma_{\ell,t} \ll \omega \qquad \frac{|\Delta_t|}{L} \ll \omega^2$

*Discussion of the hypotheses.* (T1) is interpreted as a consequence of the central limit theorem under HWLE (H1), meaning that if $(\alpha_{\ell'} g_{\ell'})_{\ell' \in [L]}$ are independent under $\mathbf{E}_{\mathbf{P}_t}$, then

$$\mathcal{Z}_{\ell,t} := 2\alpha_\ell P^\ell_t + \sum_{\ell' \in [L] \setminus \{\ell\}} \alpha_{\ell'} g_{\ell'}$$

suitably rescaled converges to a normally distributed variable. This use of the central limit theorem should be interpreted with caution, because **the error on the central limit theorem is of order** $\frac{\sigma_{\ell,t}}{\sqrt{L}}$ (see the Berry-Esseem inequality, for instance Theorem 3, Chapter V of [36], see also the discussion in Appendix C of [3]), which is precisely the order of $\alpha_\ell g_\ell$. Furthermore, the Central Limit Theorem cannot be expected to hold if the genetic variance is too small, that is, $\sigma^2_{\ell,t} \lesssim 1/L^2$. This pathological situation does not occur when (N2) is satisfied, from (35). So the requirement (N2) for the gene's-eye view to hold is also require for the trait's-eye view to hold.

Similarly, (T2) is compatible with (35) when (N1) is satisfied. □

*Alternative derivation of Eq. (6).* Let us define

$$\forall i \in \{0,1,2\}, \qquad w_{\ell,i}(\mathbf{P}_t) := \mathbf{E}_{\mathbf{P}_t}\left[e^{W(\mathcal{Z}_{\ell,t}+\alpha_\ell g_\ell - 2\alpha_\ell P^\ell_t)} \;\middle|\; g_\ell = i\right]$$

Standard computations from population genetics yield

$$s^0_\ell(\mathbf{P}_t) = 2N\frac{(w_{\ell,2}(\mathbf{P}_t) - w_{\ell,1}(\mathbf{P}_t))P^\ell_t + (w_{\ell,1}(\mathbf{P}_t) - w_{\ell,0}(\mathbf{P}_t))(1-P^\ell_t)}{w_{\ell,2}(\mathbf{P}_t)(P^\ell_t)^2 + w_{\ell,1}(\mathbf{P}_t)2P^\ell_t(1-P^\ell_t) + w_{\ell,0}(\mathbf{P}_t)(1-P^\ell_t)^2}. \tag{50}$$

Furthermore, the Gaussian approximation (T1) allows us to compute for $i \in \{0,1,2\}$

$$w_{\ell,i}(\mathbf{P}_t) \propto \exp\left[-\frac{1}{2(\sigma^2_{\ell,t}+\omega^2)}(\alpha_\ell i - 2\alpha_\ell P^\ell_t + \Delta_t)^2\right],$$

where $\Delta_t := \bar{z}_t - \eta$.

We then use (T2) to write $\sigma^2_{\ell,t} + \omega^2 \simeq \omega^2$ and $\alpha_\ell |\Delta_t| \ll \omega^2$. These two approximations yield

$$\begin{aligned} w_{\ell,i}(\mathbf{P}_t) &\propto \; e^{-\frac{\Delta_t^2}{2(\sigma^2_{\ell,t}+\omega^2)}}\left(1 - \frac{1}{2\omega^2}\left((\alpha_\ell i - 2\alpha_\ell P^\ell_t)^2 + 2\Delta_t(\alpha_\ell i - 2\alpha_\ell P^\ell_t)\right)\right) \\ &\propto \; 1 - (i - 2P^\ell_t)\frac{\alpha_\ell}{\omega^2}\Delta_t - \frac{\alpha^2_\ell}{2\omega^2}(i-2P^\ell_t)^2 \end{aligned}$$

Carrying this in (50) we find

$$s^0_\ell(\mathbf{P}_t) \simeq -\frac{\alpha_\ell}{\omega^2_e}\Delta_t - \frac{\alpha^2_\ell}{\omega^2_e}\left(\frac{1}{2} - P^\ell_t\right)$$

We thus recover (6). □

**Remark 3.** *This alternative derivation of the selection coefficient is much more computationally cumbersome, which is why we will use our first approach when discussing extensions of the model in Section 11.*

### 8 When mutations are rare at a given locus

This section answers a reviewer's remark by discussing the link between our model and what is known as the "infinite-sites model". We first clarify the two distinct models which can be labelled "infinite-sites", then discuss the simplifications which they enable.

#### 8.1 Clarification: the two infinite-sites models

From a simulation point of view, one way to implement an infinite-sites model is simply to take large $L$ and small $|\bar{\theta}|$. In this case, almost all copies of a given allele in the population originate from a single mutation. This is the infinite-sites model with back mutations, as used for instance in [6], and which we will focus on. This can be seen as a "house-of-cards" model [42], in which the genetic effect of a mutation at a given locus is 0 or $\alpha_\ell$ with respective probabilities $\theta_\ell^-/|\theta_\ell|$ and $\theta_\ell^+/|\theta_\ell|$.

Another possibility is the following: when a mutation occurs at a locus $\ell$ on an allele with genetic effect $\alpha$, the new allele has effect $\alpha + \tilde{\alpha}$, where $\tilde{\alpha}$ is sampled from some fixed distribution of mutational effects. This is used for instance in [19], and seems to be the default in the SLiM software [18]. This model also underlies the infinite-allele model such as studied in [27, 45]. This is also the underlying model in Fisher's geometric fitness landscape [41, 38], in which the effect of mutations on a multidimensional trait is sampled from a parent-independent fixed distribution. We defer an analysis of this model to further work, but our approach can be extended to it effortlessly.

#### 8.2 Simplifications under rare mutations

Here, we make the following claims

- when $|\bar{\theta}| \ll 1$, the fixed-point equation (14) becomes

$$\Delta^* \simeq 2\,\hat{I}(\Delta^*) - \eta \tag{51}$$

with

$$\hat{I}(\delta) := \mathbb{E}\left[L\alpha \frac{\theta^+ e^{-2\delta\alpha/\omega_e^2}}{\theta^- + \theta^+ e^{-2\delta\alpha/\omega_e^2}}\right],$$

and the equation (31) for the variance $\sigma^2$ becomes

$$\sigma^2 \simeq 2L\mathbb{E}\left[\alpha^2 \frac{2\theta^+\theta^-}{\theta^- + \theta^+ e^{2s^*L\alpha}} \int e^{s^*L\alpha p - \frac{\alpha^2}{\omega_e^2}p(1-p)}\mathrm{d}p\right], \tag{52}$$

- when $|\bar{\theta}| \ll 1$ and $\omega_e^{-2} \ll L^2$ (weak/moderate selection), then the previous equations become

$$\sigma^2 \simeq 2\mathbb{E}\left[\alpha \frac{e^{2s^*L\alpha} - 1}{2s^*} \times \frac{2\theta^+\theta^-}{\theta^- + \theta^+ e^{2s^*L\alpha}}\right], \tag{53}$$

- when $|\bar{\theta}| \ll 1$ and $\omega_e^{-2} \gg L$ (moderate/strong selection), (51) can be further simplified to

$$\hat{I}(\Delta^*) \simeq \eta. \tag{54}$$

As noticed in [12] (Eq. A.2), assuming $(\alpha_\ell, \theta_\ell)$ is constant across loci, this equation can be solved for $s^*$ explicitly to find

$$2s^*L\alpha \simeq \ln\left(\frac{\theta^-}{\theta^+}\right) + \ln\left(\frac{\eta}{2L\alpha - \eta}\right).$$

It should be noted that equations (51-54) are easy to solve numerically, having no pole at $\{0, 1\}$.

The derivation is as follows. When $|\theta_\ell| \ll 1$, the distribution

$$2\frac{\theta_\ell^+\theta_\ell^-}{|\theta_\ell|} p^{2\theta_\ell^+ - 1}(1-p)^{2\theta_\ell^- - 1}\mathrm{d}p$$

is very close to a *Bernoulli*$(\theta_\ell^+/|\theta_\ell|)$ distribution (mathematically speaking, there is weak convergence of the former to the latter when we let $|\theta_\ell| \to 0$ keeping $\theta_\ell^+/|\theta_\ell|$ constant). This leads to a simplified formula for

$\Pi_{\Delta^*,\alpha,\theta}$ in (13)

$$\int p\Pi_{\Delta^*,\alpha,\theta}(p)\mathrm{d}p = \frac{\int_{(0,1)} p^{2\theta^+}(1-p)^{2\theta^- -1} e^{2s^*L\alpha p - \frac{\alpha^2}{\omega_e^2}p(1-p)}\mathrm{d}p}{\int_{(0,1)} p^{2\theta^+ -1}(1-p)^{2\theta^- -1} e^{2s^*L\alpha p - \frac{\alpha^2}{\omega_e^2}p(1-p)}\mathrm{d}p} \simeq \frac{\theta^+ e^{2s^*L\alpha}}{\theta^- + \theta^+ e^{2s^*L\alpha}}.$$

This is known as a Li-Bulmer formula (after [29, 7]), and can also be interpreted as a ratio of the flux of fixation probabilities (see for instance [6]). In particular, we may note that the underdominant term $-\frac{a^2}{\omega_e^2}p(1-p)$ in $\Pi_{\delta,a,\theta}$ can be neglected when most loci are close to fixation due to small mutation rates.

Applying this to fixed-point equation (14), we get (51). If we also have $\omega_e^{-2} \gg L$, then from (35) it holds $\Delta^* \ll 1$. This allows us to neglect $\Delta^*$ in (51), yielding (54).

We turn to $\mathbb{E}^*[\alpha^2 P_t(1-P_t)]$ to get the genetic variance $\sigma^2$ in (31). We find

$$\int p(1-p)\Pi_{\delta,\alpha,\theta}(p)\mathrm{d}p = \frac{\int p^{2\theta^+}(1-p)^{2\theta^-} e^{2s^*L\alpha p - \frac{\alpha^2}{\omega_e^2}p(1-p)}\mathrm{d}p}{\int p^{2\theta^+ -1}(1-p)^{2\theta^- -1} e^{2s^*L\alpha p - \frac{\alpha^2}{\omega_e^2}p(1-p)}\mathrm{d}p} \simeq \frac{2\theta^+\theta^-}{\theta^- + \theta^+ e^{2s^*L\alpha}} \int e^{2s^*L\alpha p - \frac{\alpha^2}{\omega_e^2}p(1-p)}\mathrm{d}p.$$

Carrying this in (31) yields (52). If we also have $\omega_e^{-2} \ll L^2$ (weak/moderate selection), then we may ignore the second term in the exponential and obtain

$$\int p(1-p)\Pi_{\delta,\alpha,\theta}(p)\mathrm{d}p \simeq \frac{2\theta^+\theta^-}{\theta^- + \theta^+ e^{2s^*L\alpha}} \times \frac{e^{2s^*L\alpha} - 1}{2s^*L\alpha}.$$

Carrying this in (31) yields (53).

### 9 Derivation of the criterion for selection to increase genetic variance.

Fix the values of $L, \eta$ of the model. Assume $\theta_\ell = \theta$ and $\alpha_\ell = 1/L$ for some fixed $\theta \in (0,+\infty)^2$. In this section, we will alleviate notation by writing $\mathbb{E}_s$ for the expectation of a typical locus $P_t$ with distribution

$$C_{s,\theta} p^{2\theta^+ -1}(1-p)^{2\theta^- -1} e^{2sp}\mathrm{d}p$$

where $C_{s,\theta}$ is a normalization constant.

Recall the mutational equilibrium

$$z_M := 2L\mathbb{E}_0[P_t]$$

Here we prove the following: assume

$$(1-z_M)(\eta - z_M) > 0 \tag{55}$$

Then there exists an interval $I = (0, \varepsilon]$ such that for any $s \in I$

$$\mathbb{E}_s[P_t(1-P_t)] > \mathbb{E}_0[P_t(1-P_t)]$$

The derivation is as follows. By symmetry, it is enough to treat the case where $1 > z_M$, which is equivalent to

$$2\theta^+ < |\theta| \tag{56}$$

In this case, the criterion 55 translates to $\eta > z_M$. This implies $\Delta^* < 0$, and therefore from (38) $s^* > 0$. We therefore only need to show

$$\left.\frac{\mathrm{d}}{\mathrm{d}s}\right|_{s=0} \mathbb{E}_s[P_t(1-P_t)] > 0$$

Simple computations show

$$\left.\frac{\mathrm{d}}{\mathrm{d}s}\right|_{s=0} \mathbb{E}_s[P_t(1-P_t)] = 2\left(\mathbb{E}_0[P_t^2(1-P_t)] - \mathbb{E}_0[P_t(1-P_t)]\mathbb{E}_0[P_t]\right). \tag{57}$$

Under $\mathbb{E}_0$, $P_t$ has distribution $Beta(2\theta^+, 2\theta^-)$. In particular this yields for any $a, b > 0$

$$\mathbb{E}_0[P_t^a(1-P_t)^b] = \frac{2\theta^+(2\theta^+ + 1)\dots(2\theta^+ + a)2\theta^-(2\theta^- + 1)\dots(2\theta^- + b)}{2|\theta|(2|\theta|+1)\dots(2|\theta|+a+b)}$$

where $|\theta| = \theta^+ + \theta^-$. In particular

$$\begin{aligned}\mathbb{E}_0[P_t] =& \frac{\theta^+}{|\theta|}\\ \mathbb{E}_0[P_t(1-P_t)] =& \frac{2\theta^+ 2\theta^-}{2|\theta|(2|\theta|+1)}\\ \mathbb{E}_0[P_t^2(1-P_t)] =& \frac{2\theta^+(2\theta^+ + 1)2\theta^-}{2|\theta|(2|\theta|+1)(2|\theta|+2)}\end{aligned}$$

It follows from (56)

$$\begin{aligned}\mathbb{E}_0[P_t^2(1-P_t)] - \mathbb{E}_0[P_t(1-P_t)]\mathbb{E}_0[P_t] =& 2\frac{\theta^+\theta^-}{|\theta|^2(2|\theta|+1)(2|\theta|+2)}\left(|\theta|(2\theta^+ + 1) - \theta^+(2|\theta|+2)\right)\\ =& 2\frac{\theta^+\theta^-}{|\theta|^2(2|\theta|+1)(2|\theta|+2)}\left(|\theta| - 2\theta^+\right)\\ >& 0\end{aligned}$$

where we used (56). The result follows, carrying this in (57).

### 10 A first-order correction for the Bulmer effect

Here, we derive a first-order correction for the Bulmer effect using the Quasi-Linkage Equilibrium approach, following [33, 34]. The fit of this correction is shown in Fig. S8, suggesting that the Quasi-Linkage Equilibrium approach is reliable.

#### 10.1 Method

To make the derivation more readable, we will still write $\xi, \Pi, \Delta^*$ for the uncorrected objects defined in (6), (13) and (4), whereas we will define $\hat{\xi}, \hat{\Pi}, \hat{\Delta}^*$ as their corrected counterparts. Crucially, we will denote by $P_t^\ell$ the **uncorrected** frequencies of trait-increasing alleles at locus $\ell$, meaning its stationary distribution is given by $\Pi_{\Delta^*,\alpha_\ell,\theta_\ell}$. On the other hand, we will let $\hat{P}_t^\ell$ be the corrected counterpart to $P_t^\ell$, with stationary distribution given by $\hat{\Pi}_{\hat{\Delta}^*,\alpha_\ell,\theta_\ell}$.

Our derivation relies on the Quasi-Linkage Equilibrium approach [34]. In particular, we consider LD between two loci to be small (but non-negligible) and LD between three loci to be negligible. We also assume Hardy-Weinberg equilibrium, so that $g_\ell$ is still a Binomial if $g$ is a randomly sampled genome. Let us write $D_t^{\ell,\ell'}$ for LD between loci $\ell$ and $\ell'$, such that the covariance between $g_\ell$ and $g_{\ell'}$ for any genome randomly sampled at time $t$ is $2D_t^{\ell,\ell'}$ (the factor 2 comes from the fact that a diploid genome $g$ can be seen as two haploid genomes and that LD only builds up significantly within a haploid genome). In Section 10.2, we show that the selection coefficient acting on locus $\ell$ can be obtained as a function of $(\hat{\mathbf{P}}_t, \mathbf{D}_t)$.

Due to random drift, $(D_t^{\ell,\ell'})_{t\geq 0}$ fluctuates randomly, however if we assume that these fluctuations are fast with respect to the evolution of $\hat{P}_t^\ell$ (which occurs on the order of $2N$ generations), then we may invoke time-averaging as in Section 4.2 and replace $D_t^{\ell,\ell'}$ with its expected value $\mathbb{E}\left[D_t^{\ell,\ell'}|\hat{P}_t^\ell, \hat{P}_t^{\ell'}\right]$. Furthermore, it is a standard computation (see for instance equation (27) in [34], assuming recombination is large and therefore that their $rc_{ij}$ dominates their denominator) that

$$\mathbb{E}^*[D_t^{\ell,\ell'}|\hat{P}_t^\ell = p_\ell, \hat{P}_t^{\ell'} = p_{\ell'}] \simeq -\frac{1}{2r_{\ell,\ell'}\omega_e^2}\alpha_\ell\alpha_{\ell'}p_\ell(1-p_\ell)p_{\ell'}(1-p_{\ell'}), \tag{58}$$

where $r_{\ell_1\ell_2}$ is the rate of recombination between $\ell_1$ and $\ell_2$. In particular, in our system with uniform cross-over $r_{\ell_1\ell_2} = \frac{|\ell_1-\ell_2|}{L}2N$. Let us note (this will be crucial) that the harmonic recombination rate defined with

$$\frac{1}{\bar{r}_h} := \frac{1}{L(L-1)}\sum_{\substack{\ell_1,\ell_2\in[L]\\ \ell_1<\ell_2}}\frac{1}{r_{\ell_1,\ell_2}}$$

satisfies

$$\frac{1}{\bar{r}_h} \simeq \frac{L^2}{2N}\left(\ln(L)+\gamma-1\right), \tag{59}$$

where $\gamma$ is the Euler-Mascheroni constant ($\gamma \simeq .58$). This approximation is correct with order $L/N$.

$$\begin{aligned}
\sum_{\substack{\ell_1,\ell_2\in[L]\\ \ell_1<\ell_2}} \frac{1}{r_{\ell_1,\ell_2}} =& \frac{L}{2N} \sum_{\substack{\ell_1,\ell_2\in[L]\\ \ell_1<\ell_2}} \frac{1}{|\ell_1-\ell_2|}\\
\simeq& \frac{L}{2N}\sum_{\ell_1\in[L]} \ln(L-\ell_1)+\gamma\\
\simeq& \frac{L^2}{2N}\left(\ln(L)+\gamma+\frac{1}{L}\sum_{\ell\in[L]}\ln\left(1-\frac{\ell}{L}\right)\right)\\
\simeq& \frac{L^2}{2N}\left(\ln(L)+\gamma+\int_{(0,1)}\ln(1-x)\,\mathrm{d}x\right)\\
\simeq& \frac{L^2}{2N}\left(\ln(L)+\gamma-1\right),
\end{aligned}$$

where in the second line we used the asymptotic behavior of the harmonic series, and in the fourth line we approximated a Riemann sum with the corresponding integral.

##### 10.2 Correcting the selection coefficient

Here, we show that the selection coefficient at locus $\ell$ corrected for the Bulmer effect should be taken to be

$$\hat{\xi}_{\hat{\Delta}^*,\alpha_\ell}(P_t^\ell) := \xi_{\hat{\Delta}^*,\alpha_\ell}(\hat{P}_t^\ell)+\epsilon \hat{h}_{\hat{\Delta}^*,\alpha_\ell}(\hat{P}_t^\ell), \tag{60}$$

where $\xi$ is defined as before in (6), $\hat{\Delta}^*$ is the deviation to the optimum corrected for LD, and for $\delta\in\mathbb{R}, a\in\mathbb{R}_+$ we define $\hat{h}_{\delta,a}$ with

$$|\bar{\theta}|\hat{h}_{\delta,a}(p) := \frac{a^2}{\omega_e^2}\mathbb{E}^*[(L\alpha)^2P_t(1-P_t)](1-2p)+a\mathbb{E}^*\left[L^2\alpha\left(\theta^+(1-P_t)-\theta^-P_t\right)\right]. \tag{61}$$

and

$$\epsilon := \frac{\ln(L)+\gamma-1}{L\omega^2}|\bar{\theta}|. \tag{62}$$

**Interpretation** Before anything else, one should note from (35) that $\hat{h}_{\Delta^*,\alpha}$ is always of order 1, and therefore that the correction is always of order $\epsilon$. We see from (61) that LD manifests itself as both a positive frequency-dependent effect (analogous to the second term of $\xi$ in (6)) and a directional effect (analogous to the first term of $\xi$). The underdominant effect is classical, same as described for instance in [33]: it can be interpreted as saying LD reduces selection against rare variants with large effects by hiding their effect, linking them with compensating alleles at other loci. The directional effect is of the same form as the effect of mutation on $P_t$ in (7). In particular, it brings the population towards the mutational equilibrium $z_M$ and away from the selection optimum $\eta$. So the conclusion is that the Bulmer effect, due to insufficient recombination, drives the population away from the selection optimum, as can be seen in Fig. 2 from the main text or Fig. S8 below, and as described in [38].

**Derivation of the corrected selection coefficient** We compute $\hat{s}_\ell$ defined analogously to (19)

$$\hat{s}_\ell : \begin{cases} [0,1]^L\times[0,1]^{\binom{[L]}{2}} & \longrightarrow \mathbb{R}\\ \mathbf{p},\mathbf{d} & \longmapsto 2N\frac{\mathbf{Cov}_{\mathbf{p},\mathbf{d}}[W(Z(g)),g_\ell]}{\mathbf{Var}_{\mathbf{p}}[g_\ell]}\end{cases},$$

where $\binom{[L]}{2}$ is the set of pairs of $[L]$, $\mathbf{Cov}_{\mathbf{p},\mathbf{d}}$ is the covariance associated to a random variable $g\in\{0,1,2\}^L$ such that $g_\ell$ has expectation $2p_\ell$ and $(g_\ell,g_{\ell'})$ have covariance $2d_{\ell,\ell'}$. A bias-variance decomposition shows

$$\hat{s}_\ell(\mathbf{p},\mathbf{d}) = -\frac{1}{4\omega_e^2p_\ell(1-p_\ell)}\left(\mathbf{Cov}_{\mathbf{p},\mathbf{d}}[(Z(g)-\bar{Z}(\mathbf{p}))^2,g_\ell]+2\mathbf{Cov}_{\mathbf{p},\mathbf{d}}[Z(g),g_\ell](\bar{Z}(\mathbf{p})-\eta)\right),$$

where $\bar{Z}(\mathbf{p}) = \mathbf{E_p}[Z(g)]$. On one hand, we have

$$\mathbf{Cov_{p,d}}[Z(g), g_\ell] = 2\alpha_\ell p_\ell(1-p_\ell) + 2\sum_{\ell' \in [L]\setminus\{\ell\}} \alpha_{\ell'} d_{\ell,\ell'}.$$

On the other hand, we have

$$\begin{aligned}\mathbf{Cov_{p,d}}[(Z(g)-\bar{Z}(\mathbf{p}))^2, g_\ell] \simeq \alpha_\ell^2 \mathbf{Cov_{p,d}}[(g_\ell - 2p_\ell)^2, g_\ell] + \sum_{\ell' \in [L]\setminus\{\ell\}} \alpha_{\ell'}^2 \mathbf{Cov_{p,d}}[(g_{\ell'} - 2p_{\ell'})^2, g_\ell] \\ + 2\sum_{\ell' \in [L]\setminus\{\ell\}} \alpha_\ell \alpha_{\ell'} \mathbf{Cov_{p,d}}[(g_\ell - 2p_\ell)(g_{\ell'} - 2p_{\ell'}), g_\ell],\end{aligned}$$

where we neglected third-order linkage of the form

$$(g_{\ell_1} - 2p_{\ell_1})(g_{\ell_2} - 2p_{\ell_2})(g_{\ell_3} - 2p_{\ell_3}).$$

Tedious computations yield

$$\begin{aligned}\mathbf{Cov_{p,d}}[(g_\ell - 2p_\ell)^2, g_\ell] &= 2p_\ell(1-p_\ell)(1-2p_\ell) \\ \mathbf{Cov_{p,d}}[(g_{\ell'} - 2p_{\ell'})^2, g_\ell] &= 2(1-2p_{\ell'})d_{\ell,\ell'} \\ \mathbf{Cov_{p,d}}[(g_\ell - 2p_\ell)(g_{\ell'} - 2p_{\ell'}), g_\ell] &= 2(1-2p_\ell)d_{\ell,\ell'}.\end{aligned}$$

It follows

$$\mathbf{Cov_{p,d}}[(Z(g)-\bar{Z}(\mathbf{p}))^2, g_\ell] \simeq 2\alpha_\ell\,(1-2p_\ell)\left(\alpha_\ell p_\ell(1-p_\ell) + 2\sum_{\ell' \in [L]\setminus\{\ell\}} \alpha_{\ell'} d_{\ell',\ell}\right) + 2\sum_{\ell' \in [L]\setminus\{\ell\}} \alpha_{\ell'}^2 (1-2p_{\ell'}) d_{\ell,\ell'}.$$

This yields

$$\begin{aligned}\hat{s}_\ell(\mathbf{p},\mathbf{d}) = -\frac{\alpha_\ell}{2\omega_e^2}(1-2p_\ell)\left(\alpha_\ell + 2\sum_{\ell' \in [L\setminus\{\ell\}]} \alpha_{\ell'} \frac{d_{\ell,\ell'}}{p_\ell(1-p_\ell)}\right) \\ + \frac{1}{2\omega_e^2}\sum_{\ell' \in [L]\setminus\{\ell\}} \alpha_{\ell'}^2 (1-2p_{\ell'}) \frac{d_{\ell,\ell'}}{p_\ell(1-p_\ell)} \\ - \frac{1}{\omega_e^2}\left(\alpha_\ell + \sum_{\ell' \in [L]\setminus\{\ell\}} \alpha_{\ell'} \frac{d_{\ell,\ell'}}{p_\ell(1-p_\ell)}\right)\left(\bar{Z}(\mathbf{p}) - \eta\right).\end{aligned}$$

Using the time-averaging assumption on $d_{\ell,\ell'}$ from Section 10.1, we find from (58) that $\hat{s}_\ell(\hat{\mathbf{P}}_t, \mathbf{D}_t)$ can be approximated with

$$\begin{aligned}-\frac{\alpha_\ell^2}{2\omega_e^2}(1-2\hat{P}_t^\ell)\left(1 - \sum_{\ell' \in [L\setminus\{\ell\}]} \frac{\alpha_{\ell'}^2 L}{\omega^2|\ell-\ell'|}\hat{P}_t^{\ell'}(1-\hat{P}_t^{\ell'})\right) \\ - \frac{\alpha_\ell}{4\omega_e^2}\sum_{\ell' \in [L]\setminus\{\ell\}} \frac{\alpha_{\ell'}^3 L}{\omega^2|\ell-\ell'|}(1-2\hat{P}_t^{\ell'})\hat{P}_t^{\ell'}(1-\hat{P}_t^{\ell'}) \\ - \frac{\alpha_\ell}{\omega_e^2}\left(1 - \sum_{\ell' \in [L]\setminus\{\ell\}} \frac{\alpha_{\ell'}^2 L}{2\omega^2|\ell-\ell'|}\hat{P}_t^{\ell'}(1-\hat{P}_t^{\ell'})\right)\hat{\Delta}_t.\end{aligned}$$

Then, we use the following mean-field approximations (the validity of which would deserve further scrutiny)

$$\begin{aligned}\sum_{\ell' \in [L\setminus\{\ell\}]} \frac{\alpha_{\ell'}^2 L}{\omega^2|\ell-\ell'|}\hat{P}_t^{\ell'}(1-\hat{P}_t^{\ell'}) &\simeq 2L(\ln(L)+\gamma-1)\mathbb{E}^*\left[\frac{\alpha^2}{\omega^2}P_t(1-P_t)\right] \\ \sum_{\ell' \in [L]\setminus\{\ell\}} \frac{\alpha_{\ell'}^3 L}{\omega^2|\ell-\ell'|}(1-2\hat{P}_t^{\ell'})\hat{P}_t^{\ell'}(1-\hat{P}_t^{\ell'}) &\simeq 2L(\ln(L)+\gamma-1)\mathbb{E}^*\left[\frac{\alpha^3}{\omega^2}(1-2P_t)P_t(1-P_t)\right],\end{aligned}$$

where $2(\ln(L)+\gamma-1)$ comes from summing up over $[L]\setminus\{\ell\}$ the $1/|\ell-\ell'|$, and replacing the resulting local harmonic recombination rate with its mean value given in (59). This only holds approximately for loci which are not too close to the boundary $\{1, L\}$. Note in particular that this mean-field approximation replaces the corrected $\hat{P}_t$ with its uncorrected counterpart $P_t$, which is acceptable at first-order.

We therefore obtain that the corrected selection coefficient should be

$$\xi_{\hat{\Delta}_t,\alpha_\ell}(\hat{P}^\ell_t) + \epsilon h_{\hat{\Delta}_t,\alpha_\ell}(\hat{P}^\ell_t),$$

with $\epsilon$ defined in (62) and

$$h_{\delta,a}(p) := \frac{a^2}{\omega_e^2}\mathbb{E}^*[(L\alpha)^2 P_t(1-P_t)](1-2p) + \frac{a}{\omega_e^2}\omega_e^{-2}\left(-\frac{1}{2}\mathbb{E}^*[L^2\alpha^3(1-2P_t)P_t(1-P_t)] + \delta\mathbb{E}^*\left[(L\alpha)^2 P_t(1-P_t)\right]\right).$$

To get the result, first we replace $\hat{\Delta}_t$ with its expected value $\hat{\Delta}^* := \mathbb{E}\left[\hat{\Delta}^*\right]$, as in Section 4. Second, we use that $h$ is itself small, and that to a first order $\hat{\Delta}^* \simeq \Delta^*$ to write $h_{\hat{\Delta}^*,\alpha_\ell} \simeq h_{\Delta^*,\alpha_\ell}$. Finally, to get (60), we only have to show that $h_{\Delta^*,\alpha_\ell} = \hat{h}_{\Delta^*,\alpha_\ell}$, where $\hat{h}$ is defined in (61). That is, we need to show

$$\omega_e^{-2}\left(-\frac{1}{2}\mathbb{E}^*[L^2\alpha^3(1-2P_t)P_t(1-P_t)] + \delta\mathbb{E}^*\left[(L\alpha)^2 P_t(1-P_t)\right]\right) = \mathbb{E}^*\left[L^2\alpha\left(\theta^+(1-P_t) - \theta^- P_t\right)\right]. \tag{63}$$

This follows from the equation of $P_t$ in (7): which we recall

$$\mathrm{d}P_t = \omega_e^{-2}\left(-\alpha\Delta^* + \frac{\alpha^2}{2}(2P_t - 1)\right)P_t(1-P_t)\,\mathrm{d}t + \left(\theta^+(1-P_t) - \theta^- P_t\right)\mathrm{d}t + \sqrt{P_t(1-P_t)}\,\mathrm{d}B^P_t.$$

Multiplying by $L^2\alpha$ and taking the expectation at equilibrium we find

$$0 = \omega_e^{-2}\mathbb{E}^*\left[\left(-(L\alpha)^2\Delta^* + \frac{L^2\alpha^3}{2}(2P_t - 1)\right)P_t(1-P_t)\right] + \mathbb{E}^*\left[L^2\alpha\left(\theta^+(1-P_t) - \theta^- P_t\right)\right].$$

This yields (63).

##### 10.3 Correcting the stationary distribution

Here, we claim that the corrected stationary distribution at locus $\ell$ has density $\hat{\Pi}_{\hat{\Delta}^*,\alpha_\ell,\theta_\ell}$, where for $\delta \in \mathbb{R}, a \in \mathbb{R}_+, \theta \in (0,+\infty)^2$ we have

$$\hat{\Pi}_{\delta,a,\theta}(p) \simeq \Pi_{\delta,a,\theta}(p)\left(1 + 2\epsilon\left(\hat{H}_{\delta,a}(p) - \int \hat{H}_{\delta,a}(p')\Pi_{\delta,a,\theta}(p')\mathrm{d}p'\right)\right), \tag{64}$$

where $\hat{H}_{\delta,a}$ is a primitive of $\hat{h}_{\delta,a}$ from (61), given by

$$|\bar{\theta}|\hat{H}_{\delta,a}(p) := \frac{a^2}{\omega_e^2}\mathbb{E}^*[(L\alpha)^2 P_t(1-P_t)]p(1-p) + a\mathbb{E}^*\left[L^2\alpha\left(\theta^+(1-P_t) - \theta^- P_t\right)\right]p. \tag{65}$$

The derivation is straightforward, we define $\hat{\Pi}$ analogously to $\Pi$ in (13) with $\hat{\xi}$ instead of $\xi$, and use a first-order expansion in $\epsilon$.

##### 10.4 Correcting the fixed-point equation

Here, we claim that the corrected deviation to the optimum $\hat{\Delta}^*$ satisfies

$$\hat{\Delta}^* \simeq \Delta^* + \frac{4\epsilon}{1 - 2I'(\Delta^*)}L\mathbb{E}\left[\alpha\int p\left(\hat{H}_{\Delta^*,\alpha}(p) - \int \hat{H}_{\Delta^*,\alpha}(p')\Pi_{\Delta^*,\alpha,\theta}(p')\mathrm{d}p'\right)\Pi_{\Delta^*,\alpha,\theta}(p)\mathrm{d}p\right]. \tag{66}$$

**Interpretation** The integral in (66) corresponds to the covariance, for $p$ sampled with law $\Pi_{\Delta^*,\alpha,\theta}$ with $\hat{H}_{\Delta^*,\alpha}(p)$. If $\Pi_{\Delta^*,\alpha,\theta}$ is symmetric, then the covariance of $p$ with $p(1-p)$ is zero and therefore only the second term of $\hat{H}_{\Delta^*,\alpha}$ in (65) comes into play: the correction is then proportional to the variance of $p$ and in particular is positive. This means that LD leads to an increase in mutational bias.

**Derivation** The derivation is as follows.

First, from (64), the mean trait value is given by

$$\begin{aligned}2L\mathbb{E}^*\left[\alpha\hat{P}_t\right] =&2L\mathbb{E}\left[\alpha\int p\hat{\Pi}_{\hat{\Delta}^*,\alpha,\theta}(p)\mathrm{d}p\right]\\ \simeq&2L\mathbb{E}\left[\alpha\int p\left(1+2\epsilon\left(\hat{H}_{\hat{\Delta}^*,\alpha}(p)-\int\hat{H}_{\hat{\Delta}^*,\alpha}(p')\Pi_{\hat{\Delta}^*,\alpha,\theta}(p')\mathrm{d}p'\right)\right)\Pi_{\hat{\Delta}^*,\alpha,\theta}(p)\mathrm{d}p\right].\end{aligned}$$

It follows that the corrected fixed-point equation is

$$\begin{aligned}\hat{\Delta}^* \simeq&2L\mathbb{E}\left[\alpha\int p\left(1+2\epsilon\left(\hat{H}_{\hat{\Delta}^*,\alpha}(p)-\int\hat{H}_{\hat{\Delta}^*,\alpha}(p')\Pi_{\hat{\Delta}^*,\alpha,\theta}(p')\mathrm{d}p'\right)\right)\Pi_{\hat{\Delta}^*,\alpha,\theta}(p)\mathrm{d}p\right]-\eta\\ \simeq&2I(\hat{\Delta}^*)-\eta+4\epsilon L\mathbb{E}\left[\alpha\int p\left(\hat{H}_{\hat{\Delta}^*,\alpha}(p)-\int\hat{H}_{\hat{\Delta}^*,\alpha}(p')\Pi_{\hat{\Delta}^*,\alpha,\theta}(p')\mathrm{d}p'\right)\Pi_{\hat{\Delta}^*,\alpha,\theta}(p)\mathrm{d}p\right].\end{aligned}$$

Let us write $\hat{\Delta}^* = \Delta^* + c_\Delta$ for some small correction $c_\Delta$. Using that $\Delta^*$ satisfies the fixed-point equation (14) and the first-order expansion $I(\hat{\Delta}^*) \simeq I(\Delta^*) + c_\Delta I'(\Delta^*)$, we find

$$c_\Delta \simeq c_\Delta 2I'(\Delta^*)+4\epsilon L\mathbb{E}\left[\alpha\int p\left(\hat{H}_{\hat{\Delta}^*,\alpha}(p)-\int\hat{H}_{\hat{\Delta}^*,\alpha}(p')\Pi_{\hat{\Delta}^*,\alpha,\theta}(p')\mathrm{d}p'\right)\Pi_{\hat{\Delta}^*,\alpha,\theta}(p)\mathrm{d}p\right].$$

This yields (66).

#### 10.5 Correcting the genetic variance

We first recall that when LD is non-negligible, one should differentiate between the genetic variance $\sigma_G^2$, defined as the variance of $Z(g)$ where $g$ is a genome randomly sampled within the population, and the genic variance $\sigma_g^2$ defined as the trait variance we would have under HWLE. In particular, a mean-field approximation as in (31) shows that the corrected genic variance $\hat{\sigma}_g^2$ satisfies

$$\hat{\sigma}_g^2 \simeq 2L\mathbb{E}^*[\alpha^2\hat{P}_t(1-\hat{P}_t)].$$

We claim

$$\begin{aligned}\hat{\sigma}_g^2 \simeq&\sigma^2+\frac{4c_\Delta}{\omega_e^2}L\mathbb{E}\left[\alpha^3\left(\int p(1-p)\left(p-\int p'\Pi_{\Delta^*,\alpha,\theta}(p')\mathrm{d}p'\right)\Pi_{\Delta^*,\alpha,\theta}(p)\mathrm{d}p\right)\right]\\ &+4\epsilon L\mathbb{E}\left[\alpha^2\int p(1-p)\left(\hat{H}_{\Delta^*,\alpha}(p)-\int\hat{H}_{\Delta^*,\alpha}(p')\Pi_{\Delta^*,\alpha,\theta}(p')\mathrm{d}p'\right)\Pi_{\Delta^*,\alpha,\theta}(p)\mathrm{d}p\right]\\ \hat{\sigma}_G^2 \simeq&\left(1-\frac{\sigma^2}{\omega^2}(\ln(L)-1+\gamma)\right)\hat{\sigma}_g^2,\end{aligned}$$

where $c_\Delta := \hat{\Delta}^* - \Delta^*$ is determined by (66).

The derivation is as follows. The formula for $\hat{\sigma}_g^2$ is obtained in two steps. First, a straightforward application of (64) yields

$$\begin{aligned}\hat{\sigma}_g^2 \simeq 2L\mathbb{E}^*&\left[\alpha^2\int p(1-p)\Pi_{\hat{\Delta}^*,\alpha,\theta}(p)\mathrm{d}p\right]\\ &+4\epsilon L\mathbb{E}\left[\alpha^2\int p(1-p)\left(\hat{H}_{\hat{\Delta}^*,\alpha}(p)-\int\hat{H}_{\hat{\Delta}^*,\alpha}(p')\Pi_{\hat{\Delta}^*,\alpha,\theta}(p')\mathrm{d}p'\right)\Pi_{\hat{\Delta}^*,\alpha,\theta}(p)\mathrm{d}p\right].\end{aligned}$$

Second, we replace $\hat{\Delta}^*$ with $\Delta^* + c_\Delta$ in $\Pi_{\hat{\Delta}^*,\alpha,\theta}$ and use a Taylor expansion in $c_\Delta$.

The second formula is well-known since [8], and can be derived as follows. We have from (58) that

$$\begin{aligned}\hat{\sigma}_G^2 =&\sum_{\ell\in[L]}2\alpha_\ell^2\hat{P}_t^\ell(1-\hat{P}_t^\ell)+2\sum_{\substack{\ell,\ell'\in[L]\\ \ell<\ell'}}\alpha_\ell\alpha_{\ell'}2D_t^{\ell,\ell'}\\ \simeq&2L\mathbb{E}^*[\alpha^2\hat{P}_t(1-\hat{P}_t)]-2\sum_{\substack{\ell,\ell'\in[L]\\ \ell<\ell'}}\alpha_\ell^2\alpha_{\ell'}^2\frac{L}{|\ell-\ell'|\omega^2}\hat{P}_t^\ell(1-P_t^\ell)\hat{P}_t^{\ell'}(1-P_t^{\ell'})\\ \simeq&2L\mathbb{E}^*[\alpha^2\hat{P}_t(1-\hat{P}_t)]-\frac{4L^2(\ln(L)+\gamma-1)}{\omega^2}\mathbb{E}^*[\alpha^2\hat{P}_t(1-\hat{P}_t)]^2,\end{aligned}$$

where in the last line we used a mean-field approximation and (59). We therefore get the result.

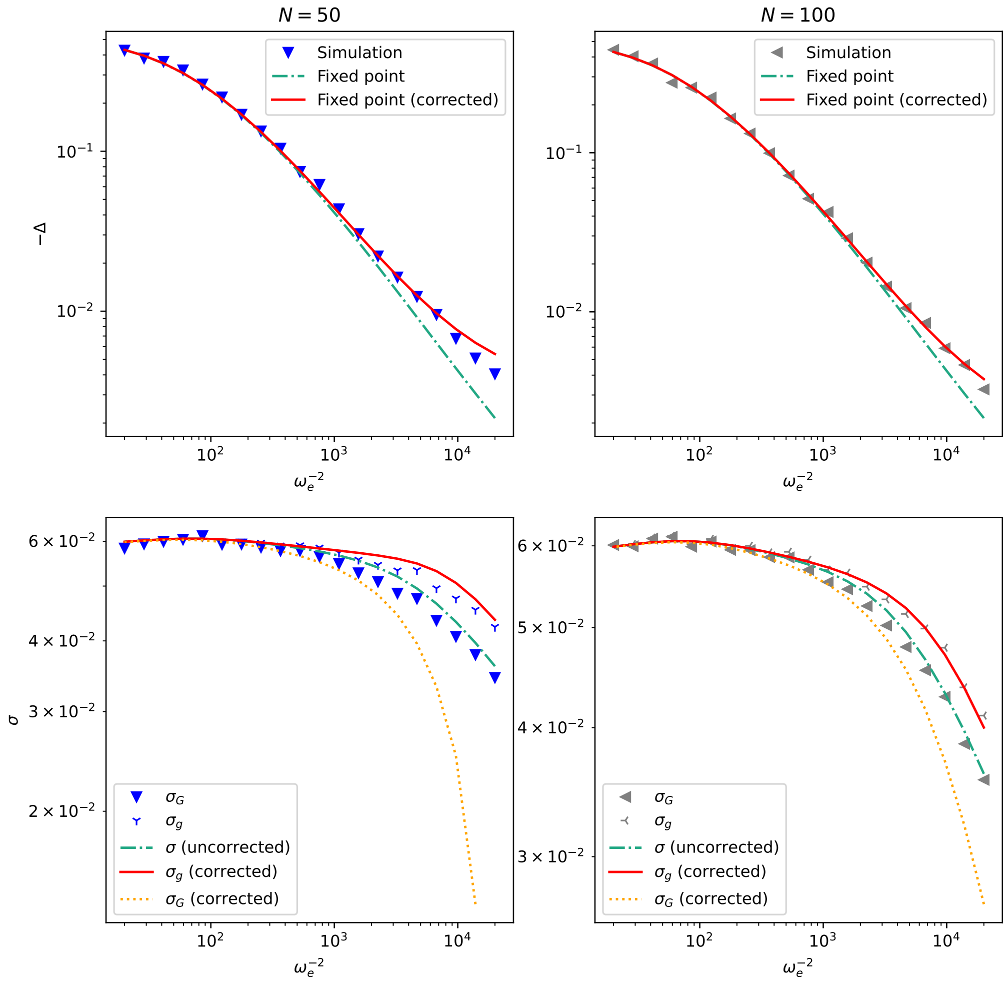


Figure 8: Simulated populations with $L = 100, \theta = (.1, .2), \eta = 1.2$, and $\alpha$ exponentially distributed and renormalized. The lines show the corrections for the Bulmer effect just derived.

### 11 Some extensions

Here we show how our computations can be extended to handle special cases. In Section 11.1 we mention polyploidy, in Section 11.2 pleiotropy, in Section 11.3 dominance. We devote the whole Section 12 to discuss epistasis. We always start from an analog of the diffusion approximation (5), with selection coefficient given by (19). In each extension, the analog of assumptions (A1-6) and (N1-3) is assumed to hold.

#### 11.1 Polyploidy

For a polyploid (or haploid) population in which each organism has $k$ haploid genomes with $k \geq 1$, the following alterations need to be made to the diffusion equation (5)

- For $\mathbf{p} \in [0,1]^L$, $g_\ell$ under $\mathbf{E_p}$ has law $Binomial(k, p_\ell)$.
- In (5), the definition of $\omega_e$ is $\omega_e^{-2} := (kN)\omega^{-2}$, and the definition of $\theta_\ell$ is $\theta_\ell := kN\mu_\ell$.

In this case the computations of (6) yield

$$s_\ell(\mathbf{p}) = \frac{\alpha_\ell}{\omega^2}\left(\eta - k\sum_{\ell' \in [L]} \alpha_{\ell'} p_{\ell'}\right) + \frac{\alpha_\ell^2}{\omega^2}\left(p_\ell - \frac{1}{2}\right)$$

All of the previous results can be obtained from this, defining

$$\Delta_t = k\sum_{\ell' \in [L]} \alpha_{\ell'} P_t^{\ell'} - \eta$$

#### 11.2 Pleiotropy

We give here an outlook as to how the theory can be adapted to account for pleiotropy, deferring to future work a rigorous treatment of edge cases.

We now assume that $\alpha_\ell = (\alpha_\ell^i)_{i\in[d]}$ takes values in $\mathbb{R}^d$ with $d$ fixed. In particular, we allow $\alpha_\ell^i < 0$. This is because when $d = 1$, up to replacing $(\alpha_\ell, (\theta_\ell^+, \theta_\ell^-), P_t^\ell)$ with $(-\alpha_\ell, (\theta_\ell^-, \theta_\ell^+), 1 - P_t^\ell)$, we may always assume $\alpha_\ell > 0$, but that is no longer true when $\alpha_\ell$ has dimension $d$.

In this setting, $Z(g) = \sum_\ell \alpha_\ell g_\ell$ is a $d-$dimensional vector encoding $d$ traits. The fitness function is given by

$$\forall z \in \mathbb{R}^d, \qquad W(z) = -\frac{1}{2}(z-\eta)^\top \omega^{-2}(z-\eta)$$

where $^\top$ denotes transposition, $\omega^2$ is a positive definite matrix and $\eta \in \mathbb{R}^d$. The same computations as in Section 3.2 show that the selection coefficient at locus $\ell$ is $\xi_{\Delta_t,\alpha_\ell}(P_t^\ell)$ where $\Delta_t$ is as in (4) and for $a, \delta \in \mathbb{R}^d$ we define

$$\xi_{\delta,a} : \begin{cases} [0,1] & \longrightarrow \mathbb{R} \\ p & \longmapsto -a^\top \omega_e^{-2}\delta \ + \ a^\top \omega_e^{-2} a \left(p - \frac{1}{2}\right) \end{cases} \tag{67}$$

with $\omega_e^{-2} := 2N\omega^{-2}$.

Using the same kind of reasoning as in Section 4.1, we argue that we may write

$$\xi_{\Delta_t,\alpha_\ell}(P_t^\ell) \simeq \xi_{\Delta^*,\alpha_\ell}(P_t^\ell) \tag{68}$$

where $\Delta^* := \mathbb{E}^*[\Delta_t]$.

It follows that $P_t^\ell$ is evolves according to a frequency-dependent Wright-Fisher diffusion (3) with selection coefficient $\xi_{\Delta^*,\alpha_\ell}$. In particular, $P_t^\ell$ has stationary distribution $\Pi_{\Delta^*,\alpha_\ell,\theta_\ell}$ where $\Pi$ is defined as in (13) replacing the original definition of $\xi$ by its multi-dimensional counterpart (67). We can then find the $d$-dimensional fixed point equation for $\Delta^*$ as

$$\Delta^* = 2L\mathbb{E}\left[\int \alpha p \Pi_{\Delta^*,\alpha,\theta}(p)\mathrm{d}p\right] - \eta \tag{69}$$

Global observables such as the genetic variance-covariance matrix $\sigma^2 = 2L\mathbb{E}^*[\alpha\alpha^\top P_t(1-P_t)]$ can be obtained from this as in Section 5.2.

Future work could tackle the analysis of this system when $d \gg 1$ as as was explored in [39]. We could also investigate the effect of the variance-covariance matrix of the $(\alpha^i)_{i\in[d]}$, in particular when some traits are very correlated. We also note that proving existence and uniqueness of solutions to (69) will require more work than what we did in the single-trait case in Section 2.1.3.

#### 11.3 Dominance

We introduce the model in Section 11.3.1, compute the selection coefficient in Section 11.3.2, the evolution of the trait under moderate/strong selection ($\omega_e^{-2} \gg L$) in Section 11.3.4, and obtain the fixed point equation for $\Delta^*$ and the stationary distribution of the typical locus in Section 11.3.3.

##### 11.3.1 Model

We now account for dominance. Locus $\ell$ is now characterized by allelic effect $\alpha_\ell$, dominance effect $D_\ell$ and mutation rate $\theta_\ell$. The trait value and logfitness of a genotype $g = (g_\ell)_{\ell \in [L]}$ is then

$$Z(g) = \sum_{\ell \in [L]} \alpha_\ell g_\ell + D_\ell \mathbb{1}_{[g_\ell = 1]} \qquad\qquad W(z) = -\frac{1}{2\omega^2}(z-\eta)^2$$

For a pair $(a, D) \in \mathbb{R}_+ \times \mathbb{R}$ we define the corresponding **average effect of gene substitution** as

$$\beta_{a,D}(p) := a + D(1-2p) \tag{70}$$

##### 11.3.2 Selection coefficient

Here we show the selection coefficient $s_\ell(\mathbf{p})$ defined from (19) can be rewritten $s_\ell(\mathbf{P}_t) = \xi_{\Delta_t, \alpha_\ell, D_\ell}(P_t^\ell)$ where for $\delta \in \mathbb{R}, a \in \mathbb{R}_+, D \in \mathbb{R}$ we define

$$\xi_{\delta,a,D}(p) := -\frac{\beta_{a,D}(p)}{\omega_e^2}\delta + \omega_e^{-2}\left(\beta_{a,D}(p)^2\left(p - \frac{1}{2}\right) + 2aDp(1-p)\right) \tag{71}$$

$$\Delta_t := \mathbf{E}_{\mathbf{P}_t}[Z(g)] - \eta \;=\; 2\sum_{\ell \in [L]} \left(\alpha_\ell P_t^\ell + D_\ell P_t^\ell (1 - P_t^\ell)\right) \;-\; \eta. \tag{72}$$

*Derivation.* For a general $\mathbf{p} \in [0,1]^L$ we write

$$\begin{aligned}\mathbf{Cov_p}[(Z(g)-\eta)^2, g_\ell] =& \mathbf{Cov_p}[\mathbf{E_p}[(Z(g)-\eta)^2 \,|g_\ell], g_\ell] \\ =& \mathbf{Cov_p}[\mathbf{E_p}[Z(g)-\eta \,|g_\ell]^2, g_\ell] + \mathbf{Cov_p}[\mathbf{Var_p}[Z(g)-\eta \,|g_\ell], g_\ell]\end{aligned}$$

We can write

$$\mathbf{Var_p}[Z(g)-\eta \,|g_\ell] = \sum_{\ell' \in [L] \setminus \{\ell\}} \mathbf{Var_p}[\alpha_{\ell'} g_{\ell'} + D_{\ell'} \mathbb{1}_{[g_{\ell'}=1]}]$$

In particular

$$\mathbf{Cov_p}[\mathbf{Var_p}[Z(g)-\eta \,|g_\ell], g_\ell] = 0.$$

We are then left with

$$\mathbf{Cov_p}[(Z(g)-\eta)^2, g_\ell] = \mathbf{Cov_p}[\mathbf{E_p}[Z(g)-\eta \,|g_\ell]^2, g_\ell].$$

Appyling this to $\mathbf{P}_t$, we get

$$\mathbf{E}_{\mathbf{P}_t}[Z(g)-\eta \,|g_\ell] = \Delta_t + \alpha_\ell(g_\ell - 2P_t^\ell) + D_\ell(\mathbb{1}_{[g_\ell=1]} - 2P_t^\ell(1-P_t^\ell))$$

using the definition of $\Delta_t$ in (72). It follows

$$\begin{aligned}\mathbf{Cov}_{\mathbf{P}_t}[(Z(g)-\eta)^2, g_\ell] = \mathbf{Cov}_{\mathbf{P}_t}\Big[&\alpha_\ell^2(g_\ell - 2P_t^\ell)^2 + D_\ell^2(\mathbb{1}_{[g_\ell=1]} - 2P_t^\ell(1-P_t^\ell))^2 \\ &+ 2\alpha_\ell(g_\ell - 2P_t^\ell)D_\ell(\mathbb{1}_{[g_\ell=1]} - 2P_t^\ell(1-P_t^\ell)) \\ &+ 2\Delta_t\left(\alpha_\ell(g_\ell - 2P_t^\ell) + D_\ell(\mathbb{1}_{[g_\ell=1]} - 2P_t^\ell(1-P_t^\ell))\right), g_\ell\Big].\end{aligned}$$

We compute

$$\begin{aligned}\mathbf{Cov}_{\mathbf{P}_t}[(g_\ell - 2P_t^\ell)^2, g_\ell] =& (1-2P_t^\ell)2P_t^\ell(1-P_t^\ell) \\ \mathbf{Cov}_{\mathbf{P}_t}[\mathbb{1}_{[g_\ell=1]}, g_\ell] =& (1-2P_t^\ell)2P_t^\ell(1-P_t^\ell) \\ \mathbf{Cov}_{\mathbf{P}_t}[\mathbb{1}_{[g_\ell=1]}g_\ell, g_\ell] =& \mathbf{Cov}_{\mathbf{P}_t}[\mathbb{1}_{[g_\ell=1]}, g_\ell]\end{aligned}$$

In particular

$$\begin{aligned}\mathbf{Cov}_{\mathbf{P}_t}[(g_\ell - 2P_t^\ell)(\mathbb{1}_{[g_\ell=1]} - 2P_t^\ell(1-P_t^\ell)), g_\ell] =& \mathbf{Cov}_{\mathbf{P}_t}[(1-2P_t^\ell)\mathbb{1}_{[g_\ell=1]} - 2P_t^\ell(1-P_t^\ell)g_\ell, g_\ell] \\ =& (1-2P_t^\ell)^2 2P_t^\ell(1-P_t^\ell) - \left(2P_t^\ell(1-P_t^\ell)\right)^2.\end{aligned}$$

It follows

$$\mathbf{Cov}_{\mathbf{P}_t}[(Z(g)-\eta)^2, g_\ell] = \alpha_\ell^2(1-2P_t^\ell)2P_t^\ell(1-P_t^\ell) + D_\ell^2(1-2P_t^\ell)^3 2P_t^\ell(1-P_t^\ell)$$
$$+ 2\alpha_\ell D_\ell \left((1-2P_t^\ell)^2 2P_t^\ell(1-P_t^\ell) - (2P_t^\ell(1-P_t^\ell))^2\right)$$
$$+ 2\Delta_t\left(\alpha_\ell + D_\ell(1-2P_t^\ell)\right) 2P_t^\ell(1-P_t^\ell).$$

This lets us obtain

$$\frac{\mathbf{Cov}_{\mathbf{P}_t}[(Z(g)-\eta)^2, g_\ell]}{2P_t^\ell(1-P_t^\ell)} = \left(\alpha_\ell + D_\ell(1-2P_t^\ell)\right)^2(1-2P_t^\ell) - 4\alpha_\ell D_\ell P_t^\ell(1-P_t^\ell) + 2(\alpha_\ell + D_\ell(1-2P_t^\ell))\Delta_t$$

We get from (70)

$$s_\ell(\mathbf{P}_t) = \omega_e^{-2}\Big(\beta_{\alpha_\ell,D_\ell}(P_t^\ell)^2\left(P_t^\ell - \frac{1}{2}\right) + 2\alpha_\ell D_\ell P_t^\ell(1-P_t^\ell) - \beta_{\alpha_\ell,D_\ell}(P_t^\ell)\Delta_t\Big)$$

The result follows. □

##### 11.3.3 Genetic architecture

The same argument as in Section 4.2 lets us write

$$\xi_{\Delta_t,\alpha_\ell,D_\ell}(P_t^\ell) \simeq \xi_{\Delta^*,\alpha_\ell,D_\ell}(P_t^\ell)$$

where $\Delta^* = \mathbb{E}^*[\Delta_t]$. Then $P_t^\ell$ follows a frequency-dependent Wright-Fisher diffusion (3) with selection coefficient $\xi_{\Delta^*,\alpha_\ell,D_\ell}$. It follows that $P_t^\ell$ has stationary distribution $\Pi_{\Delta^*,\alpha_\ell,D_\ell,\theta_\ell}$ where we define for $\delta \in \mathbb{R}, a \in \mathbb{R}_+, D \in \mathbb{R}, \theta \in (0,+\infty)^2$

$$\Pi_{\delta,a,D,\theta}(p) := C_{\delta,a,D,\theta}\, p^{2\theta^+-1}(1-p)^{2\theta^--1}e^{2\int_0^p \xi_{\delta,a,D}(u)\,\mathrm{d}u} \tag{73}$$

with $C_{\delta,a,D,\theta}$ a normalization constant.

On the other hand we can compute from (72)

$$\Delta^* \simeq 2L\mathbb{E}^*[\alpha P_t + DP_t(1-P_t)] - \eta$$

From there we obtain the fixed point equation

$$\Delta^* \simeq 2I(\Delta^*) - \eta \tag{74}$$

where

$$I(\delta) := \mathbb{E}^*\left[\int L(\alpha p + Dp(1-p))\Pi_{\delta,\alpha,D,\theta}(p)\mathrm{d}p\right].$$

The macroscopic observables can be computed from this as in Section 5.2. In particular, let us show the genetic variance is

$$\sigma^2 = 2L\mathbb{E}^*\left[\beta_{\alpha,D}(P_t)^2 P_t(1-P_t)\right] + 2L\mathbb{E}^*\left[2D^2\left(P_t(1-P_t)\right)^2\right]. \tag{75}$$

The first term is the **additive genetic variance**, the second term is the **dominance variance**.

Finally, we note that when mutation rates are small ($|\bar{\theta}| \ll 1$), as in Section 8 we may assume that $\Pi_{\delta,\alpha,D,\theta}$ is close to a Bernoulli distribution, and in particular the distribution of $P_t$ is very concentrated near the boundary $\{0,1\}$. If we further assume that selection is moderate or weak ($\omega_e^{-2} \ll L^2$), then it can be checked that $\Pi_{\delta,\alpha,D,\theta}$ is approximately independent of $D$, and is close to a

$$\mathit{Bernoulli}\left(\frac{\theta^+ e^{-2\delta\alpha/\omega_e^2}}{\theta^- + \theta^+ e^{-2\delta\alpha/\omega_e^2}}\right).$$

Carrying this in the fixed-point equation (74), we recover (51), and in particular we see that dominance has no effect on the stationary distribution of $P_t$ and the value of $\Delta^*$. It does however have an impact on $\sigma^2$.

*Derivation of* (75). From the new definition of $Z(g)$ and the independence of the $(g_\ell)_{\ell\in[L]}$, we have

$$\mathbf{Var}_{\mathbf{P}_t}[Z(g)] = \sum_{\ell\in[L]} \mathbf{Var}_{\mathbf{P}_t}[\alpha_\ell g_\ell + D_\ell \mathbb{1}_{[g_\ell=1]}]$$

We compute

$$\mathbf{Var}_{\mathbf{P}_t}[\alpha_\ell g_\ell + D_\ell \mathbb{1}_{[g_\ell=1]}] = \alpha_\ell^2 \mathbf{Var}_{\mathbf{P}_t}[g_\ell] + D_\ell^2 \mathbf{Var}_{\mathbf{P}_t}[\mathbb{1}_{[g_\ell=1]}] + 2\alpha_\ell D_\ell \mathbf{Cov}_{\mathbf{P}_t}[g_\ell, \mathbb{1}_{[g_\ell=1]}]$$
$$= 2\alpha_\ell^2 P_t^\ell(1-P_t^\ell) + D_\ell^2 2P_t^\ell(1-P_t^\ell)\left(1-2P_t^\ell(1-P_t^\ell)\right) + 2\alpha_\ell D_\ell \mathbf{Cov}_{\mathbf{P}_t}[g_\ell, \mathbb{1}_{[g_\ell=1]}] \quad (76)$$

where in the second equality we used that $g_\ell$ is a $Binomial(2, P_t^\ell)$ variable and $\mathbb{1}_{[g_\ell=1]}$ is a $Bernoulli(2P_t^\ell(1-P_t^\ell))$ variable.

Using $\mathbb{1}_{[g_\ell=1]} g_\ell = \mathbb{1}_{[g_\ell=1]}$, we write

$$\mathbf{Cov}_{\mathbf{P}_t}[g_\ell, \mathbb{1}_{[g_\ell=1]}] = \mathbf{E}_{\mathbf{P}_t}[\mathbb{1}_{[g_\ell=1]}] - \mathbf{E}_{\mathbf{P}_t}[g_\ell]\mathbf{E}_{\mathbf{P}_t}[\mathbb{1}_{[g_\ell=1]}]$$
$$= 2P_t^\ell(1-P_t^\ell)\left(1-2P_t^\ell\right)$$

Carrying this into (76) we find

$$\mathbf{Var}_{\mathbf{P}_t}[\alpha_\ell g_\ell + D_\ell \mathbb{1}_{[g_\ell=1]}] = 2\alpha_\ell^2 P_t^\ell(1-P_t^\ell) + D_\ell^2 2P_t^\ell(1-P_t^\ell)\left(1-2P_t^\ell(1-P_t^\ell)\right) + 2\alpha_\ell D_\ell \times 2P_t^\ell(1-P_t^\ell)\left(1-2P_t^\ell\right)$$

which can be rewritten

$$\mathbf{Var}_{\mathbf{P}_t}[\alpha_\ell g_\ell + D_\ell \mathbb{1}_{[g_\ell=1]}] = 2P_t^\ell(1-P_t^\ell)\left(\alpha_\ell^2 + 2\alpha_\ell D_\ell(1-2P_t^\ell) + D_\ell^2(1-2P_t^\ell(1-P_t^\ell))\right).$$

Using the definition of $\beta$ in (70) we find

$$\mathbf{Var}_{\mathbf{P}_t}[\alpha_\ell g_\ell + D_\ell \mathbb{1}_{[g_\ell=1]}] = 2P_t^\ell(1-P_t^\ell)\left(\beta_{\alpha_\ell,D_\ell}(P_t^\ell)^2 + D_\ell^2\left(1-2P_t^\ell(1-P_t^\ell) - (1-2P_t^\ell)^2\right)\right)$$

which yields the result. □

##### 11.3.4 Evolution of the trait under moderate/strong selection

Here obtain the following stationary system analog to Eq. (7-9)

$$\mathrm{d}P_t = \xi_{\Delta^*,\alpha,D}(P_t)P_t(1-P_t)\,\mathrm{d}t + \left(\theta^+(1-P_t) - \theta^- P_t\right)\mathrm{d}t + \sqrt{P_t(1-P_t)}\,\mathrm{d}B_t^P$$
$$\Delta^* = 2L\mathbb{E}^*[\alpha P_t + DP_t(1-P_t)] - \eta$$
$$\mathrm{d}\varepsilon_t = -\rho\varepsilon_t\,\mathrm{d}t + \omega_e\sqrt{2\rho}\,\mathrm{d}B_t^\Delta$$

where $B^P, B^\Delta$ are Brownian motions, $\varepsilon_t := \Delta_t - \Delta^*$ and

$$\rho := \frac{1}{\tau} \times \frac{\mathbb{E}^*\left[2(L\beta_{\alpha,D}(P_t))^2 P_t(1-P_t)\right]}{|\bar{\theta}|}$$

*Derivation.* Applying Itô's formula yields

$$\mathrm{d}\Delta_t = 2\sum_{\ell\in[L]}\left(\alpha_\ell + D_\ell(1-2P_t^\ell)\right)\mathrm{d}P_t^\ell - \sum_{\ell\in[L]} 2D_\ell P_t^\ell(1-P_t^\ell)\mathrm{d}t$$
$$= 2\sum_{\ell\in[L]}\beta_{\alpha_\ell,D_\ell}(P_t^\ell)\mathrm{d}P_t^\ell - \sum_{\ell\in[L]} 2D_\ell P_t^\ell(1-P_t^\ell)\mathrm{d}t$$

We define as before

$$\tau := \frac{L\omega_e^2}{|\bar{\theta}|}$$

We can then proceed just as in Section 4, using mean-field approximations and time-averaging to write

$$\mathrm{d}\Delta_t = \frac{1}{\tau}\Bigg( -\Delta_t \times \frac{1}{|\bar{\theta}|} \times \mathbb{E}^*\left[2(L\beta_{\alpha,D}(P_t))^2 P_t(1-P_t)\right] + \frac{L\omega_e^2}{|\bar{\theta}|} \times \mathbb{E}^*\left[2L\beta_{\alpha,D}(P_t)\left(\theta^+(1-P_t) - \theta^- P_t\right)\right]$$
$$+ \frac{1}{|\bar{\theta}|} \times \mathbb{E}^*\left[2L^2\beta_{\alpha,D}(P_t)^3\left(P_t - \frac{1}{2}\right)P_t(1-P_t) + 4L^2\alpha D\beta_{\alpha,D}(P_t)P_t^2(1-P_t)^2\right] - \frac{L\omega_e^2}{|\bar{\theta}|}\mathbb{E}^*[2LDP_t(1-P_t)]\Bigg)\mathrm{d}t$$
$$+ \frac{1}{\sqrt{\tau}} \times \sqrt{\frac{\omega_e^2}{|\bar{\theta}|}\mathbb{E}^*\left[(2L\beta_{\alpha,D}(P_t))^2 P_t(1-P_t)\right]}\mathrm{d}B_t^\Delta \quad (77)$$

The result follows. □

### 12 Epistasis

Adding epistasis to the model means the trait $Z(g)$ involves interactions between loci. Some work has already been done on the subject, for instance [44, 4]. There are many ways such interactions can go, depending on assumptions about gene regulatory networks (the typical model for such networks is the NK fitness landscape [23]). We will not go in-depth, but briefly state how the fixed-point equation can be adapted to two extreme cases.

Let $(\alpha_{\ell,\ell'})_{1\le\ell<\ell'\le L}$ be i.i.d random variables and for $\ell<\ell'$, define $\alpha_{\ell',\ell}:=\alpha_{\ell,\ell'}$. Assume that $L$ is even (this assumption is made to simplify the model). Consider the two trait functions

$$Z_{sc}(g) := \sum_{1\le\ell\le L/2} \alpha_{2\ell-1,2\ell}\, g_{2\ell-1} g_{2\ell} \tag{78}$$

$$Z_{sg}(g) := \sum_{1\le\ell<\ell'\le L} \alpha_{\ell,\ell'}\, g_\ell g_{\ell'} \tag{79}$$

The first model $Z_{sc}$ considers that we can partition $[L]$ into $L/2$ pairs of interacting loci. This is a special case of a general model we will refer to as the *small clumps* model, in which $[L]$ is partitioned into small clumps of interacting loci. In terms of the NK fitness model, this corresponds to a small value of K, that is, each locus interacts with a small number of other loci, in such a way that if $\ell$ interacts with $\ell'$ and $\ell'$ interacts with $\ell''$, then $\ell$ interacts with $\ell''$.

The second model $Z_{sg}$, inspired by the Sherrington-Kirkpatrick model from *spin-glass* theory [28], considers that all loci interact loosely with each other. In terms of the NK model, this corresponds to K=N-1, that is, each locus interacts with every other.

As in Section 3 the diffusion approximation under LE is given by

$$\mathrm{d}P_t^\ell = s_\ell(\mathbf{P}_t)P_t^\ell(1-P_t^\ell)\mathrm{d}t + (\theta_\ell^+(1-P_t^\ell) - \theta_\ell^- P_t^\ell)\mathrm{d}t + \sqrt{P_t^\ell(1-P_t^\ell)}\mathrm{d}B_t^\ell \tag{80}$$

where (under HWLE) we have

$$s_\ell(\mathbf{P}_t) := 2N\frac{\mathbf{Cov}_{\mathbf{P}_t}[W(g), g_\ell]}{\mathbf{Var}_{\mathbf{P}_t}[g_\ell]} \tag{81}$$

with $W(g) = -\frac{1}{2\omega^2}(Z(g)-\eta)^2$ and $Z$ the chosen trait function.

We start with a reminder on diffusions and potentials in Section 12.1, then consider the "small clumps" model in Section 12.2, and the "general diffuse interactions" model in Section 12.3.

#### 12.1 Reminder on diffusions and potentials

Consider a neutral Wright-Fisher diffusion on $[0,1]^L$

$$\forall \ell\in[L],\quad \mathrm{d}X_t^\ell = (\theta_\ell^+(1-X_t^\ell) - \theta_\ell^- X_t^\ell)\mathrm{d}t + \sqrt{X_t^\ell(1-X_t^\ell)}\mathrm{d}B_t^\ell.$$

for independent Brownian motions $(B^\ell)_{\ell\in[L]}$. The stationary distribution of $(X_t^\ell)_{\ell\in[L]}$ has density on $[0,1]^L$ given by

$$C\prod_{\ell\in[L]}(p_\ell^{2\theta_\ell^+-1}(1-p_\ell)^{2\theta_\ell^--1}).$$

Now, we want to add a potential to the system, that is, we fix a function $F:[0,1]^L\to\mathbb{R}$ and consider the new equation

$$\forall \ell\in[L],\quad \mathrm{d}Y_t^\ell = (\partial_\ell F(\mathbf{Y}_t))Y_t^\ell(1-Y_t^\ell)\mathrm{d}t + (\theta_\ell^+(1-Y_t^\ell) - \theta_\ell^- Y_t^\ell)\mathrm{d}t + \sqrt{Y_t^\ell(1-Y_t^\ell)}\mathrm{d}B_t^\ell.$$

In this equation, the first term corresponds to the **gradient of the potential**. The stationary distribution of $(Y_t^\ell)_{\ell\in[L]}$ then has the density

$$C\prod_{\ell\in[L]}(p_\ell^{2\theta_\ell^+-1}(1-p_\ell)^{2\theta_\ell^--1})\exp(2F(\mathbf{p})).$$

In this section, we will want to apply this property in two settings. The first is the most natural example of potential given by selection of the form (81), which we summarize in the following claim.

**Claim 12.1** (Wright's formula [46])**.** *If $s_\ell$ is of the form* (81)*, then $s_\ell$ corresponds to the gradient of the potential defined as $F(\mathbf{p}) = N\mathbf{E}_\mathbf{p}[W(g)]$. Formally,*

$$\forall \ell\in[L],\qquad s_\ell(\mathbf{p}) = N\partial_\ell\mathbf{E}_\mathbf{p}[W(g)]. \tag{82}$$

*In particular, the stationary distribution of* $(P_t^\ell)_{\ell\in[L]}$ *in* (80) *has density given by*

$$C \prod_{\ell\in[L]} (p_\ell^{2\theta_\ell^+-1}(1-p_\ell)^{2\theta_\ell^--1})\ \exp\left(2N\mathbf{E_p}[W(g)]\right). \tag{83}$$

*for some normalization constant* $C$.

*Derivation of Claim 12.1.* This is known as Wright's formula [46]. We simply write

$$\mathbf{E_p}[W(g)] = (1-p_\ell)^2\mathbf{E_p}[W(g)|g_\ell=0] + 2p_\ell(1-p_\ell)\mathbf{E_p}[W(g)|g_\ell=1] + p_\ell^2\mathbf{E_p}[W(g)|g_\ell=2].$$

From this and the fact that $\partial_\ell\mathbf{E_p}[W(g)|g_\ell=i]=0$ for $i\in\{0,1,2\}$ we get

$$\begin{aligned}
\partial_\ell\mathbf{E_p}[W(g)] =& -2(1-p_\ell)\mathbf{E_p}[W(g)|g_\ell=0] + 2(1-2p_\ell)\mathbf{E_p}[W(g)|g_\ell=1] + 2p_\ell\mathbf{E_p}[W(g)|g_\ell=2]\\
=&\frac{1}{p_\ell(1-p_\ell)}\big((1-p_\ell)^2\mathbf{E_p}[W(g)|g_\ell=0](0-2p_\ell) + 2p_\ell(1-p_\ell)\mathbf{E_p}[W(g)|g_\ell=1](1-2p_\ell)\\
&\qquad + p_\ell^2\mathbf{E_p}[W(g)|g_\ell=2](2-2p_\ell)\big)\\
=&\frac{\mathbf{E_p}[W(g)(g_\ell-2p_\ell)]}{p_\ell(1-p_\ell)}\\
=&\frac{\mathbf{Cov_p}[W(g),g_\ell]}{p_\ell(1-p_\ell)}\\
=&\frac{s_\ell(\mathbf{p})}{N}.
\end{aligned}$$

□

The second use of a potential will require the following crucial result stated without proof

**Lemma 12.2** (Poincaré's lemma (See e.g Lemma 4-1.2 in [16]))**.** *Consider a differentiable function* $\mathbf{f}\equiv(f_\ell)_{\ell\in[L]}$ *on* $[0,1]^L$*. Suppose for any* $\ell,\ell'\in[L]$ *with* $\ell\neq\ell'$ *we have*

$$\partial_\ell f_{\ell'} = \partial_{\ell'} f_\ell \tag{84}$$

*Then* $\mathbf{f}$ *is the gradient of the potential defined as*

$$F(\mathbf{p}) := \int_0^1 \mathbf{s}(t\mathbf{p})\cdot\mathbf{p}\mathrm{d}t$$

*where* $\cdot$ *is the standard scalar product on* $\mathbb{R}^{[L]}$.

#### 12.2 Epistasis between small clumps of loci

##### 12.2.1 The base model

We start with the model $Z_{sc}$ defined in (78), before generalizing. We consider locus $2\ell$ for $\ell\in[L/2]$. We can write the selection coefficient at locus $2\ell$ as in (19)

$$s_{2\ell}(\mathbf{p}) = -\frac{1}{2\omega_e^2}\times\frac{\mathbf{Cov_p}\left[(Z_{sc}(g)-\eta)^2, g_{2\ell}\right]}{2p_{2\ell}(1-p_{2\ell})}$$

with $\omega_e^{-2} := 2N\omega^{-2}$ the effective strength of selection. Using the independence of the $(g_\ell)_{\ell\in[L]}$ under HWLE, this can be rewritten splitting the term of $Z_{sc}$ depending on $g_{2\ell}$ and the rest

$$s_{2\ell}(\mathbf{p}) = -\frac{\mathbf{Cov_p}\left[\alpha_{2\ell-1,2\ell}^2\left(g_{2\ell-1}g_{2\ell}-\mathbf{E_p}\left[g_{2\ell-1}g_{2\ell}\right]\right)^2, g_{2\ell}\right] + 2\mathbf{Cov_p}\left[\alpha_{2\ell-1,2\ell}g_{2\ell-1}g_{2\ell}, g_{2\ell}\right]\mathbf{E_p}\left[Z_{sc}(g)\ -\eta\right]}{4\omega_e^2 p_{2\ell}(1-p_{2\ell})}.$$

Defining as above $\Delta_t := \mathbf{E}_{\mathbf{P}_t}\left[Z_{sc}(g)\ -\eta\right]$ we get

$$s_{2\ell}(\mathbf{P}_t) = -\frac{\mathbf{Cov}_{\mathbf{P}_t}\left[\alpha_{2\ell-1,2\ell}^2\left(g_{2\ell-1}g_{2\ell}-\mathbf{E}_{\mathbf{P}_t}\left[g_{2\ell-1}g_{2\ell}\right]\right)^2, g_{2\ell}\right] + 2\mathbf{Cov_P}\left[\alpha_{2\ell-1,2\ell}g_{2\ell-1}g_{2\ell}, g_{2\ell}\right]\Delta_t}{4\omega_e^2 P_t^{2\ell}(1-P_t^{2\ell})}.$$

We rewrite this as $s_{2\ell}(\mathbf{P}_t) = \tilde{s}_{\alpha_{2\ell-1,2\ell},\Delta_t}((P_t^{2\ell-1},P_t^{2\ell}))$ where for $\alpha,\delta\in\mathbb{R}$ and $\mathbf{p}\equiv(p_1,p_2)\in[0,1]^2$ we define

$$\tilde{s}_{\alpha,\delta}(\mathbf{p}) := -\frac{\mathbf{Cov_p}\left[\alpha^2\left(g_1g_2-\mathbf{E_p}\left[g_1g_2\right]\right)^2, g_1\right] + 2\mathbf{Cov_P}\left[\alpha g_1g_2, g_1\right]\delta}{4\omega_e^2 p_1(1-p_1)}$$

where, for $\mathbf{p} \in [0,1]^2$, $\mathbf{E_p}$ (resp. $\mathbf{Cov_p}$) is the expectation (resp. covariance) of a $\{0,1,2\}^2$-valued vector $g$ of independent random variables such that $g_\ell$ has law $Binomial(2, p_\ell)$. Because $Z_{sc}$ is symetric in $(g_{2\ell-1}, g_{2\ell})$ we also find that $s_{2\ell-1}(\mathbf{P}_t) = \tilde{s}_{\alpha_{2\ell-1,2\ell},\Delta_t}((P_t^{2\ell}, P_t^{2\ell-1}))$.

Let $(\tilde{P}_t^1, \tilde{P}_t^2, \tilde{\theta}_1, \tilde{\theta}_2, \tilde{\alpha})$ be equal to $(P_t^{2\ell_U-1}, P_t^{2\ell_U}, \theta_{2\ell_U-1}, \theta_{2\ell_U}, \alpha_{2\ell_U-1,2\ell_U})$ for $\ell_U$ sampled uniformly at random on $[L/2]$. This represents the typical clump of loci. We can write the dynamics of $(\tilde{P}_t^1, \tilde{P}_t^2)$ assuming $\Delta_t \simeq \Delta^* := \mathbb{E}^*[\Delta_t]$ as the following autonomous system

$$\begin{aligned}
\mathrm{d}\tilde{P}_t^1 &= \tilde{s}_{\tilde{\alpha},\Delta^*}(\tilde{P}_t^2, \tilde{P}_t^1)\tilde{P}_t^1(1-\tilde{P}_t^1)\mathrm{d}t + (\tilde{\theta}_1^+(1-\tilde{P}_t^1) - \tilde{\theta}_1^- \tilde{P}_t^1)\mathrm{d}t + \sqrt{\tilde{P}_t^1(1-\tilde{P}_t^1)}\mathrm{d}\tilde{B}_t^1 \\
\mathrm{d}\tilde{P}_t^2 &= \tilde{s}_{\tilde{\alpha},\Delta^*}(\tilde{P}_t^1, \tilde{P}_t^2)\tilde{P}_t^2(1-\tilde{P}_t^2)\mathrm{d}t + (\tilde{\theta}_2^+(1-\tilde{P}_t^2) - \tilde{\theta}_2^- \tilde{P}_t^2)\mathrm{d}t + \sqrt{\tilde{P}_t^2(1-\tilde{P}_t^2)}\mathrm{d}\tilde{B}_t^2.
\end{aligned}$$

As in Section 4, we can then find the stationary distribution of $(\tilde{P}_t^1, \tilde{P}_t^2)$ and a fixed point equation for $\Delta^*$. To do this, we must first check that the function

$$(p_1, p_2) \mapsto (\tilde{s}_{\tilde{\alpha},\Delta^*}(p_1, p_2), \tilde{s}_{\tilde{\alpha},\Delta^*}(p_1, p_2))$$

is the gradient of a potential.

For this particular example, we will in fact prove a stronger result. We will find a fitness function $W_{\alpha,\Delta_t}$ such that

$$\tilde{s}_{\tilde{\alpha},\Delta^*}(\mathbf{p}) = 2N \frac{\mathbf{Cov_p}\left[W_{\tilde{\alpha},\Delta_t}(g_1, g_2), g_1\right]}{2p_1(1-p_2)}$$

so that Wright's formula (Claim 12.1) yields the desired result. Tedious computations show that the following definition of $W_{\alpha,\Delta^*}$ yields the desired result

$$\forall \alpha, \delta \in \mathbb{R}, g_1, g_2 \in \{0,1,2\}, \quad W_{\alpha,\delta}(g_1, g_2) := -\frac{\alpha^2}{2\omega^2} g_1 g_2 \left(g_1 g_2 - 4(g_1 - 1)(g_2 - 1)\right) - \frac{\delta g_1 g_2}{\omega^2}.$$

From this, we obtain the density of the stationary distribution of $(\tilde{P}_t^1, \tilde{P}_t^2)$ as $\Pi_{W_{\tilde{\alpha},\Delta^*},\tilde{\Theta}}$ where $\tilde{\Theta} := (\tilde{\theta}_1, \tilde{\theta}_2)$ and we define for $\mathbf{p} = (p_1, p_2) \in [0,1]^2$, $W$ a function on $\{0,1,2\}^2$ and $\Theta = (\theta_1^+, \theta_1^-, \theta_2^+, \theta_2^-) \in \mathbb{R}_+^4$

$$\Pi_{W,\Theta}(\mathbf{p}) := C(p_1^{2\theta_1^+-1}(1-p_1)^{2\theta_1^--1})(p_2^{2\theta_2^+-1}(1-p_2)^{2\theta_2^--1}) \exp\left(4N\mathbf{E_p}[W(g_1, g_2)]\right). \tag{85}$$

The fixed point equation for $\Delta^*$ is then

$$\Delta^* = 4L\mathbb{E}\left[\tilde{\alpha} \int\int p_1 p_2 \Pi_{W_{\tilde{\alpha},\Delta^*},\tilde{\Theta}}(p_1, p_2)\mathrm{d}p_1\mathrm{d}p_2\right] - \eta. \tag{86}$$

##### 12.2.2 General model with small clumps of loci

Under the assumption of small clumps of loci, we assume there exists a partition of $[L]$ into small subsets $(\mathcal{I}_k)_{k\in[n]}$ such that the trait value $Z(g)$ is the sum from contributions from each subset

$$Z(g) = \sum_{k=1}^{n} Z_k(g_{\mathcal{I}_k}). \tag{87}$$

where $g_{\mathcal{I}_k} := (g_\ell)_{\ell\in\mathcal{I}_k}$ and $Z_k$ is a function on $\{0,1,2\}^{\mathcal{I}_k}$. We assume that there are many clumps of loci ($n \gg 1$), and that for each $k$, $Z_k$ is independently sampled as a random function on $\{0,1,2\}^{\mathcal{I}_k}$ with a distribution that only depends on $\#\mathcal{I}_k$. For instance, we could consider

$$Z_k(g) = \sum_{\mathcal{J}\subseteq\mathcal{I}_k} \alpha_{\mathcal{J}} \prod_{\ell\in\mathcal{J}} g_\ell$$

where the $(\alpha_{\mathcal{J}})_{\mathcal{J}\subseteq\mathcal{I}_k}$ have some distribution. In this subsection we derive the fixed-point equation for $\Delta^*$ in this system.

We can write the selection coefficient at locus $\ell \in \mathcal{I}_k$ as in (19)

$$s_\ell(\mathbf{p}) = -\frac{1}{2\omega_e^2} \times \frac{\mathbf{Cov_p}\left[(Z(g)-\eta)^2, g_\ell\right]}{2p_\ell(1-p_\ell)}$$

Using the independence of $g_{\mathcal{I}_k}$ and $(g_{\mathcal{I}_{k'}})_{k'\neq k}$, this can be rewritten as above

$$s_\ell(\mathbf{P}_t) = -\frac{\mathbf{Cov}_{\mathbf{P}_t}\left[\left(Z_k(g_{\mathcal{I}_k}) - \mathbf{E}_{\mathbf{P}_t}\left[Z_k(g_{\mathcal{I}_k})\right]\right)^2, g_\ell\right] + 2\mathbf{Cov}_{\mathbf{P}_t}\left[Z_k(g_{\mathcal{I}_k}), g_\ell\right]\Delta_t}{4\omega_e^2 P_t^\ell(1-P_t^\ell)}.$$

where $\Delta_t := \mathbf{E}_{\mathbf{P}_t}\left[Z(g) - \eta\right]$.

Let us now assume (this is derived at the end of the section) that this can be rewritten as the gradient of a potential

**Claim 12.3.** *For each $k \in [n]$ and $\delta \in \mathbb{R}$, we can find a potential function $F_{k,\delta}$ on $[0,1]^{\mathcal{I}_k}$ such that for any $\mathbf{p} \in [0,1]^L$ such that $\delta = \mathbf{E_p}[Z(g) \ - \eta]$, we have*

$$\forall \ell \in \mathcal{I}_k, \qquad s_\ell(\mathbf{p}) = \partial_\ell F_{k,\delta}(\mathbf{p}_{\mathbf{I}_k})$$

*where $\mathbf{p}_{\mathcal{I}_k} = (p_\ell)_{\ell \in \mathcal{I}_k}$.*

If we assume $\Delta_t \simeq \Delta^*$, we can then rewrite the dynamics at clump $\mathcal{I}_k$ as an autonomous multiloci Wright-Fisher diffusion

$$\forall \ell \in \mathcal{I}_k, \quad \mathrm{d}P_t^\ell = \partial_\ell F_{k,\Delta^*}(\mathbf{P}_t^{\mathcal{I}_k}) P_t^\ell (1-P_t^\ell)\mathrm{d}t + (\theta_\ell^+(1-P_t^\ell) - \theta_\ell^- P_t^\ell)\mathrm{d}t + \sqrt{P_t^\ell(1-P_t^\ell)}\mathrm{d}B_t^\ell$$

In particular, the stationary density of $(P_t^\ell)_{\ell \in \mathcal{I}_k}$ can be obtained as

$$\forall \mathbf{p} \in [0,1]^{\mathcal{I}_k}, \qquad \Pi_{F_{k,\Delta^*},\theta_{\mathcal{I}_k}}(\mathbf{p}) = C_{F_{k,\Delta^*},\theta_{\mathcal{I}_k}} \prod_{\ell \in \mathcal{I}_k} p_\ell^{2\theta_\ell^+ - 1}(1-p_\ell)^{2\theta_\ell^- - 1} \exp\left[2F_{k,\Delta^*}(\mathbf{p})\right]$$

where $C_{F_{k,\Delta^*},\theta_{\mathcal{I}_k}}$ is a normalization constant. Taking a clump uniformly at random exactly as in the previous section, we may obtain from (87) a fixed point equation for $\Delta^*$ as

$$n\mathbb{E}\left[\int_{[0,1]^{\mathcal{I}_k}} \mathbf{E_p}[Z_k(g)]\Pi_{F_{k_U,\Delta^*},\theta_{\mathcal{I}_{k_U}}}(\mathbf{p})\mathrm{d}\mathbf{p}\right] - \eta = \Delta^*$$

where, for $\mathbf{p} \in [0,1]^{\mathcal{I}_k}$, $\mathbf{E_p}$ is the expectation of a $\{0,1,2\}^{\mathcal{I}_k}$-valued vector $g$ of independent random variables such that $g_\ell$ has law $Binomial(2, p_\ell)$, and $k_U$ is a uniform random variable on $[n]$.

*Derivation of Claim 12.3.* For $\ell \in \mathcal{I}_k$, $\delta \in \mathbb{R}$ and $\mathbf{p} \in [0,1]^{\mathcal{I}_k}$, we define

$$\tilde{s}_{\ell,\Delta_t}(\mathbf{p}) := -\frac{\mathbf{Cov_p}\left[\left(Z_k(g_{\mathcal{I}_k}) - \mathbf{E_p}\left[Z_k(g_{\mathcal{I}_k})\right]\right)^2, g_\ell\right] + 2\mathbf{Cov_p}\left[Z_k(g_{\mathcal{I}_k}), g_\ell\right]\Delta_t}{4\omega_e^2 p_\ell(1-p_\ell)}.$$

To apply Poincaré's lemma (Lemma 12.2), we must show (84), that is, that for any $\ell, \ell' \in \mathcal{I}_k$ with $\ell \neq \ell'$ we have

$$\partial_\ell \tilde{s}_{\ell',\delta} = \partial_{\ell'}\tilde{s}_{\ell,\delta}$$

We start by rewriting $\tilde{s}_{\ell,\delta}$, opening the square and rearranging the terms as

$$\tilde{s}_{\ell,\delta}(\mathbf{p}) = \frac{2\mathbf{E_p}\left[Z_k(g_{\mathcal{I}_k})(g_\ell - 2p_\ell)\right]\mathbf{E_p}\left[Z_k(g_{\mathcal{I}_k})\right] - \mathbf{E_p}\left[\left(Z_k(g_{\mathcal{I}_k})^2 + 2Z_k(g_{\mathcal{I}_k})\Delta_t\right)(g_\ell - 2p_\ell)\right]}{4\omega_e^2 p_\ell(1-p_\ell)}$$

Using the same computation as in the warm-up we can show that for any $\ell \in [L]$ and function $f$ on $\{0,1,2\}^L$ we have

$$\begin{aligned}\partial_\ell \mathbf{E_p}[f(g)] &= \frac{\mathbf{Cov_p}[f(g), g_\ell]}{2p_\ell(1-p_\ell)} \\ &= \frac{\mathbf{E_p}[f(g)(g_\ell - 2p_\ell)]}{p_\ell(1-p_\ell)}\end{aligned}$$

Applying this to $\tilde{s}_{\ell,\delta}(\mathbf{p})$

$$\begin{aligned}\partial_{\ell'}\tilde{s}_{\ell,\delta}(\mathbf{p}) = \frac{1}{4\omega_e^2 p_\ell(1-p_\ell)p_{\ell'}(1-p_{\ell'})} &\left(2\mathbf{E_p}\left[Z_k(g_{\mathcal{I}_k})(g_\ell - 2p_\ell)(g_{\ell'} - 2p_{\ell'})\right]\mathbf{E_p}\left[Z_k(g_{\mathcal{I}_k})\right]\right. \\ &+ 2\mathbf{E_p}\left[Z_k(g_{\mathcal{I}_k})(g_\ell - 2p_\ell)\right]\mathbf{E_p}\left[Z_k(g_{\mathcal{I}_k})(g_{\ell'} - 2p_{\ell'})\right] \\ &\left. - \mathbf{E_p}\left[\left(Z_k(g_{\mathcal{I}_k})^2 + 2Z_k(g_{\mathcal{I}_k})\Delta_t\right)(g_\ell - 2p_\ell)(g_{\ell'} - 2p_{\ell'})\right]\right)\end{aligned}$$

This is symmetric in $\ell, \ell'$, (84) follows. □

#### 12.3 Diffuse epistasis between all loci

##### 12.3.1 The base model

Here we assume $Z$ takes the form $Z_{sg}$ in (78). We start by noticing that the independence of $g_\ell$ and $(g_{\ell'})_{\ell' \in [L] \setminus \{\ell\}}$ yields the decomposition

$$\mathbf{Cov_p}\left[(Z(g)-\eta)^2, g_\ell\right] = \mathbf{Cov_p}\left[\left(Z(g) - \mathbf{E_p}\left[Z(g)\,|g_\ell\right]\right)^2, g_\ell\right] + \mathbf{Cov_p}\left[\left(\mathbf{E_p}\left[Z(g)\,|g_\ell\right] - \eta\right)^2, g_\ell\right] \tag{88}$$

which lets us write

$$s_\ell(\mathbf{P}_t) = -\frac{\epsilon_\ell(t)}{2\omega_e^2} - \frac{\mathbf{Cov}_{\mathbf{P}_t}\left[\left(\mathbf{E}_{\mathbf{P}_t}\left[Z(g)\,|g_\ell\right] - \mathbf{E}_{\mathbf{P}_t}\left[Z(g)\right]\right)^2, g_\ell\right] + 2\mathbf{Cov}_{\mathbf{P}_t}\left[Z(g), g_\ell\right]\Delta_t}{4\omega_e^2 P_t^\ell(1-P_t^\ell)}.$$

where $\Delta_t := \mathbf{E}_{\mathbf{P}_t}\left[Z(g)\ -\eta\right]$ and we define

$$\epsilon_\ell(t) := \frac{\mathbf{Cov}_{\mathbf{P}_t}\left[\left(Z(g) - \mathbf{E}_{\mathbf{P}_t}\left[Z(g)\,|g_\ell\right]\right)^2, g_\ell\right]}{2P_t^\ell(1-P_t^\ell)}.$$

Standard computations (same as in Section 3.2) allow us to obtain

$$s_\ell(\mathbf{P}_t) = -\frac{\epsilon_\ell(t)}{2\omega_e^2} + \frac{\hat{\alpha}_\ell(t)^2}{\omega_e^2}\left(P_t^\ell - \frac{1}{2}\right) - \frac{\hat{\alpha}_\ell(t)\Delta_t}{\omega_e^2}.$$

where

$$\hat{\alpha}_\ell(t) := \sum_{\ell' \neq \ell} \alpha_{\ell,\ell'} 2P_t^{\ell'}. \tag{89}$$

Let us define the typical locus $(P_t, \hat{\alpha}^*, \theta)$ as $(P_t^{\ell_U}, \hat{\alpha}^*_{\ell_U}, \theta_{\ell_U})$ for $\ell_U$ picked uniformly at random on $[L]$. The equation for the typical locus becomes tractable under the three following approximations

- $|\epsilon_\ell(t)| \ll \omega_e^2$ (discussed in Claim 12.4)
- a mean-field approximation on $\hat{\alpha}_\ell(t)$ lets us write $\hat{\alpha}_\ell(t) \simeq \hat{\alpha}^*_\ell := 2L\mathbb{E}\left[\alpha\right]\mathbb{E}^*[P_t]$. This is typically satisfied if $P_t^\ell, P_t^{\ell'}$ and $\alpha_{\ell,\ell'}$ are asymptotically independent for $\ell'$ chosen uniformly at random on $[L] \setminus \{\ell\}$. We will not discuss this assumption quantitatively, but we note that the Sherrington-Kirkpatrick from statistical physics tend to display phase transitions for low temperature (corresponding to small values of $\omega_e^2$) to a state in which this mean-field assumption is violated (this is known as Replica Symmetry Breaking) [40].
- We may replace $\Delta_t$ with $\Delta^*$ in the equation for $P_t$ through a mean-field approximation or time-averaging.

Under these three approximations, the equation of the typical locus $P_t$ is well approximated by

$$\mathrm{d}P_t = \xi_{\Delta^*,\hat{\alpha}^*}(P_t)P_t(1-P_t)\mathrm{d}t + (\theta^+(1-P_t) - \theta^- P_t)\mathrm{d}t + \sqrt{P_t(1-P_t)}\mathrm{d}B_t$$

where $\xi$ is defined as in (6). The corresponding stationary distribution of $P_t$ is

$$\Pi_{\hat{\alpha}^*,\Delta^*,\theta}(p) := Cp^{2\theta^+-1}(1-p)^{2\theta^--1}\exp\left[2\int_0^p \xi_{\Delta^*,\hat{\alpha}^*}(p')\mathrm{d}p'\right].$$

Finally, from the definition of $\hat{\alpha}$ in (89), the definition of $\Delta_t$, and the second approximation the fixed point system for $(\alpha^*, \Delta^*)$ is

$$\Delta^* = 2\mathbb{E}\left[\alpha\right]\mathbb{E}\left[L\int p\Pi_{\hat{\alpha}^*,\Delta^*,\theta}(p)\mathrm{d}p\right]^2 - \eta$$
$$\hat{\alpha}^* = 2\mathbb{E}\left[\alpha\right]\mathbb{E}\left[L\int p\Pi_{\hat{\alpha}^*,\Delta^*,\theta}(p)\mathrm{d}p\right]$$

where in both equations the second expectation is over $\theta$. We rewrite this as

$$\Delta^* = \frac{(\hat{\alpha}^*)^2}{2\mathbb{E}\left[\alpha\right]} - \eta \tag{90}$$

$$\hat{\alpha}^* = 2\mathbb{E}\left[\alpha\right]\mathbb{E}\left[L\int p\Pi_{\hat{\alpha}^*,\Delta^*,\theta}(p)\mathrm{d}p\right]. \tag{91}$$

It only remains to discuss the first assumption used. We claim

**Claim 12.4.** *Assume* $L|\bar{\theta}|\mathbb{E}\left[\alpha^2\right] + L^2|\bar{\theta}|\mathbb{E}\left[\alpha\right]^2 \ll \omega_e^2$*. Then* $|\epsilon_\ell(t)| \ll \omega_e^2$*.*

*Derivation of claim 12.4.* We start by noticing

$$\mathbf{Cov_p}\left[\left(Z(g) - \mathbf{E_p}\left[Z(g)\;|g_\ell\right]\right)^2, g_\ell\right] = \mathbf{Cov_p}\left[\mathbf{Var_p}\left[Z(g)\;|g_\ell\right], g_\ell\right]$$

Using the independence of the $(g_{\ell'})_{\ell' \in [L]}$, we write for $\mathbf{p} \in [0,1]^L$

$$\mathbf{Var_p}[Z(g)|g_\ell] = g_\ell^2 \sum_{\ell' \neq \ell} \mathbf{Var_p}[g_{\ell'}]\alpha_{\ell,\ell'}^2 \quad + \quad 2g_\ell \sum_{\substack{\ell',\ell'' \in [L] \setminus \{\ell\} \\ \ell' \neq \ell''}} \mathbf{Cov_p}[g_{\ell'}, g_{\ell'}g_{\ell''}]\alpha_{\ell,\ell'}\alpha_{\ell',\ell''} \quad + \quad R$$

where $R$ is independent of $g_\ell$. This yields

$$\mathbf{Var_p}[Z(g)|g_\ell] = g_\ell^2 \sum_{\ell' \neq \ell} 2p_{\ell'}(1-p_{\ell'})\alpha_{\ell,\ell'}^2 \quad + \quad 2g_\ell \sum_{\substack{\ell',\ell'' \in [L] \setminus \{\ell\} \\ \ell' \neq \ell''}} 2p_{\ell'}(1-p_{\ell'})2p_{\ell''}\alpha_{\ell,\ell'}\alpha_{\ell',\ell''} \quad + \quad R$$

From there it can be shown

$$\begin{aligned}\mathbf{Cov_p}\left[\left(Z(g) - \mathbf{E_p}\left[Z(g)\;|g_\ell\right]\right)^2, g_\ell\right] = 2p_\ell(1-p_\ell)(1-2p_\ell) &\sum_{\ell' \in [L] \setminus \{\ell\}} 2p_{\ell'}(1-p_{\ell'})\alpha_{\ell,\ell'}^2 \\ + 4p_\ell(1-p_\ell) &\sum_{\substack{\ell',\ell'' \in [L] \setminus \{\ell\} \\ \ell' \neq \ell''}} 2p_{\ell'}(1-p_{\ell'})2p_{\ell''}\alpha_{\ell,\ell'}\alpha_{\ell',\ell''}.\end{aligned}$$

We thus obtain

$$\epsilon_\ell(t) = (1-2P_t^\ell) \sum_{\ell' \in [L] \setminus \{\ell\}} 2P_t^{\ell'}(1-P_t^{\ell'})\alpha_{\ell,\ell'}^2 + 2 \sum_{\substack{\ell',\ell'' \in [L] \setminus \{\ell\} \\ \ell' \neq \ell''}} 2P_t^{\ell'}(1-P_t^{\ell'})2P_t^{\ell''}\alpha_{\ell,\ell'}\alpha_{\ell',\ell''}.$$

A mean-field approximation, considering $\alpha_{\ell',\ell''}, P_t^{\ell'}, P_t^{\ell''}$ to be close to independent yields

$$|\epsilon_\ell(t)| \sim L\mathbb{E}\left[\alpha^2\right]\mathbb{E}^*[P_t(1-P_t)] + L^2\mathbb{E}^*[P_t(1-P_t)]\mathbb{E}\left[\alpha\right]^2$$

By analogy with Section 5, we may consider $\mathbb{E}^*[P_t(1-P_t)] \sim |\bar{\theta}|$ and therefore

$$|\epsilon_\ell(t)| \sim L|\bar{\theta}|\mathbb{E}\left[\alpha^2\right] + L^2|\bar{\theta}|\mathbb{E}\left[\alpha\right]^2.$$

This yields the result. □

##### 12.3.2 The general formulation

We now take $Z$ to be some unspecified function on $\{0,1,2\}^L$. We start by decomposing the selection coefficient at locus $\ell$ into an additive, dominant and an epistatic component. We can find an approximate stationary distribution for the typical locus and the corresponding fixed point equation for $\Delta^*$ when the epistatic component can be neglected and the additive and dominant components are close to constant. Both these requirements will be stated as quantitative assumptions on population observables.

**Decomposition of the selection coefficient.** Let us define the following coefficients

$$\hat{\alpha}_\ell(t) := \frac{1}{2}\left(\mathbf{E}_{\mathbf{P}_t}[Z(g)\;|g_\ell = 2] - \mathbf{E}_{\mathbf{P}_t}[Z(g)\;|g_\ell = 0]\right) \tag{92}$$

$$\hat{D}_\ell(t) := \frac{1}{2}\left(\mathbf{E}_{\mathbf{P}_t}[Z(g)\;|g_\ell = 2] + \mathbf{E}_{\mathbf{P}_t}[Z(g)\;|g_\ell = 0] - 2\mathbf{E}_{\mathbf{P}_t}[Z(g)\;|g_\ell = 1]\right) \tag{93}$$

$$\hat{\beta}_\ell(t) := \hat{\alpha}_\ell(t) + \hat{D}_\ell(t)(1-2P_t^\ell) \tag{94}$$

$$\epsilon_\ell(t) := \mathbf{Cov}_{\mathbf{P}_t}\left[\mathbf{Var}_{\mathbf{P}_t}\left[Z(g)\;|g_\ell\right], g_\ell\right]/\mathbf{Var}_{\mathbf{P}_t}[g_\ell] \tag{95}$$

$$\Delta_t := \mathbf{E}_{\mathbf{P}_t}[Z(g)] - \eta. \tag{96}$$

We call $\hat{\alpha}_\ell(t)$ and $\hat{D}_\ell(t)$ the effective additive and dominance coefficients at locus $\ell$ at time $t$, $\hat{\beta}_\ell(t)$ the effective average effect at locus $\ell$ at time $t$, $\epsilon_\ell(t)$ the effective contribution of locus $\ell$ to epistasis variance. We derive the following decomposition

**Claim 12.5.** *We have*

$$s_\ell(\mathbf{P}_t) = s_\ell^{Dom}(t) - \frac{\epsilon_\ell(t)}{2\omega_e^2} \tag{97}$$

*where, by analogy with* (71) *we define*

$$s_\ell^{Dom}(t) := \omega_e^{-2}\left(\hat\beta_\ell(t)^2\left(P_t^\ell - \frac{1}{2}\right) + 2\hat\alpha_\ell(t)\hat D_\ell(t)P_t^\ell(1-P_t^\ell) - \hat\beta_\ell(t)\Delta_t\right).$$

**Remark 4.** *If we neglect $\epsilon_\ell(t)$, then we recover the same coefficient as for dominance (Section 11.3), with the difficulty that $\hat\alpha_\ell$ and $\hat D_\ell$ are functions of time. Looking at Eq. (92-95), we see neglecting $\epsilon_\ell(t)$ is acceptable if $g_\ell$ has more impact on the mean trait through $\mathbf{E_p}[Z(g)|g_\ell]$ than on the variance of the trait through $\mathbf{Var_p}[Z(g)|g_\ell]$.*

*Derivation of Claim 12.5.* We write the following decomposition as in (88)

$$\mathbf{Cov_p}\left[(Z(g)-\eta)^2, g_\ell\right] = \mathbf{Cov_p}\left[\left(Z(g) - \mathbf{E_p}\left[Z(g)|g_\ell\right]\right)^2, g_\ell\right] + \mathbf{Cov_p}\left[\left(\mathbf{E_p}\left[Z(g)|g_\ell\right]-\eta\right)^2, g_\ell\right] \tag{98}$$

The first term is $\epsilon_\ell(t)$. We thus obtain from (81)

$$\begin{aligned} s_\ell(\mathbf{P}_t) &= -\frac{\epsilon_\ell(t)}{2\omega_e^2} - \frac{\mathbf{Cov_p}\left[\left(\mathbf{E_p}\left[Z(g)|g_\ell\right]-\eta\right)^2, g_\ell\right]}{2\omega_e^2\mathbf{Var_p}[g_\ell]} \\ &= -\frac{\epsilon_\ell(t)}{2\omega_e^2} + \frac{\mathbf{Cov_p}\left[W\left(\mathbf{E_p}\left[Z(g)|g_\ell\right]\right), g_\ell\right]}{\mathbf{Var_p}[g_\ell]} \end{aligned}$$

Note that with the notation Eq. (92-93), we have

$$\mathbf{E_p}\left[Z(g)|g_\ell\right] = \mathbf{E_p}\left[Z(g)|g_\ell = 0\right] + g_\ell\hat\alpha_\ell(t) + \mathbb{1}_{[g_\ell=1]}\hat D_\ell(t) \tag{99}$$

From there, the same computations as in Section 11.3 yield

$$\mathbf{Cov_p}\left[W\left(\mathbf{E_p}\left[Z(g)|g_\ell\right]\right), g_\ell\right] = s_\ell^{Dom}(t)P_t^\ell(1-P_t^\ell).$$

□

**Stationary solution and fixed-point equation** As always, the typical locus $(P_t, \hat\alpha(t), \hat D(t), \theta)$ is defined as $(P_t^\ell, \hat\alpha_\ell(t), \hat D_\ell(t), \theta_\ell)$ where $\ell$ is picked uniformly at random. In the general model, it is not obvious that the marginal over a typical locus of the stationary solution (83) converges to a unique well-defined distribution as $L \to +\infty$. Nevertheless, we can find the stationary distribution when the three following conditions are satisfied

- We may neglect the epistasis selection term: $|\epsilon(t)| \ll \omega_e^2$
- The effective additive and dominance terms $\hat\alpha(t)$ and $\hat D(t)$ remain effectively constant, equal to their expected value $\hat\alpha^* := \mathbb{E}^*[\hat\alpha(t)]$ and $\hat D^* := \mathbb{E}^*[\hat D(t)]$.
- We may replace (through a mean-field approximation or time-averaging) $\Delta_t$ with $\Delta^*$ in the equation for $(P_t)_{t\geq 0}$.

Under these approximations, we may write the dynamics of $(P_t)_{t\geq 0}$ as

$$\mathrm{d}P_t = \xi_{\Delta^*,\hat\alpha^*,\hat D^*}(P_t)P_t(1-P_t)\,\mathrm{d}t + \left(\theta^+(1-P_t) - \theta^- P_t\right)\mathrm{d}t + \sqrt{P_t(1-P_t)}\,\mathrm{d}B_t^P$$

where $\xi_{\delta,\alpha,D}$ is defined as in (71).

In particular, the stationary distribution of $P_t$ conditional on $(\hat\alpha^*, \hat D^*, \theta)$ is $\Pi_{\Delta^*,\hat\alpha^*,\hat D^*,\theta}$ where $\Pi_{\delta,a,d,\theta}$ is defined as in (73).

From there, we may as above find a fixed-point system for $(\Delta^*, \hat\alpha^*, \hat D^*)$ by expressing them as functions of the law of a certain number of independent typical loci (in the example $Z_{sg}$ above, $\Delta^*$ was expressed as a function of two typical loci, whereas $\hat\alpha^*$ was a function of one typical locus).

We expect this fixed-point system to be accurate when $Z$ is a polynomial of $g$ of low order $k$ (for instance order 2 in (79)) in which case the fixed-point system can be further simplified (replacing the integral over $[0,1]^L$ with one over $[0,1]^k$ corresponding to a "typical" $k-$tuple of loci). If $Z$ instead is a polynomial of high order, for instance

$$Z(g) = \prod_{\ell\in[L]}(g_\ell - 1)$$

then we cannot expect the fixed-point system to be accurate.

### 13 Numerical methods to solve the fixed-point equation

Solving the fixed-point equation (14) requires the computation of integrals against $\Pi_{\delta,a,\theta}$ defined in (13). In particular we must compute integrals of the form

$$\mathcal{I}_{a,b,c,d} := \frac{1}{Beta(a,b)} \int_0^1 x^{a-1}(1-x)^{b-1} e^{cx - dx(1-x)} \mathrm{d}x$$

with $a, b > 0$ possibly small, $c \in \mathbb{R}, d \in \mathbb{R}_+$, and $Beta$ is the beta function (added here for mathematical convenience).

We use two methods to compute this integral. The first method is used to obtain theoretical predictions in every figure except Figure 4 (that is, Figures 3, 5, S4-6)

$$\begin{aligned}\mathcal{I}_{a,b,c,d} &\simeq \sum_{k\in[k_{max}]} \frac{d^k}{k!} \int_0^1 x^{a+k-1}(1-x)^{b+k-1} e^{cx} \mathrm{d}x \\ &= \sum_{k\in[k_{max}]} \frac{d^k}{k!} \frac{(a)_k (b)_k}{(a+b)_{2k}} {}_1F_1(a+k; a+b+2k; c)\end{aligned}$$

where $k_{max}$ is as large as computationally possible, ${}_1F_1$ is the confluent hypergeometric function, and $(a)_k := a(a+1)\ldots(a+k-1)$ is the rising factorial, with $(a)_0 = 1$. This method fails when $d > 1$, which in our setting corresponds to $\omega_e < \alpha$.

The second method is the one we used in Figure 3 in the main text and assumes low mutation rates ($|\bar{\theta}| \ll 1$), following SI 8. In this figure, we solve for $\Delta^*$ by writing $\xi_{\delta,a} \simeq -\frac{\delta a}{\omega_e^2}$ in the definition of $\Pi$ in (13), neglecting Robertson's underdominant term. This is justified because we expect allele frequencies to be very close to $\{0,1\}$ under low mutation rates.

With this approximation, we may compute as in Section 5.3

$$\int p \Pi_{\delta,a,\theta}(p) \mathrm{d}p \simeq \frac{\theta^+}{|\theta|} \times \frac{{}_1F_1(2\theta^+ + 1 \ ; \ 2|\theta| + 1 \ ; \ 2s^* La)}{{}_1F_1(2\theta^+ \ ; \ 2|\theta| \ ; \ 2s^* La)}$$

which we use in (14) to compute $\Delta^*$. To compute the genetic variance, we write

$$\begin{aligned}\int p(1-p)\Pi_{\Delta^*,a,\theta}(p)\mathrm{d}p &= \frac{\int p^{2\theta^+}(1-p)^{2\theta^-} e^{2s^* Lap - 2\frac{a^2}{\omega_e^2} p(1-p)} \mathrm{d}p}{\int p^{2\theta^+ - 1}(1-p)^{2\theta^- - 1} e^{2s^* Lap - 2\frac{a^2}{\omega_e^2} p(1-p)} \mathrm{d}p} \\ &\simeq \frac{\int e^{2s^* Lap - 2\frac{a^2}{\omega_e^2} p(1-p)} \mathrm{d}p}{\int p^{2\theta^+ - 1}(1-p)^{2\theta^- - 1} e^{2s^* Lap} \mathrm{d}p}\end{aligned}$$

where in the numerator, we wrote $p^{2\theta^+}(1-p)^{2\theta^-} \simeq 1$ (using $|\bar{\theta}| \ll 1$) and in the denominator, we wrote $p(1-p) \simeq 0$ using that allele frequencies are close to $\{0,1\}$ with high probability. This can be rewritten

$$\int p(1-p)\Pi_{\Delta^*,a,\theta}(p)\mathrm{d}p \simeq Beta(2\theta^+, 2\theta^-) \frac{\int e^{2s^* Lap - 2\frac{a^2}{\omega_e^2} p(1-p)} \mathrm{d}p}{{}_1F_1(2\theta^+ \ ; \ 2|\theta| \ ; \ 2s^* La)}$$

which can be computed efficiently using the scipy package.

Future work could look for more precise and effective ways to implement these computations.
